# Discovery of Natural Bispecific Antibodies: Is Psoriasis Induced by a Toxigenic *Corynebacterium simulans* and Maintained by CIDAMPs as Autoantigens?

**DOI:** 10.1101/2023.09.04.556232

**Authors:** Jens-Michael Schröder

**Affiliations:** University-Hospital Schleswig-Holstein, Campus Kiel Dpt. of Dermatology, Kiel, Germany

**Keywords:** psoriasis, autoantigen, bispecific antibody, Corynebacterium simulans, phage, disordered protein, epitope, Cationic Intrinsically Disordered Antimicrobial Peptides

## Abstract

The high abundance of *Corynebacterium simulans* in psoriasis skin suggests a contribution to the psoriasis etiology via cell envelope components, which may cause skin inflammation and immune responses. This hypothesis was tested in an exploratory study, where Western Blot (WB) analyses with extracts of heat-treated *C. simulans* and psoriasis serum-derived IgG exhibited a single 16 kDa- WB-band. Proteomic analyses revealed, among others, ribosomal proteins as candidate *C. s.*-antigens. A peptidomic analysis unexpectedly showed that psoriasis-serum-derived IgG already contained 31 immunopeptides originating from *Corynebacteria ssp*., suggesting the presence of natural bispecific antibodies (BsAbs). Moreover, peptidomic analyses revealed 372 “DECOY”-peptides with similarity to virus- and phage proteins, including *Corynebacterium diphtheriae phage*, and similarity to diphtheria toxin. Strikingly, upon a peptidomic analysis for peptides of human origin, 64 epitopes of major psoriasis autoantigens were identified, which originated from the spacer region of filaggrin, from hornerin repeats, SPINK9, keratin 9, caspase 14, desmoplakin, suprabasin, keratin 2, keratin1, keratin 6C, apolipoprotein A1, a Selene-binding protein, H1.8 linker histone, and the transcription factor BCLAF3. Most identified antigens represent potential “Cationic Intrinsically Disordered Antimicrobial Peptides (CIDAMPs)”, which are generated within the fully differentiated epidermis. These may form complexes with bacterial disordered protein regions, representing chimeric antigens containing discontinuous epitopes. In addition, among 128 low-abundance immunopeptides, 48 are putatively psoriasis-relevant such as epitopes of IL-12, and the receptors of PGE2, vitamin D3, and IL-10. Further, 47 immunopeptides originated from tumor antigens such as CT47A, SDCCAG3, BRCA2, MAGEA6, RNASE4, and the endogenous retrovirus HERV-K. I propose that persistent infection with a toxigenic *C. simulans* initiates psoriasis, which is exacerbated as an autoimmune disease by CIDAMPs as autoantigens. The discovery of natural BsAbs allows the identification of antigen epitopes from microbes, viruses, autoantigens, and tumor-antigens, and may help to develop epitope-specific peptide-vaccines and therapeutic approaches with antigen-specific regulatory T cells to improve immune tolerance in an autoimmune disease-specific-manner.

## Introductionary Overview

Psoriasis is a common chronic inflammatory human skin disease, existing on a fairly broad spectrum in terms of clinical manifestations. The exact etiological and pathogenic mechanisms are still not known in detail. Current knowledge indicates an overlap with other inflammatory as well as autoimmune disorders. Clinical observations of psoriasis can be interpreted as consequences of the activation of different arms of the immune system: Whereas responses by the adaptive immune system dominate in chronic plaque psoriasis, innate immune reactions, and autoinflammatory responses are dominant in guttate psoriasis and generalized pustular psoriasis^1^. It is still a matter of current debate if psoriasis is a primary autoimmune disease or secondarily evolving into autoimmunity as seen in other chronic inflammatory diseases^2^. Serological autoimmune phenomena, namely diverse circulating specific autoantibodies to the stratum corneum identified in psoriasis patients^3.4,5^ support the idea of an autoimmune disease. Nevertheless, autoreactive CD8^+^T-cells, located within the lesional epidermis^6^ remain among the most important cellular players in the pathogenesis of psoriasis and current therapeutic strategies and clinical and experimental evidence suggest that both, autoimmune and autoinflammatory reactions are central to psoriasis^1^. Indeed, several studies have attempted to address this and several putative autoantigens in psoriasis have been proposed. One of these is keratin 17. The involvement was proposed because of keratin 17 similarity to streptococcal M-proteins^7^. Others are complexes of self-DNA with the host defense peptide cathelicidin (LL37)^8^ or melanocytic ADAMTSL5^9^, which may act as auto-antigens in some cases of psoriasis.

Infections can provoke or exacerbate psoriasis^10^. There is strong evidence that guttate psoriasis is preceded by tonsillar *Streptococcus pyogenes* infection^11^ and disease exacerbation has been linked with skin colonization by *Staphylococcus aureus*, *Malassezia spp.*, and *Candida albicans*^11^. A role, if any, of viruses present in lesional skin is currently speculative^12^.

There is an ongoing discussion that commensal microbes could cause autoimmunity. Numerous scientific studies in recent years have shown significant skin dysbiosis with a decrease in microbiome diversity in psoriasis skin. *Corynebacterium simulans*, *Corynebacterium kroppenstedtii*, *Finegoldia* spp., and *Neisseria* spp. were found to be increased in the psoriasis skin and *Burkholderia* spp., *Cutibacterium*, and *Lactobacilli* were decreased when compared with healthy skin^13,14,15,16,17,18,19,20^. Whether any of the pathogens found to be highly abundant in lesional psoriatic skin is a trigger factor in psoriasis is yet unknown.

### Corynebacterium simulans and C. kroppenstedtii: Putative Psoriasis-trigger?

Corynebacteria, which are dominant members of skin microbiota, were recently shown to promote, independently of other microbes, in an IL-23-dependent manner, a dramatic increase in the number and activation of a subset of γδ T cells in a mouse model^21^. This effect is highly conserved among species of *Corynebacterium* and dependent on the expression of a dominant component of the cell envelope, mycolic acid. Thus, it is tempting to speculate that *C. simulans sp. nov.,* a recently discovered and skin, but not mucosal surfaces colonizing germ^22^, and/or *C. kroppenstedtii* may have a role in psoriasis etiology. The *Corynebacterium* is a genus of bacteria of growing clinical importance, in particular due to an increasing number of infections caused by these opportunistic pathogens^23^. In contrast to the lipophilic coryneform taxa, which represent the dominating *Corynebacterium* species of healthy skin and are causing opportunistic infections, *C. simulans* are Gram-positive, non-lipophilic, facultatively anaerobic and fermenting bacteria, showing a diphtheroid arrangement^22^. In contrast to lipophilic coryneform taxa, nonlipophilic coryneform taxa are rather rare members of the resident healthy skin flora^22^.

It is interesting to note that *C. simulans* contains a high percentage of meso-diaminopimelic acid (DAP) in the murein cell wall^22^, another characteristic shared with all *Corynebacteria*. DAPs in peptidoglycans (PGNs) are recognized by NOD1 receptors, whereas muramyl dipeptide structures in PGNs are recognized by NOD2. Both NOD1 and NOD2 receptors are intracellular receptors, which originally led to the suggestion that only intracellular bacteria are recognized. Strikingly, there are high levels of NOD1-stimulatory activity in culture supernatants of *Corynebacteria* as compared with that found in bacterial cell extracts^24^. The NOD1-stimulatory activity, unlike NOD2 and TLR4- stimulatory activities, is highly stable and cannot be removed by boiling or treatment at extreme pH conditions^24^. Thus NOD1-stimulatory activity should be present and persisting where *C. simulans* colonizes the skin. This would further implicate that NOD1-receptor actions could be trigger factors in psoriasis.

NOD1 activation in epithelial cells, including keratinocytes, leads to type I interferon (IFNß) signaling resulting in a characteristic chemokine profile (CXCL9-11), and NFĸB activation with induction of IL-1ß, IL-8, IL-18, CXCL1, CXCL9-11, hBD-2 and also S100A7, S100A8 und S100A9^25,26,27,28,29^ - all cytokines, which are highly expressed in psoriasis lesions. Administration of NOD1 ligands into mice induced chemokines and recruitment of acute inflammatory cells, an activity that was abolished in NOD1-null mice^29^. Microarray analysis revealed that NOD1 stimulation induces only a restricted number of genes in intestinal epithelial cells, suggesting that NOD1 functions as a pathogen recognition molecule to induce the expression of molecules involved in the early stages of the innate immune response^29^. NOD1 is of critical importance for the recruitment of neutrophils upon bacterial infection. NOD1(-/-) mice, that have been infected with *C. difficile* showed a reduced recruitment of neutrophils, which correlated with impaired production of the neutrophil attractant CXCL1, but not CCL2, XCL1, and other monocyte- or lymphocyte attracting/activating cytokines/chemokines^30^. Further results of this study suggested that NOD1 directly recognizes *C. difficile* to induce the recruitment of neutrophils to the infected site, indicating that NOD1 regulates host susceptibility to *C. difficile* and suggests that NOD1-mediated neutrophil recruitment is an important immune response against this pathogen^30^ and possibly also of relevance upon skin infection caused by *Corynebacteria*. Further, IL-36 cytokines, which are highly expressed in psoriasis^30,31^ and are powerful inducers of the antimicrobial peptide hBD2, which is present at excessive amounts in psoriatic scales^32^, are induced by pathogenic bacteria and are a key initiator of immune responses and pathological inflammation within epithelial tissues, being global discriminators of harmless microbes and invasive pathogens within epithelial tissue^33^.

### Are *C. simulans* Phages Virulence Factors in Psoriasis?

Using improved sequencing technology to identify additional microbial diversity and less abundant taxa of the skin microbiome, a recent study could reconstruct a complete genome sequence of *C. simulans*^34^. It has been shown that many bacterial species contain a significant pan-genome defined as the aggregate coding potential of sub-strains within that clade. Strain variation is an important consideration, as flexible regions of the genome can encode different properties of transmissibility, virulence, antibiotic resistance, or other functions. Strain-level populations of common skin commensals have been recently identified and are heterogeneous and comprised of multiple subspecies clades^35^ and were recently investigated for *C. simulans* populations of the foot. As a result, a total of 411 variants were identified, which occurred primarily in protein-coding regions, most of which were part of highly conserved protein complexes^34^. For example, 16 of the 55 genes encode ribosomal components. However, some of these variants may be derived from non-*Corynebacterium* genomes^34^. Another interesting finding was the discovery of a putative septicolysin toxin gene, which was found to have three variants, two of them nonsynonymous, appearing to be a novel protein sharing only 93% protein identity with the closest homologs belonging to other *Corynebacteria*^34^. Septicolysins are pore-forming toxins that are commonly associated with bacterial pathogenesis^36^.

As a further interesting result, an integrated *C. simulans* phage was identified, which was absent in another skin strain. This phage has well-defined attachment sites, complete head, and tail proteins, integrases, and a type VI secretion system^34^. This phage also contains lytic elements that encode chitinase-like proteins, which are similar to lysozyme-like families of hydrolases that drive virulence^34^. *Corynebacterium* phages are particularly poorly characterized with only two characterized genomes, neither of which occur in greater than trace amounts in larger skin populations^35^. Corynebacterium- derived phages are an important source of toxins. The best known are those producing diphtheria toxin (DT) from *Corynebacterium diphtheria*, causing diphtheria, also in the skin. Cutaneous diphtheria is normally associated with colonization of pre-existing skin lesions, such as wounds, by both DT-producing and non-DT-producing *<u>C. diphtheriae</u>* strains^37^. In *C. diphtheriae*, diphtheria toxin is encoded by the tox gene of some temperate corynephages such as beta. Beta-like corynephages are capable of inserting into the *C. diphtheriae* chromosome at specific sites. Transcription of the phage-encoded tox gene, and many chromosomally encoded genes, is regulated by the DtxR protein in response to Fe^2+^ levels. Characterizing DtxR-dependent gene regulation is pivotal in understanding diphtheria pathogenesis and mechanisms of iron-dependent gene expression, although this has been hampered by a lack of molecular genetic tools in *C. diphtheriae* and related coryneform species^38^. Thus, it would be interesting to speculate that *C. simulans* in psoriatic skin lesions bear phages, which may secrete lytic proteins, which may act as virulence factors, stimulating innate immunity reactions towards *C. simulans* and also the phages.

Direct interactions between phages and the human immune system are just beginning to be systematically investigated. Phages are abundant within the human body. Most of them are present at sites of bacterial colonization, including the skin^39^. Bacteriophages are an integral part of our relationships with bacteria and tri-kingdom interactions between bacteria, bacteriophages, and the human immune system play important roles in health and disease^40^. It has been proposed that phages may contribute to the stability and resilience of the microbiome^41^. Bacteriophages have indirect effects on mammalian immunity, where phage genetic elements serve as virulence factors allowing for bacterial colonization and invasion of their mammalian host. Bacterial expression of phage-encoded proteins increases epithelial adhesion, invasion, biofilm formation, and it inhibits neutrophil phagocytosis. Other phage-encoded proteins are involved in exotoxin production or delivery, cytotoxicity, intracellular infection, and superantigen production^40^.

Phages can be recognized by several host cell-surface and intracellular pattern-recognition receptors (PRRs). Pathways involving sensing of single-stranded DNA and double-stranded DNA and the induction of IFN responses are most commonly implicated^40^. Several studies identified a role of TLR9, an endosomal PRR expressed in pDC^42^ and keratinocytes^43^, in phage recognition^44^ TLR9 recognizes unmethylated CpG motifs abundant in the DNA of phages as well as the bacteria that produce them^45,46^. TLR9 signals through the adaptor protein MyD88 and induces the expression of proinflammatory and antiviral cytokines^45^. Recent research on phage sensing in the gut has also established a role for TLR9 in promoting inflammatory responses to phages. An oral cocktail of *E. coli*-tailed phages led to significantly increased IFN-γ-producing CD4+ T cells, driven by DC sensing of phage DNA through TLR9^45^.

Bacteriophages interact with cells of the adaptive immune system^40^. Phage-derived peptides are presented by APCs to naïve T cells on MHC-II molecules. Naïve CD4^+^ T cells specific for the antigen-MHC undergo activation with the release of IFN-γ and proliferation into Th cells. Phage-specific Th cells are capable of activating naïve phage-specific B cells which eventually result in phage-specific antibodies^40^. Thus, bacteriophages could have a role in psoriasis pathogenesis. In a recent study, phage and bacterial components of the skin microbiome in patients with psoriasis and healthy family controls have been investigated. Here, among phage species with abundant host bacteria, *Acinetobacter phage Presley* and *Pseudomonas phage O4* were found to be more abundant in psoriasis lesional skin than healthy controls^47^.

The cytokine signature seen in psoriasis lesions, at least in part, points towards a virus infection (or virus-infection imitating infection) of keratinocytes as a trigger factor in psoriasis. This hypothesis is based on the recent findings of Wolk et al^48^.

They showed that antiviral proteins (AVPs) such as MX1, BST2, ISG15, and OAS2 were strongly elevated in lesional psoriatic skin and they found further that interleukin-29 (IL-29) was the only mediator whose expression correlated with the AVP levels. Neutralization of IL-29 in psoriatic skin reduced AVP expression. Accordingly, IL-29 raised AVP levels in isolated keratinocytes, epidermis models, and human skin explants, but did not influence antibacterial protein production, and AVP induction correlated with increased antiviral defense of IL-29-treated keratinocytes^48^. It would be interesting to elucidate whether bacteriophages cause AVP-induction and induce IL-29 in keratinocytes.

All the findings mentioned above would favor the hypothesis that infection by phage-bearing *C. simulans*, expressing phage-derived toxin(s), could be a putative initial trigger factor in psoriasis.

### Psoriasis Serum Contains Bispecific Antibodies towards *C. simulans* and *C. kroppenstedtii* Antigens

If *C. simulans* (and possibly *C. kroppenstedtii*) is infecting skin in psoriasis patients, one would expect immune responses toward *C. simulans*. To test this hypothesis, in an exploratory study using Western Blot (WB) technology, pooled serum obtained from hospitalized psoriasis patients was tested for immunoreactivity towards *C. simulans* antigens. The presence of several immunoreactive bands, mainly towards 14 – 25 kDa proteins, but also towards proteins >70 kDa confirmed this hypothesis and prompted us to go into more detail: We purified IgG from pooled psoriasis serum using protein A affinity chromatography and repeated the WB experiment with the IgG fraction and *C. simulans* extracts. Interestingly, only extracts from *C. simulans* bacteria (strain DSM 44415), which were cultured at planktonic conditions and extracted after heat treatment (95°C), or after sonication, revealed a single 16 kDa band upon WB with psoriasis-derived IgG (Fig.1). Since heat- or ultrasound- treatment transfers complexes of intrinsically disordered proteins (IDPs) or IDP-regions (IDPRs) into free IDPs or IDPRs^49^, the *C. simulans* antigen(s) seems to be an IDP/IDPR. To get information about *C. simulans* antigens being recognized by psoriasis-IgG, the 16 kDa-WB-band as well as a the corresponding area of an SDS-PAGE analysis of a freshly generated *C. simulans* extract was cut out and analyzed by proteomic analyses of trypsin-digested proteins by LC-MS/MS. Using Scaffold Software^TM^ (version 5.2.1), a bioinformatic tool for validating MS/MS-based proteomic data^53^, numerous non-human proteins were identified in the Universal Protein Resource (UniProt) databank, which fitted well with proteins of *Corynebacterium simulans* or *C. kroppenstedtii*.

**Figure 1:**
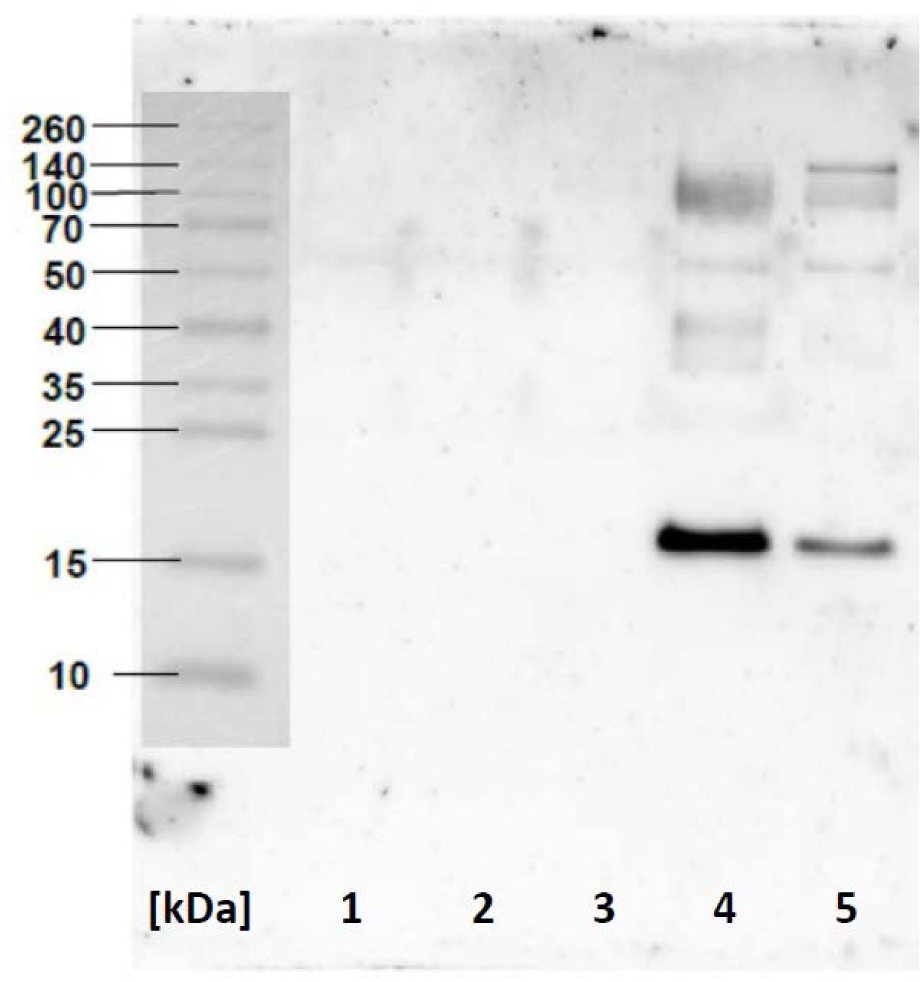
Western Blot Analysis of *Corynebacterium simulans* Extract with Psoriasis IgG. *C. simulans* (strain DSM 44415), grown at static or planktonic conditions, was lysed either by heating to 95 °C or sonication, and the extracts were applied in DTT-containing loading buffer and then separated on a 12% acrylamide:bisacrylamide gel as decribed^50^. Lane MW: Spectra^TM^ Protein Ladder; Lane 1: *C.s.*, static culture, heat treatment followed by immediate application; Lane 2: *C.s.*, static culture, heat treatment followed by storage in the deep freezer; Lane 3: *C.s*., planctonic culture, treated with bacterial lysis-buffer, followed by storage in the deep freezer; Lane 4: *C.s*., planctonic culture, heat treatment and immediate application; Lane 5: *C.s*., planctonic culture, sonication and immediate application. Western Blot (WB)-analyses were performed with an IgG preparation (isolated from a nonspecified pool of psoriasis-serum using standard methods^51^ (1mg/ml). Antigen-bound psoriasis-IgG was detected after treatment of the membrane with goat-anti-human IgG, peroxidase-coupled, and development with Roche LumiLight as described^50^. Note the presence of a WB-band only when the samples were heated or sonicated and immediately used, conditions known to dissociate complexes of Intrinsically Disordered Proteins and Regions (IDP(R)s)^52^.

Protein analysis of a SDS-PAGE slice of a heat-treated *C. simulans* extract, corresponding to the 16 kDa WB-band, apart from poorly or uncharacterized proteins, an ABC-type transport system, substrate-binding protein (44 kDa), the chaperone protein DnaK (66 kDa), thiamine-phosphate synthase (22 kDa) and the ribosomal proteins 30S S17 (10,6 kDa), 30S S18 (12,2 kDa), 50S L29 (8,5 kDa) and 50S L7/L12 (13,6 kDa) were identified. It is currently speculative, which of these proteins represents the *C. simulans* antigen(s), although the 50S ribosomal protein L7/L12 and some other ribosomal proteins would fit best with its MW.

To my surprise, proteomic analyses of the 16 kDa-band of the WB experiment revealed a very low abundance of the proteins, but unexpectedly, a very high number of individual, from different *C. simulans* proteins originating peptides, which were present in relatively high amounts, as revealed by a peptidomic analysis (Tab.1). These peptides did not originate from the *C. simulans* proteins separated on the gel, suggesting that these peptides represent obvious epitopes of *C. simulans* antigens, which were already bound to the paratopes of psoriasis-IgG. A Scaffold®Peptide Report of the WB- experiments (Tab.1) showed the presence of 31, mainly 7-mer to 12-mer peptides. The relative amounts of these peptides, given as “TICs” (Total Ion Chromatograms), revealed some quantitatively dominating immunopeptides such as a single, exclusive peptide (VADRSGGYTR) of 50S ribosomal protein L17 *C. kroppenstedtii*. In addition, peptides from several other ribosomal proteins were also seen, which includes 30S ribosomal proteins (S3, S4, S5, S7, S9, S13, and S16) and additional 50S ribosomal proteins (L3, L4, L5, L6, and L28) with sequence identity reported for *C. simulans* (Tab.1). Further, the by far most abundant immunopeptides came from a putative iron-dependent peroxidase and an ABC-type biotin transport system and permease protein of *C. kroppenstedtii* (Tab.1).

**Table 1:**
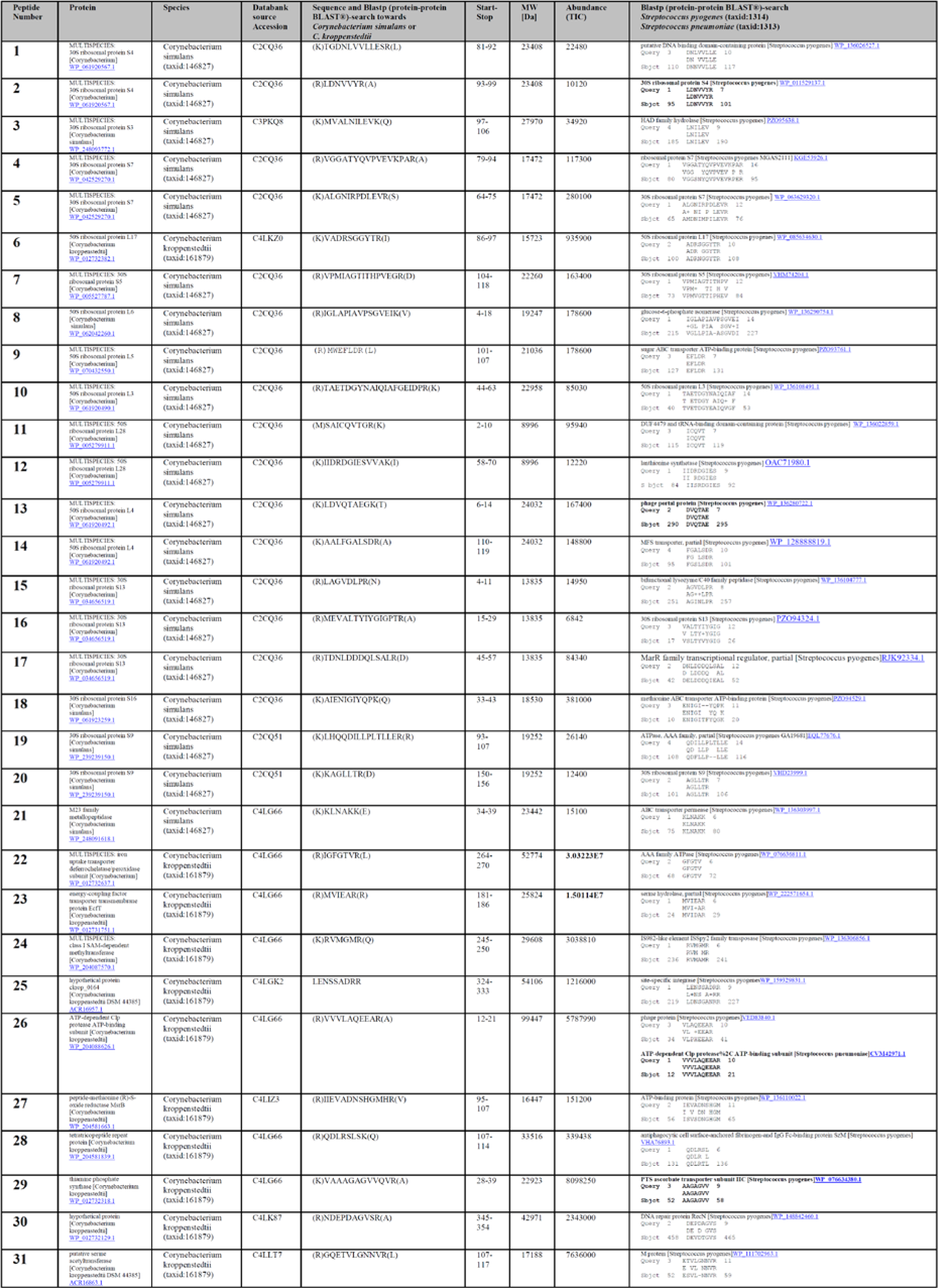
*C. simulans*-derived peptides identified within the immunoreactive 16 kDa *C. simulans*-WB-Band. The 16 kDa- WB-band of a freshly generated, heat-treated *C. simulans* extract was cut out and analyzed by proteomic analyses of on-membrane trypsin-digested proteins by LC-MS/MS. Analyzes were performed using Scaffold Software^TM^ (version 5.2.1). In the Universal Protein Resource (UniProt) databank *Corynebacterium simulans* and *C. kroppenstedtii* proteins are shown as Peptide Report. The relative abundance of the peptides is given as “TICs” (Total Ion Chromatograms). Alignments for the peptides are shown for BLASTP® (protein-protein BLAST®)- search towards *Corynebacterium simulans* (taxid:146827), *C. kroppenstedtii* (taxid:161879), *Streptococcus pyogenes* (taxid:1314) and *Streptococcus pneumoniae* (taxid:1313). Note the very high similarity or identity of several immunopeptides with peptide sequences of *S. pyogenes*.

These data support the hypothesis that IgG-antibodies obtained from psoriasis-serum are bispecific. Bispecific antibodies (BsAbs) are antibodies with two binding sites directed at two different antigens or two different epitopes on the same antigen^54^. Natural BsAbs have been identified against cyclic citrullinated peptide (CCP) and immunoglobulin G (IgG) in rheumatoid arthritis patients’ serum^55^ as well as in some patients with autoimmune thyroid disease, where BsAbs bind to thyroglobulin as well as thyroid peroxidase^56^. The epitopes, however, have not been identified.

The Scaffold®Peptide Report from proteomic analyses indicates antibody generation from chimeric antigens. I would propose that one antigen section originates from a yet not-identified 16 kDa- immunoreactive *C. simulans* protein and the other antigen section from a different *C. simulans* or *C. kroppen*stedtii proteins, yet identified upon proteomics analyses (Tab.1).

How could a chimeric peptide-antigen be generated? I propose as a source a complex of two assembling IDPs or IDPRs^57,58,59^. Such complexes are formed upon treatment of *E. coli* with antimicrobial peptide fragments of the IDP hornerin (HRNR). HRNR-peptides bind to ribosomal proteins, a likely mode of action of bactericidal activity of *Cationic Intrinsically Disordered Antimicrobial Peptides, CIDAMPs*^60^. IDPs can form hundreds of different protein complexes with other IDPs, which are rather stable at physiological conditions but can be dissociated upon heat treatment^59,52^, conditions necessary for antibody-binding to *C. simulans* antigens as shown in Fig.1, or to see CIDAMP-binding to ribosomal proteins^50^. So, I would propose that in the quaternary structure of a multimeric assembly complex of two IDP(R)s short, exposed peptide chains of each IDP form discontinuous epitopes from chimeric antigens, which both together will lead to the formation of BsAbs.

In summary, these observations support the hypothesis, that psoriasis serum contains BsAbs, which recognize discontinuous and chimeric epitopes of IDP(R)-complexes.

### Do Psoriasis Antibodies Recognize Antigens of *C. simulans* Phages?

Peptidomic analyses of on-membrane trypsin-digested proteins, present in the 16 kDa WB-band of a *C. simulans* extract, revealed, apart from the *C. simulans* proteins, several peptides declared as so-called “DECOY”- peptides. Sequences of DECOY-peptides are from a DECOY database but can match identically against a true peptide sequence (i.e., palindromic peptide sequences) in the original target database and thus may increase the false hit rate in the DECOY database and cause false discovery rate overestimation^61^. Alternatively, peptide sequences may be true because they have not been annotated, coming from yet not sequenced genomes, e.g. from the tens of thousands of viruses^62^. The finding that the human skin microbiome may contain *C. simulans* phages prompted me to suggest that these DECOY-peptides may originate from yet not annotated phage proteins. A BLAST® search (limited to viruses (taxid:10239)) of all 372 DECOY-peptides obtained from the 16 kDa WB-band of *C. simulans* (strain DSM 44415) extracts (Tab.S1) revealed indeed similarity to phage or virus proteins, supporting the hypothesis that in psoriasis lesions *C. simulans* phages had induced immune responses: Apart from many hypothetical proteins of unknown function, the majority of peptides showed similarity with the virus- or phage major and minor capsid proteins, tail fiber proteins, tail tape measure proteins, integrases, replicases, helicases, and viral ankyrin (Tab.S1). In addition, some peptides were similar to portal proteins, DNA primase and helicase, DNA polymerase, terminase family proteins as well as repressor and antirepressor proteins. Interestingly, several peptides revealed high similarity with proteins of influenza A and B viruses, SARS-COV-2, HIV-1, rotavirus A, molluscum contagiosum virus, hepatitis B and C virus, diverse human herpesvirus, measles morbillivirus, several human papillomavirus, human T-cell leukemia virus, norovirus, Dengue virus, monkeypox virus, vaccinia virus, human parainfluenza 3 virus, HIV-2, infectious bronchitis virus, infectious pancreatic necrosis virus, and cowpox virus (Tab.S1). Moreover, the existence of immunopeptides with similarity to phage-derived endolysins, lysins, toxins, lysozymes, lytic transglycosylase, holin family proteins, ADP-ribosyl transferases, diphtheria toxin, as well as very high similarity to proteins of *C. diphtheriae* (Tab.S1) may point towards immune reactions towards products of lytic and/or lysogenic *C. simulans* phages and prophages in psoriasis. Phages often maintain one of two life cycles: lytic or lysogenic. Lytic phages are free-living until they encounter their bacterial host, and produce virions that resultantly lyse the cell to release new phages ^63,64^. Lysogenic phages integrate their genomes into the chromosomes of the bacterial host cell as prophages. Here they lie dormant until triggered by some environmental factors to excise and enter the lytic circle^63^. The presence of 183 DECOY-peptides in the 16 kDa-SDS-PAGE-section of a *C. simulans* DSM 44415-extract (Tab.S2) suggests also that this strain contains a lysogenic phage. BLASTP® search for its similarity with proteins of diphtheria-causing corynephage beta (taxid 10703) revealed that most of the peptides showed similarity to diphtheria toxin (DT), DT-variants and DT- homologs (Tab. S2). When for all DECOY-peptide sequences showing similarity with the catalytic domain of diphtheria toxin (Chain A, DIPHTHERIA TOXIN [Corynephage beta] 1DDT_A), the corresponding DT sequence was aligned with the full-length DT sequence, 30 of all DECOY-peptides found in this study, are similar to DT (Fig.2). Thus, also a putative 16 kDa DT-like protein **(“Psoriasis Toxin”**) of a *Corynephage* may cause the immunoreactive WB-band (Fig.1).

**Figure 2:**
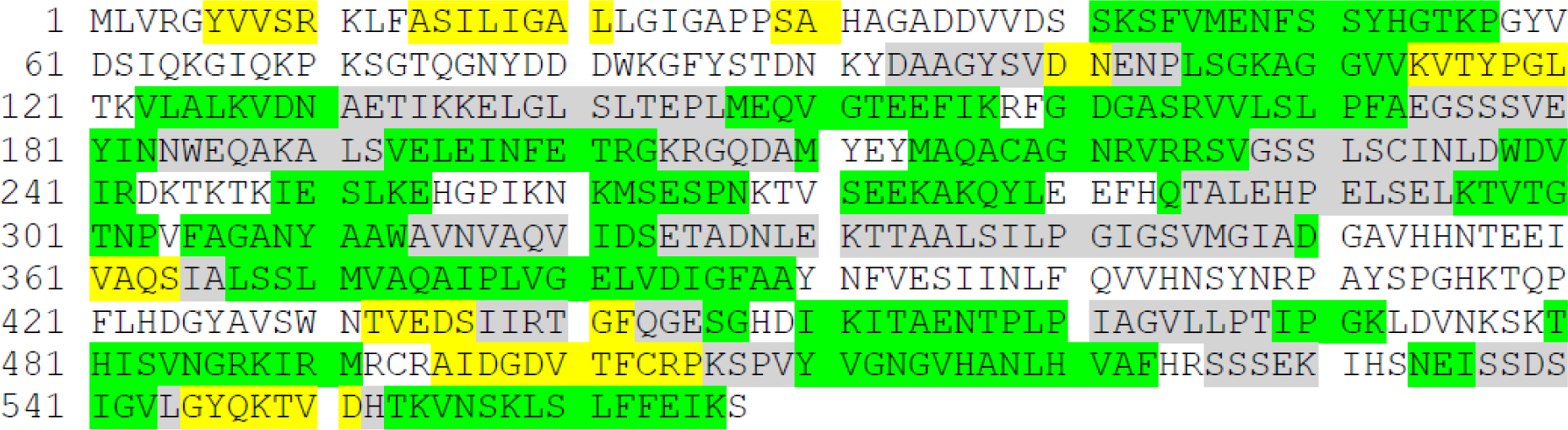
DECOY-peptides contain AA-Sequence-Motifs with Similarity to Diphtheria Toxin. The amino acid (AA) sequence of Diphtheria toxin (DT)/NAD(+)-diphthamide ADP-ribosyltransferase, with the sequence ID: <u>P00588.2</u>, is shown. BLASTP®- analyses of this study’s DECOY-peptides (Supplementary Tables 1 and 2) revealed AA-sequence motifs with similarity to DT. The corresponding DT-sequence motifs of DECOY-peptides from *C. simulans*-SDS-PAGE (16 kDa slice) are highlighted in yellow, DECOY-peptides from the WB-16 kDa-band are highlighted in green, and DECOY-peptides originating from both, are highlighted in grey. AA-sequences of DT-variants and homologs (Beta 201, Beta 286, Beta 341, Beta 371, and homolog CRM228) revealed similar patterns.

Interestingly, the highly abundant immunopeptide no.136 (Tab.S1) reveals a striking sequence-similarity with the iron ABC (ATP-Binding Cassette) transporter of *Corynebacterium diphtheria* and other *Corynebacterium* taxa. Since iron is essential for the regulation of diphtheria toxin expression^65^, the corresponding *C. simulans* protein could represent a great target of antibodies. Iron ABC transporters are important bacterial virulence factors and targets for the development of antibacterial vaccines and therapies^66^.

The majority of bacterial pathogens contain prophages, which often encode virulence factors^67^. Whereas *Vibrio cholera*, Shiga toxin-producing *E. coli*, *Corynebacterium diphtheriae*, and *Clostridium botulinum* depend on a specific prophage-encoded toxin for causing a specific disease, *Staphylococcus aureus*, *Streptococcus pyogenes*, and *Salmonella enterica serovar typhimurium* harbor a multitude of prophages and each phage-encoded virulence or fitness factor make an incremental contribution to the fitness of the lysogen^68^.

In summary, since a BLASTP® search of several DECOY-peptides revealed a high similarity to diphtheria toxin (DT) and its variants and homologs (Tab.S1 and Tab.S2, Fig.2), I would propose that in psoriasis lysogenic *C. simulans* phages release, upon triggering by yet unknown environmental factors, immune-responses-provoking DT-like toxin(s) **(“Psoriasis-Toxin”**).

### Bispecific Antibodies Against *C. simulans* Antigens Also Recognize Epidermal Proteins

Peptidomic analyses of the WB experiments to my surprise revealed also the presence of high numbers of individual peptides of human origin, which were no common contaminants. As shown in Tab.2, highly abundant peptides (10^11^ - 10^13^ TICs!) from the proteins hornerin (16 peptides), filaggrin (19 peptides), desmoplakin (3 peptides), keratin 1 (1 peptide), keratin 2 (2 peptides), keratin 6C (1 peptide), keratin 9 (4 peptides), caspase 14 (4 peptides), a linearized and truncated form of the serine protease inhibitor SPINK9 (8 peptides), and suprabasin (3 peptides), which play an important role in the development of the stratum granulosum^69^, were found in the WB-bands of heat-treated bacteria. Strikingly, many peptides have a nearly identical primary structure and differ only in one or two AA. For example, when in highly abundant FLG peptides e.g. a polar AA is replaced by Tyr or a Tyr is replaced by a polar AA (indicated in bold and red letters), the peptide’s abundance decreased by the factor 10^6^ (Tab.2), which may indicate the presence of high- and low-affinity autoantigens in psoriasis. Peptide length of the discovered autoantigens was found to be between 8 and 28 residues with the majority being 11-12-mer peptides (Tab.2). When looking for the composition of the peptides, it is striking that the identified peptides are rich in disorder-promoting amino acids. Thus, most of the 64 peptides originate from intrinsically disordered protein regions (IDPRs) of the respective proteins. The high abundance of, at least at acidic pH (as seen in healthy skin), cationic amino acids, indicates that these fulfill the structural criteria for being CIDAMPs^60^. The repeat-rich S100 fused-type protein hornerin^70^ is the most abundant CIDAMP source of healthy human skin. Also, filaggrin is a CIDAMP source, however only its cationic spacer regions^60^. Interestingly, all FLG peptides found in the WB experiment, except two, were from the spacer regions of FLG (Tab.2).

**Table 2:**
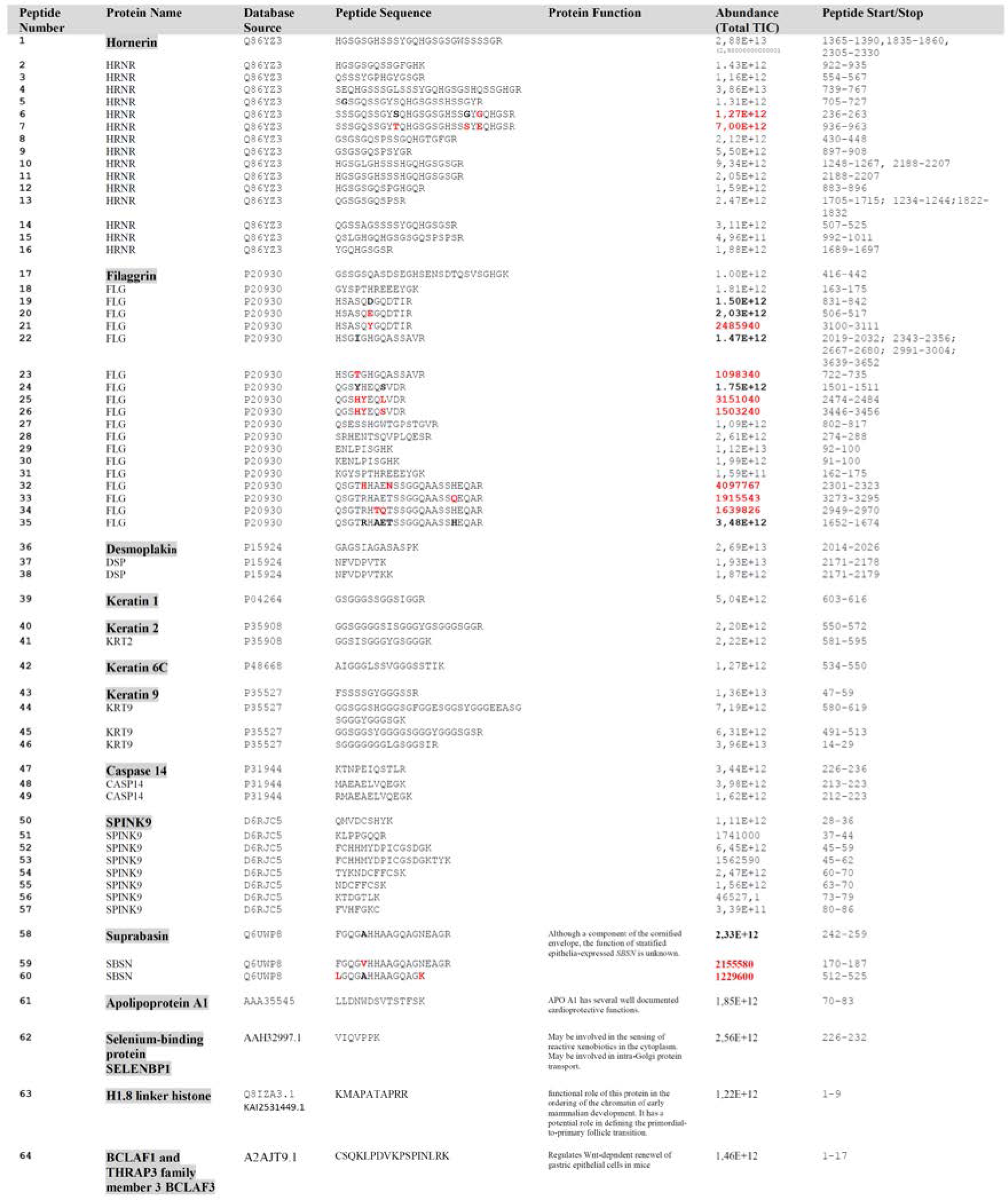
*Homo sapiens*-derived peptides identified within the immunoreactive 16 kDa *C. simulans*-WB-Band. The 16 kDa-WB-band of a freshly generated, heat-treated *C. simulans* extract was cut out and analyzed by proteomic analyses of on-membrane trypsin-digested proteins by LC-MS/MS. Analyzes were performed to identify human proteins using Scaffold Software^TM^ (version 5.2.1). In the Universal Protein Resource (UniProt) databank identified *Homo sapiens* proteins are shown as Peptide Report. The relative abundance of the peptides is given as “TICs” (Total Ion Chromatograms). Extraordinary high numbers of TICs (mostly >10^12^) were found for immunopeptides in comparison to contaminating proteins (up to 10^4^ TICs). All identified peptides originate from proteins synthesized within fully differentiated keratinocytes within the stratum granulosum. Note some marked quantitative differences of immunopeptides containing one or two AA exchanges (marked bold and red).

Search for additional, highly abundant (10^12^ TICs) immunopeptides revealed also apolipoprotein A1 as a major, non-epidermal autoantigen. Apolipoprotein A1 is an emerging cardiovascular risk factor of cardiovascular diseases^71^ and the presence of autoantibodies may help to explain why psoriasis is an independent cardiovascular risk factor^72^. A Selene-binding protein was also found (Tab.2). Its function is unknown. Maybe, it is involved in the sensing of reactive xenobiotics in the cytoplasm or involved in intra-Golgi protein transport. Interestingly, this immunopeptide is almost identical to a peptide from the *Corynebacterium* YihY/virulence factor BrkB family protein Sle (peptide 371, Tab.1). Another immunopeptide came from the H1.8 linker histone. H1 linker histones are cationic IDPs and key chromatin architectural proteins facilitating the formation of higher-order chromatin structures. They are essential for mouse development and proper stem cell differentiation. Their diversity, sequence conservation, complex expression, and distinct functions suggest that H1s mediate chromatin reprogramming and contribute to the large variations and complexity of chromatin structure and gene expression in the mammalian genome^73^. Another highly abundant immunopeptide is from BCLAF3 (regulator of apoptosis and transcription, B-cell lymphoma-2-associated transcription factor 3). BCLAF1 is associated with a multitude of biological processes, such as DNA damage response, splicing, and processing of pre-mRNA, T-cell activation, lung development, muscle cell proliferation and differentiation, autophagy, ischemia-reperfusion injury, and viral infection^74^. In a mouse model, BCLAF3 may induce differentiation in gastric glands by suppressing Wnt signaling and *Reg* family gene expression. It is suggested that *Bclaf3* likely regulates epithelial differentiation by controlling the expression of members of the *Reg* gene family, which are known to be potent modulators of epithelial renewal and stem cell maintenance^75^. Whether BCLAF3 has a role in psoriasis pathogenesis warrants further research.

In addition to the highly abundant epitopes of human autoantigens, 128 low-abundance immunopeptides (10-10^6^ TICs) were seen (Tab. S2). At least 48 selected peptides may have relevance in psoriasis pathogenesis. Here the epidermal protein dermokine was identified as autoantigen revealing three epitopes. It is of note that dermokine αβγ-deficient mice exhibited aggravated phenotypes in psoriasis-like dermatitis models, and showed barrier gene signatures similar to that seen in psoriasis^76^. A deficiency of dermokine leads to a transient cornification defect^77^. Other peptides represent epitopes of prostaglandin E2 receptor EP4, Vitamin D3 receptor, Interleukin-10 receptor subunit beta, Histamine H1 receptor, and the mineralocorticoid receptor. Further, epitopes of the Cw- 14 alpha chain of HLA class I histocompatibility antigen, Interleukin-12 subunit beta, Interleukin-12 subunit alpha, and IL-8 were detected. In addition, an epitope of nuclear receptor coactivator 2 was seen. This protein may also have a role in psoriasis because it controls immune tolerance by promoting induced T_reg_ differentiation^78^. Another low-abundance immunopeptide comes from the human endogenous retrovirus K (HERV-K). This immunopeptide (peptide no.75, Tab.S2) surprisingly shows high similarity to peptide sequence motifs of a *C. simulans* AAA-ATPase and Chain A of Diphtheria toxin, suggesting cross-reactivity of the HERV-K immunopeptide with proteins of *C. simulans* and DT. HERV-K is the most recently acquired endogenous retrovirus, which is activated and expressed in many cancers, and its subgroup, HML-2 is known to be elevated in a long list of inflammation-associated diseases^79^, including psoriasis^80,81^, where the HERV-K- dUTPase constitutes a candidate gene for the PSORS1 mutation^80^. HERV-K is upregulated in lesional psoriatic skin^82^ and proteome array studies revealed that treatment of human primary cells with HERV-K dUTPase proteins triggered the secretion of Th1 and Th17 cytokines involved in the formation of psoriatic plaques, including IL-12p40, IL-23, IL-17, tumor necrosis factor-α, IL-8, and CCL20^81^. The presence of an antibody against the zinc finger of the HERV-K Gag protein in psoriasis serum (peptide no. 75, Tab.S2) confirms the expression of HERV-K in psoriasis.

### Natural BsAbs Against *C. simulans* Antigens Recognize Cancer-related Proteins

Further analyses of low-abundance immunopeptides revealed 47 peptides originating from cancer-related proteins as autoantigens (Tab.S2). Among others, epitopes of cancer/testis antigen 47A (CT47), serologically defined colon cancer antigen 3 (SDCCAG3), breast cancer type 2 susceptibility protein (BRCA2), melanoma-associated antigen 6 (MAGEA6), ribonuclease 4 (RNASE4), and, as mentioned above, a Gag polyprotein viral zinc-finger-peptide of the human endogenous retrovirus protein K (HERV-K) were found. The latter is of particular interest because HERVs are critical for several physiological activities^83^. Although typically transcriptionally silenced in normal adult cells, dysregulation of HERV-K (HML-2) elements has been observed in cancer, including breast, germ cell tumors, pancreatic, melanoma, and brain cancer^84^. Aberrant expression of HERV-K has been shown to promote the expression of stem cell markers and promote dedifferentiation. Therefore HERV-K seems to be a potential therapeutic target based on evidence that some tumors depend on the expression of its proteins for survival^84,85^. HERVs and their products (including RNA, cytosolic DNA, and proteins) are still able to modulate and be influenced by the host immune system, suggesting a central role in the evolution and functioning of human innate immunity^86^. Indeed, HERV sequences had been major contributors in shaping and expanding the interferon network, dispersing inducible genes that have been occasionally domesticated in various mammalian lineages. Also, the HERV integration within or near genes encoding for critical immune factors has been shown to influence their activity. It also can be responsible for their polymorphic variation in the human population, such as in the case of the HERV-K provirus in the major histocompatibility complex region^86^. HERV-expressed products have been shown to modulate innate immunity effectors and therefore have been proposed to establish a protective effect against exogenous infections. These immune reactions often are related on the one side to inflammatory and autoimmune disorders, on the other side, however, to the control of excessive immune activation through their immunosuppressive properties^86^. The discovery of an epitope of the Gag polyprotein viral zinc finger of HERV-K may help to develop peptide vaccines as possible therapies against tumors^85^.

### A Proposed Mechanism of Bispecific Autoantibody-formation

How will BsAbs be formed? I propose that at first, *C. simulans* bacteria will be killed within the stratum corneum by various CIDAMPs, as already shown for elongated peptides containing sequences of peptides numbers 6,7 and 15 (Tab.2)^60^. These CIDAMPs, which may have been generated from major CIDAMP sources within the stratum corneum by bacterial proteases (e.g. from the in lesional psoriatic skin highly abundant obligate anaerobic commensal *Finegoldia magna*^17,87^ enter the bacterial membrane, bind to ribosomal proteins and kill the bacterium by formation of toxic fibers generated by CIDAMP/bacterial disordered protein complexes and/or possibly inhibit protein synthesis, which eventually leads to its lysis^50^. Complexes of CIDAMPs with ribosomal proteins or other bacterial or phage-derived IDP(R)s would form chimeric antigens, which consist of short peptide sequence motifs of both, bacterial (e.g. ribosomal proteins) and/or phage-derived IDP(R)s, and a CIDAMP as proposed in Fig.3. These will be taken up by APCs, and B-cells will then generate BsAbs, which recognize chimeric antigens originating from both, bacterial and/or phage IDP(R)s, and CIDAMPs. Thus, it is tempting to speculate that a CIDAMP-based, persisting epithelial defense reaction within the stratum corneum and stratum granulosum towards virulent *C. simulans* could eventually result in the generation of CIDAMP-like autoantigens. Since CIDAMPs are generated from cationic IDP(R)s, which are highly abundant in keratinocytes^52^, and are produced within the fully differentiated epidermal layers, it would be plausible to see CIDAMP-directed immune reactions, such as IgG- deposits, within the stratum corneum^3,88^ and autoreactive T-cell-mediated^89,90^ responses within the stratum granulosum. Keratinocyte-derived CIDAMPS as autoantigens would also help to explain the typical gyrate, demarcated lesion morphology of annular psoriasis^91^, where I would propose that the center is missing CIDAMP sources due to a loss of the stratum granulosum. This hypothesis is supported by the hornerin (HRNR) expression profile in psoriasis, which is only seen within the scaly and pustular margin of annular lesions^92^.

**Figure 3:**
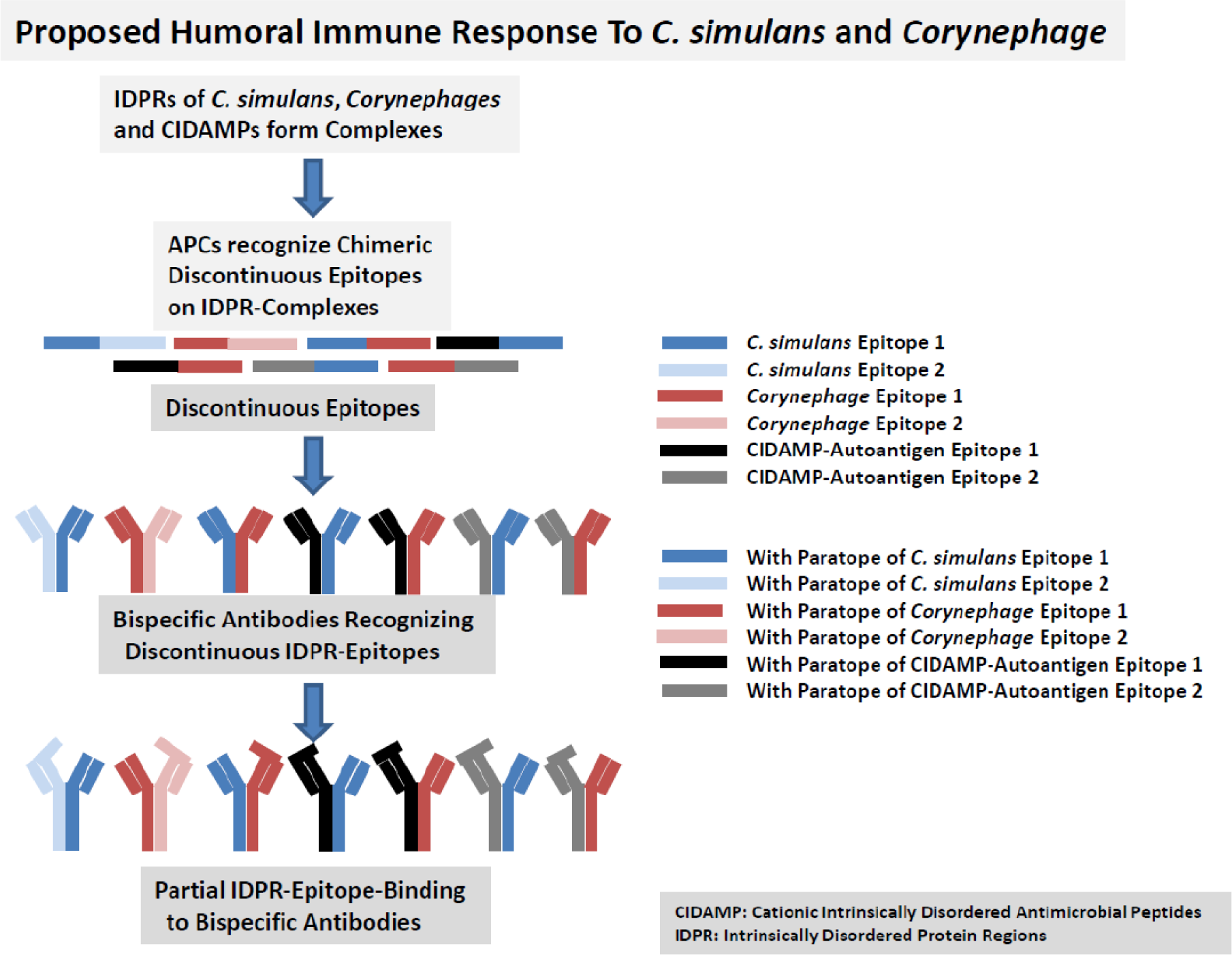
Proposed Humoral Immune Responses To *Corynebacterium simulans* in Psoriasis. Lysis of stratum granulosum cells would lead to a premature liberation of CIDAMP sources and other IDPs and IDPRs from stratum granulosum cells. Proteases from bacteria (e.g. *Finegoldia magna*) may generate CIDAMPs from its precursors. These may bind to bacterial and viral IDP(R)s, including ribosomal proteins of *C. simulans*, forming IDP(R)-complexes. Such complexes may serve as potential antigens, which contain chimeric discontinuous epitopes. Processing of these complexes within the APC will generate peptides consisting of discontinuous epitopes from different IDP(R)s. B-cells will then generate from these epitopes bispecific antibodies, which are bivalent and able to bind two different epitopes. This occurs in psoriasis, where the free paratope binds to epitopes of the yet unknown 16 kDa antigen(s) of *C. simulans* and/or the proposed *Corynephage*. The second paratope is already occupied by peptides, representing epitopes originating from *C. simulans* (and *C. kroppenstedtii*), *Corynephages*, or *C. simulans-cidal* CIDAMPs. These CIDAMPs, which may have been generated after the destruction of stratum granulosum cells, form complexes with *C. simulans*/*Corynephage* IDP(R)s, and represent autoantigens, which may cause autoimmune reactions within the stratum granulosum of psoriatic lesions.

## Major Open Questions

Immuno-proteomic and -peptidomic analyses of WBs with pooled psoriasis IgG for *C. simulans* antigens revealed a multitude of epitopes of bacterial and putatively corynephage origin as well as CIDAMP- and IDP(R)-like autoantigens. Since the nondiphtheria corynebacteria have emerged as important pathogens and several species produce toxins, including a diphtheria-like toxin, a dermo necrotic toxin, and a soluble hemolysin^93^, it would be important to fully characterize the genomes of psoriasis-derived *C. simulans* (and *C. kroppenstedtii*) strains using improved technology^34^. Since at least some strains may contain prophages, their genomes need to be characterized. WB-based immuno-peptidome analyses together with a BLASTP® search (limited to *Corynephage beta* (taxid:10703)) revealed for a high number of peptides a similarity to diphtheria toxin and DT-variants and homologs (Fig.2, Tab.S1) favoring the hypothesis of the existence of a DT-related cell-toxic “Psoriasis Toxin” with relevance in psoriasis pathogenesis. Further, it would be important to investigate, whether the *C. simulans* and *C. kroppenstedtii* strains and/or corynephages found in psoriasis lesions may correlate with the disease acuity. A personalized immuno-peptidomic analysis for bispecific IgG-bound epitopes of bacteria- and corynephage-derived antigens as well as epitopes of autoantigens would show, whether these correlate with clinical parameters (e.g. g. PASI score, psoriasis subtypes, anatomic location, and morphology of the lesions). Also, it would be interesting to ask whether variation in the disease site with the involvement of the scalp, palmoplantar region, genitals, and nails may correlate with the tissue expression profiles of the identified autoantigens, together with toxigenic *C. simulans* in the local microbiome.

Infections with *Streptococcus pyogenes* and *Staphylococcus aureus* are well-known trigger factors for psoriasis^10^. Interestingly, BLASTP® search of *C. simulans* antigens revealed similarity to *S. pyogenes*- and *S. aureus* proteins as well as proteins of *S. pyogenes* phages and *S. aureus* phages (Tab.2; Tab.S1). Since both pathogens are targets of selected CIDAMPs^60^, WB-based immuno-peptidomic analyses of psoriasis patient’s serum would show, whether also *S. pyogenes*- and *S. aureus*-antigens are parts of discontinuous and chimeric autoantigens. This would help to explain, why infections with these pathogens can trigger psoriasis. Further, WB-based immunopeptidomic analyses of *C. kroppenstedtii*, *Neisseria* spp., and *Finegoldia* spp., which are also increased in the psoriasis skin in comparison to healthy skin^17^ would show whether these taxa have a role in psoriasis pathogenesis.

## Conclusion

Since commensal microbiota regulates host immunity to pathogens^94^, it is not surprising that microbiota is also capable of regulating autoimmune responses by affecting the targets of autoimmunity directly^95^.

The observation that *C. simulans* is markedly increased in lesional and nonlesional psoriatic skin^17,96^ prompted me to hypothesize that it may have a role in psoriasis etiology. With the limited data shown in this exploratory study and the putative and unique proinflammatory properties, I propose that initially, toxigenic *C. simulans* may infect skin micro-wounds (Fig.4). Prophage-containing *C. simulans* strains may produce upon activation a DT-like “Psoriasis-Toxin”. This may be a homolog of DT, which may act similarly as DT by binding to heparin-binding epidermal growth factor-like growth factor (HB-EGF)^97^, which is highly expressed in keratinocytes^98^ and is thought to be implicated in psoriasis^99^. I would speculate that this proposed toxin may cause keratinocyte-intrinsic ribotoxic stress and NLRP1 inflammasome activation, similar as seen in a model of cutaneous diphtheria^100^. In support of this hypothesis, NLRP1 is regarded as the principal inflammasome sensor in human keratinocytes^101^, and has a role in psoriasis^102^. Further, NLRP1 activation in human keratinocytes cultivated in organotypic skin cultures induces a psoriasis-like phenotype^103^. Similar to DT, “Psoriasis Toxin” might selectively kill keratinocytes of the stratum granulosum by inhibiting protein biosynthesis via ADP-ribosylation^104^. Destruction of stratum granulosum cells by “Psoriasis Toxin” would result in a diphtheria-like skin infection. Toxic effects on stratum granulosum cells may liberate CIDAMP precursors, e.g. HRNR and FLG (Fig.4). These may be cleaved by proteases of *Finegoldia spp*., an obligate anaerobic commensal residing in deeper layers of the stratum corneum^105^ which is also increased in psoriasis^17^, and is a source of a serine protease as virulence factor^106,87^, resulting in a multitude of bioactive CIDAMPs (Fig.2). Among the CIDAMPs generated, some may kill *C. simulans* by targeting the ribosome^50^, forming complexes of CIDAMPs with bacterial IDP(R)s like ribosomal proteins. These IDP(R)-complexes may arise from the quaternary structure of a multimeric assembly of CIDAMPs and bacterial IDP(R)s, either from the juxtaposition of residues in neighboring IDP(R)s that are recognized by Abs as a single epitope or from conformational changes induced in the IDP(R) by intersubunit interactions. Such IDP(R)-complexes would represent chimeric antigens from *C. simulans*, its prophages and/or CIDAMPs, bearing scattered discontinuous epitopes (Fig.3). In consequence, binding of these chimeric antigens to MHC-II would result in the generation of bispecific antibodies, which recognize epitopes from two neighboring IDP(R)s, including CIDAMPs as autoantigens. Chronicity and severity of *C. simulans* skin infection in psoriasis would cause an increase of antibodies towards this pathogen and also towards CIDAMPs as autoantibodies (Fig.4). It is likely that in chronic plaque psoriasis CIDAMP-autoantibodies would interfere with the wound healing process, where formation of the new stratum granulosum with its high numbers of potential CIDAMP sources would cause inflammation. Maybe, this could be important in the Koebner phenomenon^104^, where upon wounding the local generation of CIDAMP-autoantigens would trigger psoriasis. On the other side, areas lacking an intact stratum granulosum would also lack the source of CIDAMP-autoantigens and the cellular target of the proposed “Psoriasis Toxin” and thus inflammation.

**Figure 4:**
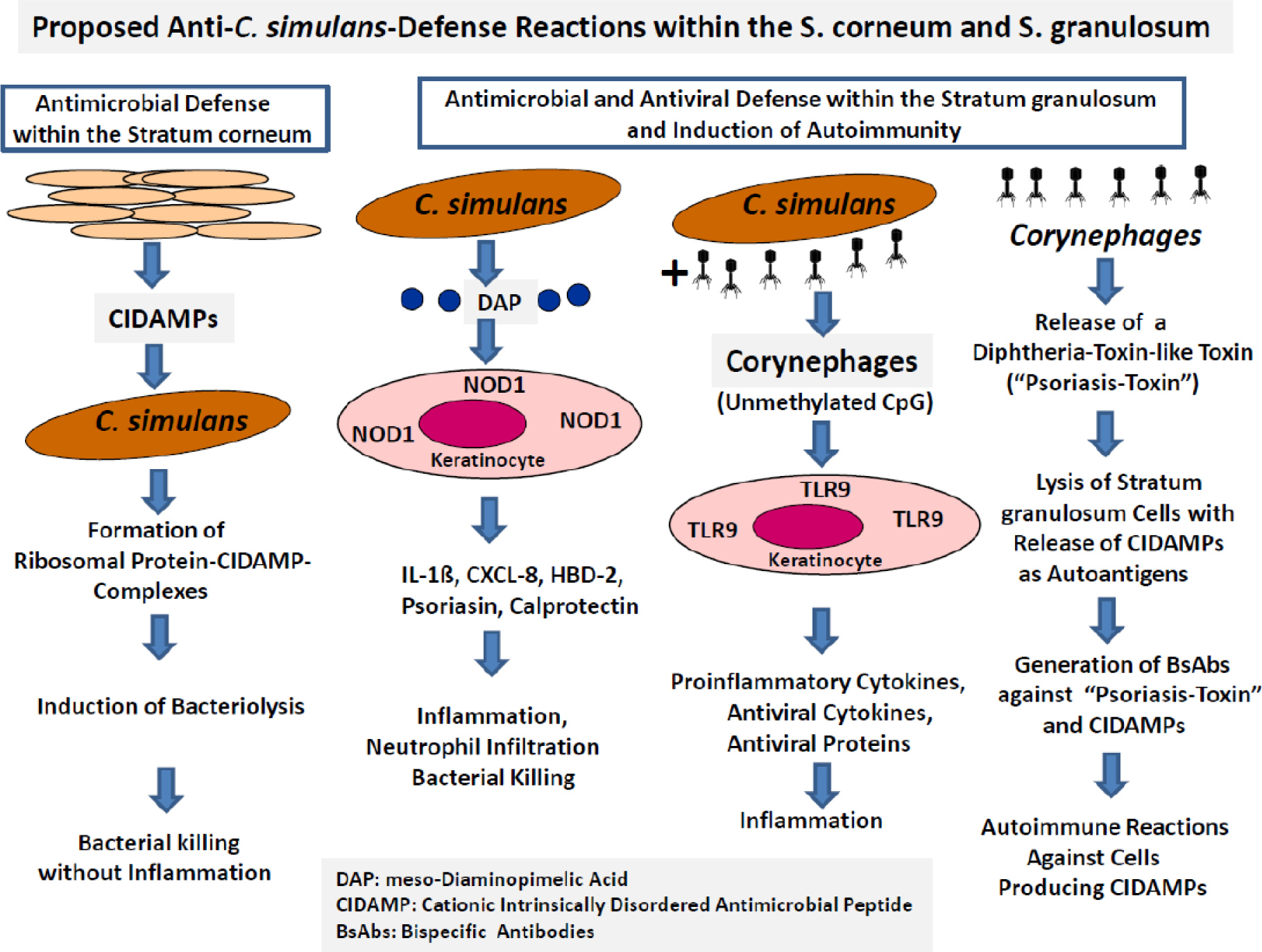
Proposed possible Etiology of Psoriasis. *Corynebacterium simulans* colonizes within the Stratum corneum, and its abundance will be controlled by stratum corneum-derived Cationic Intrinsically Disordered Antimicrobial Peptides (CIDAMPs)^60^, which are generated from CIDAMP sources like hornerin and filaggrin, without inducing inflammation. *C. simulans*, present in the deepest stratum corneum layers, release the peptidoglycan meso-diaminopimelic acid (DAP), which binds to the pattern recognition receptor NOD1 present in keratinocytes. This leads to the activation of signaling cascades with the induction of inducible antimicrobial peptides like hBD-2, S100- A7, and calprotectin, the direct and indirect induction of neutrophil-recruiting cytokines as well as antiviral defense processes. *C. simulans* strains may be infected by bacteriophages (“Corynephages”). The *C. simulans* cellular machinery replicates the viral genome and catalyzes the self-assembly of viral particles, leading to bacterial lysis and the release of infectious virions. These are rich in unmethylated CpG, a natural ligand for TLR9, activating keratinocytes to induce proinflammatory cytokines, antiviral cytokines, and antiviral proteins, leading to inflammation. Some *C. simulans* strains can initiate either direct bacterial lysis or contain prophages. These, in the bacterial chromosome integrated phages, can remain in this state or switch to a lytic cycle with an inducible release of cell-toxic proteins. A high percentage of antigenic peptides identified in this study (Tab.S1 and S2, Fig. 2) revealed similarity to the catalytic domain-containing A-chain of diphtheria toxin (DT) and various variants and homologs, supporting the hypothesis that in psoriasis *C. simulans*, like *C. diphtheriae*, contains lysogenic prophages, which may release a DT-related lytic protein (“**Psoriasis Toxin**”) that may destroy stratum granulosum cells.

## Perspectives

If future studies confirm *C. simulans* as a critical trigger factor in psoriasis, it would be interesting to target this commensal, either by a selective topical disinfectant or to block meso-DAP-caused epithelial proinflammatory cytokine- and interferon-release by topical NOD1-antagonists. Current pharmacological strategies pursuing NOD1 blockade have been recently described^107^.

The most intriguing finding of this exploratory study is the existence of natural BsAbs and IgG-bound epitopes in psoriasis serum. This allowed the characterization of psoriasis autoantigens from a serum pool, such as a multitude of CIDAMP-like peptides, and, importantly, the discovery of epitopes on IDPs or IDPRs of *C. simulans* and putative *Corynebacteriophages*. Previously, it has been suggested that IDPs will elicit weak immune responses or even be completely non-immunogenic^108^. Recent studies, however, have shown disordered antigens are important targets of natural immune responses to several pathogens, especially viruses, and are increasingly attracting attention as potential vaccine components^109^. Structural investigations have demonstrated that disordered epitopes are smaller than ordered epitopes and bona fide targets of high-affinity and specific antibody responses - an observation that was confirmed in this study (Tab.1, Tab.2, Tab.S1).

BsAbs-based immunopeptidome analyses of WBs - as shown in this limited study – allow the molecular epitope characterization of intrinsically disordered antigens of bacterial and virus origin, self-antigens, and possibly cancer/testis antigens^110^, which should be useful for improved peptide-based vaccines^111^. These form one of the most potent vaccine platforms, offer exclusive advantages over classical vaccines that use whole organisms or proteins, provide precise activation of immune responses, and have great potential for personalized immunotherapy^111^. Maybe, psoriasis patients could benefit from a peptide-based vaccination with immunopeptides of *C. simulans* and/or their *Corynephages*, in particular, the proposed “Psoriasis Toxin”. Alternatively, they could benefit from autoantigen-specific therapies. A therapy with vaccines carrying autoantigenic peptides establishes a specific immune tolerance by delivering peptide fragments containing disease-specific self-antigen epitopes to suppress excessive immune responses, thereby exerting a therapeutic effect, with high safety and specificity^112^. Due to a defect in regulatory T cells (Treg) in psoriasis^113^ such a therapeutic approach could be promising, in particular with antigen-specific Treg cells, which are believed to be more potent in regulating and improving immune tolerance in a disease-specific manner^114^. However, antigen-specific Treg cell therapies in the past were challenged by the identification of disease-relevant antigens. With the discovery of natural BsAbs as sources of epitope peptides, nonetheless, this should no longer be a problem for psoriasis and possibly many other autoimmune diseases.

## Material and Methods

### Psoriasis serum

A pool of surplus serum, obtained from hospitalized patients with psoriasis, was used in this study. Serum samples were originally taken for determination of chemokine levels in a clinical study (performed 1994-1998). This study has been approved by the local ethics commission^115^. The pooled serum was stored below -70°C until use.

### Isolation of IgG from psoriasis serum

rProtein A Sepharose® gravity-flow columns (Sigmaaldrich), prepacked with 1 mL of Protein A Sepharose® 4 Fast Flow, have been used for IgG isolation. Briefly, serum was diluted with binding buffer (0.02 M sodium phosphate, pH 7.0) and applied according to the protocol given by the manufacturer. IgG was eluted with 0.1 M glycine-HCl, pH 2.7. To preserve the activity of acid-labile IgG, samples were then immediately neutralized to pH 7.0 with 1 M Tris-HCl, pH 9.0.

### Culture of Corynebacterium simulans

*C. simulans* (strain DSM 44415)^22^ was grown for 24 h at planktonic condition (shaking),(37 °C at 170 rpm) as recommended by Leibniz Institute DSMZ-German Collection of Microorganisms and Cell Cultures in PYG medium 104 (pH 7.2). Cultures at static conditions were performed without shaking.

### Lysis of C. simulans

Cells from the bacterial culture were harvested by centrifugation (5000 rpm for 10 minutes). After aspirating the supernatant, the pellet/bacterial cells were frozen at -70 °C, resuspended in 1 ml MQ grade water, and then lysed with 1% Triton X-100.

### Heat lysis of *C. simulans*

For heat lysis, bacterial cells were resuspended in 1% Triton X-100, and heated at 95 °C for 20 min. Heat lysates were then chilled on ice for 10 min. For all lysates, cell debris was pelleted at 10,000× g for 30 min.

## Western Blot analyses

*C. simulans* lysates, either untreated, heated at 95 °C, or sonicated (using three 10-second bursts at high intensity), were applied in DTT (50mM)-containing loading buffer. For Western Blot analysis, samples (14 µl each) were separated on a 12% 37.5:1 acrylamide:bisacrylamide gel. Proteins were then transferred to a nitrocellulose membrane (pore size: 0.2 µm, Schleicher & Schuell BioScience, Dassel, Germany) using an alkaline transfer buffer (48 mM Tris, 39 mM glycine, 0.0375% (w/v) SDS and 20% EtOH (pH 9.2)). An alkaline transfer buffer is essential to get cationic proteins efficiently transferred to a membrane. After blocking for 1 h in blocking buffer (5% (w/v) bovine serum albumin (BSA) in PBS/Tris, pH 7.4 + 0.05% Tween) and after washing with PBS/Tris, the membrane was incubated with psoriasis IgG (2 µg/mL) for 3 hrs, followed by washing with PBS/Tris. The membrane was then incubated with goat anti-human IgG-HRP (Jackson) 1:10,000 at 4 °C overnight. After an additional washing step (6×) with PBS/Tris, pH 7.4, membranes were incubated with a peroxidase substrate (Roche Lumilight Western Blotting Substrate No. 12015196001) at ambient temperature and documented with a “Diana III Digital CCD Imaging System”.

### Proteomic and Peptidomics analyses of WB bands

Proteomic analyses were performed as a service by the Proteomic Facility at King’s College London. According to a protocol for in-gel digestion provided by the Proteomic Facility, the electrophoretically transferred 16 kDa-band of the Western Blot experiment and a corresponding 16 kDa SDS-PAGE section of a *C. simulans* extract were excised from the nitrocellulose membrane, washed with triethylammonium bicarbonate (TEAB) buffer, reduced with dithiothreitol solution in TEAB, alkylated with iodoacetamide in TEAB and then on-membrane digested with trypsin (Sigma, bovine sequencing grade) before analysis by LC-ESI-based mass spectrometry.

### Data analysis

LC/MS/MS data were analyzed with the Scaffold Software^TM^ (version 5.3.0) (https://www.proteomesoftware.com/products/scaffold-5).

#### Acknowledgements

I have to give credit to Davide Pennino, Ulrich Gerstel, and Britta Hansmann, who helped with pilot experiments and greatly thank Anke Rose and Jutta Quitzau for technical assistance. D. P. was supported by the MAARS (Microbes in Allergy and Autoimmunity Related to the Skin) project and U. G. and B. H. were supported by the author’s high-risk Reinhart Koselleck-DFG-project “Resistance-avoiding Antimicrobial Peptides of Healthy Human Skin”. I am very much obliged to Enno Christophers for encouraging me to write down my thoughts to the psoriasis etiology after being in retirement, to re-analyze old data with new software and for helpful discussions about the role of the stratum granulosum in psoriasis pathology.

