## Supplementary material for "Discovery of Natural Bispecific Antibodies: Is Psoriasis Induced by a Toxigenic *Corynebacterium simulans* and Maintained by CIDAMPs as Autoantigens?": Suppl Table S1

### Supplementary Table S1

| Peptide-Number | Protein accession numbers | Sequence;<br>Alignment with Chain A, DIPHTHERIA TOXIN [Corynephage beta]1DDT_A | Blastp (protein-protein BLAST®)-Search<br>Viruses (taxid:10239)<br>Corynephage beta (taxid:10703)<br>Corynebacterium diphtheriae (taxid:1717) | Abundance (TIC) |
|---|---|---|---|---|
| 1 | C4LJI9- DECOY | (?)LNVTTTAGLAAER(?)<br>Query 1 LNVT 4<br>LNVT<br>Sbjct 30 LNVT 33 | immunity-specific protein Beta371 [Corynephage beta]AAA32187.1; Endolysin [Siphoviridae sp. ctGJ32] DAF86071.1 ; lysin [Streptococcus phage Javan73] QBX22442.1; gp21, tail fiber protein [Corynebacterium phage BFK20] YP_001456751.1; protease [Stenotrophomonas phage phiSHP3] QNO13254.1; tailspike protein [Bacteriophage sp.] DAX06982.1; tail spike protein [Podoviridae sp.] DAR52772.1; LCP family protein [Corynebacterium ulcerans] WP_029974719.1 | 4435000 |
| 2 | C4LJE9- DECOY | (?)LAPAFISK(?)<br>Query 1 LAPAFIS 7<br>L PAF+S<br>Sbjct 465 LTPAFVS 471 | MDR family MFS transporter [Corynebacterium diphtheriae]WP_072564639.1; tail tape measure protein [Staphylococcus phage StB12] YP_009130737.1<br>tape measure protein [Gordonia phage Hollow] UYL87047.1; tail tape measure [Siphoviridae sp.] DAI76915.1 | 4768000 |
| 3 | C4LH89- DECOY | (?)LSSQEILR(?) | tail tape measure protein [Bacteriophage sp.] DAW97943.1; neuraminidase [Influenza A virus (A/Cambodia/W0908340/2012(H3N2))] ALX27856.1 | 818500 |
| 4 | C4LLD8-DECOY | (?)SPPSQTLVEAGK(?) | hypothetical protein IPP34_00013 [Streptococcus phage IPP34] APD22903.1; tail length tape-measure protein [Streptococcus phage Javan411] QBX18339.1; integrase [Roseobacter phage CRP-345] UAW59104.1; major capsid protein [Mycobacterium phage Phantastic] YP_009032500.1 | 1155000 |
| 5 | C4LH04-DECOY | (?)VAPAVIPDK(?) | polyprotein [hepatitis C virus genotype 1a] ACJ04213.1<br>Phage-like element PBSX protein, XkdF [uncultured Caudovirales phage] CAB5208321.1 | 870500 |
| 6 | C4LJH-DECOY | (?)INERTRDLK(?) | head scaffolding protein [Escherichia phage RB49] NP_891731.1<br>hypothetical protein [Bacteriophage sp.] DAQ92370.1<br>Tail tape measure [Bacteriophage sp.] DAK35754.1 | 1455000 |
| 7 | C4LG76-DECOY | (?)FLPAEDAR(?)<br>Query 4 AEDA 7<br>AEDA<br>Sbjct 212 AEDA 215 | immunity-specific protein Beta371 [Corynephage beta]AAA32187.1;hypothetical protein [Corynebacterium diphtheriae] WP_235696814.1; putative recombinase [Bacteroides phage Bacuni_F1] QMS42057.1; structural protein [Pectobacterium phage vB_PatP_CB1] YP_009832293.1; tail fiber protein [Pectobacterium phage Nepra] YP_009837897.1; ThyX-like thymidylate synthase [Gordonia phage Yikes] YP_009854038.1 | 3745000 |
| 8 | C4LIA8-DECOY | (?)QLNTGTNPLR(?)<br>Query 4 TGTNP 8<br>TGTNP<br>Sbjct 267 TGTNP 271<br><br>Query 3 NTGTNP 8<br>NTGTNP<br>Sbjct 519 NTGTNP 524 | Chain A, DIPHTHERIA TOXIN [Corynephage beta]1DDT_A; diphtheria toxin [Corynebacterium diphtheriae phage]CAA25302.1; N-acetylmuramoyl-L-alanine amidase [Corynebacterium simulans] WP_248093839.1; glycosylase [Corynebacterium diphtheriae] OKY24467.1 antirepressor protein KilAC domain [Bacteriophage sp.] UVX35941.1; ankyrin-like protein [Vaccinia virus] AIZ72765.1; tail sheath protein [uncultured Caudovirales phage] CAB4122182.1; ankyrin repeat-containing protein [Cowpox virus] ADZ24028.1; Ankyrin [Akhmeta virus] QEQ49571.1 | **1,41E12** |
| 9 | C4LG79-DECOY | (?)ETCQIIMR(?) | ATP-dependent DNA helicase [Ochrobactrum phage vB_OspM_OC] QIG65687.1; PB1-F2 protein [Influenza A virus (A/chicken/Hong Kong/YSC82/2008(H9N2))] AGQ83450.1; ScFv10, Lysozyme C [Siphoviridae sp.] DAP77197.1 ; ORF6 protein [Severe acute respiratory syndrome coronavirus 2] UJO97927.1; protease, partial [Human rhinovirus sp.] AAA45759.1 | 1763000 |
| 10 | C4LIK1-DECOY | (?)SSMSPLYPCAMDTPS(?) | Thymidylate synthase [Siphoviridae sp.] DAE94106.1 ; minor tail protein [Mycobacterium phage Sauce] YP_010058709.1; minor tail protein with lysin activity [Mycobacterium phage ArcherS7] YP_008061414.1 | 6092000 |
| 11 | C4LGG3-DECOY | (?)DLISPR(?) | DNA primase/helicase [Bacillus phage SP8] QPX71674.1; DNA helicase [Bacillus phage vB_BsuM-Goe2] APZ82345.1; pol protein, partial [Human immunodeficiency virus 1] ABN11033.1 | 2573000 |
| 12 | C4LGE2-DECOY | (?)TIVDVLK(?) | 1a polyprotein [Infectious bronchitis virus] QGM12377.1; hypothetical protein UFOVP1116_40 [uncultured Caudovirales phage] CAB4184855.1 ; DNA polymerase III, alpha subunit [Siphoviridae sp.] DAH16956.1 ; minor tail protein [Siphoviridae sp.] DAY93496.1 | 901200 |
| 13 | C4LJ80-DECOY | (?)INSEIVER(?)<br>Query 1 INSEIVER 8<br>I SE ER<br>Sbjct 173 IESERRER 180 | immunity-specific protein Beta371 [Corynephage beta] AAA32187.1;active core crystal toxin [Myoviridae sp.] DAI66323.1; endonuclease R [Acinetobacter phage MD-2021a] CAH1085376.1; tail length tape measure protein [Loktanella phage pCB2051-A] YP_007674929.1 | 1254000 |
| 14 | C4LIC6-DECOY | (?)FAREGVK(?) | lysostaphin [Staphylococcus phage PhiSepi-HH3] QPB07808.1 ; tail tape measure protein [Staphylococcus phage SPbeta-like] YP_009226727.1; PolyVal ADP-Ribosyltransferase [Bacteriophage sp.] DAF34700.1; surface glycoprotein [Severe acute respiratory syndrome coronavirus 2] UHG25195.1 | 1585000 |
| 15 | C4LGE8-DECOY | (?)LAPTTAAK(?) | tail tape measure protein [Klebsiella phage ST16-OXA48phi5.4] YP_009882552.1 ; prohead serine protease [Bacteriophage sp.] DAL57016.1; prohead serine protease [Siphoviridae sp.] DAX97315.1; envelope glycoprotein [Human immunodeficiency virus 1] AFE02602.1 | 1826000 |
| 16 | C4LH39-DECOY | (?)IQAEVAK(?) | Major capsid protein [Bacteriophage sp.] DAW90087.1; HNH endonuclease [Mycobacterium phage Lewan] QDK03905.1; hypothetical protein [Shigella phage ESh1] URY10670.1; protein 4.7 [Yersinia phage phiA1122] NP_848282.1; hypothetical protein EFA2_00026 [Enterococcus phage EFA-2] QJT70338.1; hypothetical protein T71t1_16 [Escherichia phage T7] UJQ71070.1 | 1963000 |
| 17 | C4LHE6-DECOY | (?)LGGSLSSTDVR(?)<br>Query 2 GGSLS 6<br>GGSLS<br>Sbjct 221 GGSLS 225<br><br>Query 1 LGGSL 5<br>LG SL<br>Sbjct 106 LGLSL 110 | Diphtheria toxin homolog CRM228 fragment A and B P00589.2; Chain A, DIPHTHERIA TOXIN [Corynephage beta]1DDT_A; lysin A [Mycobacterium phage SoSeph] QXN73772.1; gp12 [Corynebacterium phage P1201] YP_001468914.1; tail length tape measure protein [Myoviridae sp.] DAM93467.1 | 2696000 |
| 18 | CeLI70-DECOY | (?)VTVRDDAR(?) | DNA polymerase [Brevundimonas phage vB_BpoS-Papperlapapp] USN16279.1; portal protein [Bacteriophage sp.] UVX81375.1; portal protein [Siphoviridae sp.] DAP22120.1 | 9681000 |
| 19 | C4LLP5-DECOY | (?)KINEAGIAMVTLGK(?)<br>Query 4 EAGIAMV 10<br>EA + MV<br>Sbjct 219 EARVEMV 225 | immunity-specific protein Beta241 [Corynephage beta] AAA32184.1; Lysozyme [Siphoviridae sp.] DAJ55915.1; putative lysin [Corynebacterium phage P1201] YP_001468954.1; lysin [Streptococcus phage P0095] ARU13172.1; minor tail protein [Corynebacterium phage Lederberg] YP_009848996.1 | 2520000 |
| 20 | C4LLU6-DECOY | (?)LASDITR(?) | terminase large subunit [Pseudomonas phage Iggy] QEA09735.1 ; LW134 [Lumpy skin disease virus] AAN02859.1; tape measure protein [Gordonia phage CaiB] UOW93010.1 | 990900 |
| 21 | C4LKB5-DECOY | (?)RTVMRAK(?) | Spo0J Stage 0 sporulation protein J (antagonist of Soj) containing ParB-like nuclease domain [uncultured Caudovirales phage] UVY13838.1 ; hypothetical protein [Siphoviridae sp.] DAH80093.1 ; chromosome partitioning protein [Siphoviridae sp.] DAI76939.1; Large subunit terminase [Bacteriophage sp.] DAR03982.1 | 1799000 |
| 22 | C4G56-DECOY | (?)ANAFPTGPK(?) | tail spike protein [Escherichia phage C130_2] YP_009812583.1; 2OG-Fe(II) oxygenase [uncultured Mediterranean phage uvMED] BAR35486.1; putative RNA-dependent RNA polymerase [Infectious pancreatic necrosis virus] AAN04572.1; tat protein [Human immunodeficiency virus 1] ADJ00364.1<br>hypothetical protein FF83_gp272 [Serratia phage PS2] YP_009030190.1 | 1659000 |
| 23 | C4LLE4-DECOY | (?)IPYSDPGARNDR(?) | Toll-like receptor 8 rich repeat, RNA, Glycosylation [Bacteriophage sp.] DAW98581.1; tail tube protein [Bacteriophage sp.] DAI09615.1 | 4952000 |
| 24 | C4LKH4-DECOY | (?)VAALPSVR(?) | ddrB-like ParB superfamily domain protein [Bacteriophage sp.] UWH91342.1; Tail tape measure [Siphoviridae sp.] DAZ79689.1; DNA methylase [Podoviridae sp.] DAW50662.1; Inorganic Pyrophosphatase, partial [Bacteriophage sp.] UWI14060.1 | 1266000 |
| 25 | C4LJL8-DECOY | (?)RDDGGEEERIR(?)<br>Query 1 RDDG-----GEEE 8<br>RDDG GEEE<br>Sbjct 189 RDDGMVTLGGEEE 201 | phage tail protein [Corynebacterium diphtheriae] CAB0558168.1; hypothetical protein PE067_004 [Burkholderia phage PE067] ALJ98682.1; envelope glycoprotein [Human immunodeficiency virus 1] AAR31017.1; major capsid protein [Siphoviridae sp.] DAK31025.1 | **1,29E12** |
| 26 | C4LI71-DECOY | (?)LLFVFSK(?)<br>Query 2 LFVFSK 7<br>LFVFSK<br>Sbjct 186 LFVFSK 191 | site-specific DNA-methyltransferase [Corynebacterium diphtheriae] CAB0919846.1; gp89 [Corynebacterium phage P1201] YP_001468987.1; hypothetical protein PVA8_320 [Vibrio phage PVA8] URQ03306.1; long tail fiber proximal subunit [Escherichia phage moha] QHR68342.1 | **1,15E12** |
| 27 | C4LJ46-DECOY | (?)STNQAPAMTTVIMGPPK(?) | hypothetical protein [Bacteriophage sp.] UVX50895.1; pol protein, partial [Human immunodeficiency virus 1] ABJ92896.1; | 2012000 |
| 28 | C4LG68-DECOY | (?)GGNFETPAMGRLEAVMRTR(?) | integrase [Streptococcus phage Javan23] QBX15966.1; response regulator [Bacteriophage sp.] DAE37297.1 | 2256000 |
| 29 | C4LKQ7-DECOY | (?)IPALYAMASAAQIK(?) | putative MFS-type transporter [Staphylococcus phage UPMK_1] ATW69267.1; RNA dependent RNA polymerase [Bacteriophage sp.] DAQ44166.1 | 6568000 |
| 30 | C4LKL2-DECOY | (?)TLTDSVMDAVGK(?) | Replication associated protein [Bacteriophage sp.] DAR31281.1; Telomere resolvase [Bacteriophage sp.] UWG71171.1; | 6826000 |
| 31 | C4LLP7-DECOY | (?)FTTQYAQVLNADSEMLGR(?)<br>Query 3 TQYAQV 8<br>TQ AQ+<br>Sbjct 95 TQRAQI 100 | immunity-specific protein Beta241 [Corynephage beta]AAA32184.1;ADP-ribosyltransferase [Enterobacter phage fGh-Ecl01] USL85699.1; adenine-specific methyltransferase [Siphoviridae sp.] DAZ42774.1 | 6023000 |
| 32 | C4LJH0-DECOY | (?)GQGGNRKMGETDANPGDAGR(?) | GenBank: AFB75790.1 putative tail associated lysin [Lactococcus phage P335] ABI54246.1; capsid protein, partial [Norovirus Hu/GII.4/101228-a/BE-3540] AFY98775.1 | 1510000 |
| 33 | C4LGU3-DECOY | (?)IPADRR(?) | Major capsid protein [Bacteriophage sp.] DAU02489.1; polymerase PB2 [Influenza A virus (A/swine/Kentucky/15TOSU2007/2015(H3N2))] APO40360.1 | 1057000 |
| 34 | C4LKL5-DECOY | (?)LSSIEKR(?)<br>Query 2 SSIEK 6<br>SS EK<br>Sbjct 494 SSSEK 498 | Chain A, DIPHTHERIA TOXIN [Corynephage beta] 1DDT_A putative DNA primase [Corynebacterium phage P1201] YP_001468956.1; Type I restriction enzyme [Bacteriophage sp.] DAX02722.1; envelope glycoprotein [Human immunodeficiency virus 1] AQL57320.1; | 1168000 |
| 35 | C4LI78-DECOY | (?)SVSVMTMVEQNK(?) | Tail tape measure [Siphoviridae sp.] DAH41247.1; putative membrane-associated HD superfamily hydrolase [Bacteriophage sp.] DAW98318.1 | 984500 |
| 36 | C4LJA9-DECOY | (?)NTNDEGKGQK(?) | envelope glycoprotein, partial [Human immunodeficiency virus 1] QCI54352.1; scaffolding protein [Mycobacterium phage Cosmo] AJD82101.1 | 1152000 |
| 37 | C4LI32-DECOY | (?)SELMQIDDVSSR(?) | major tail protein [Bacteriophage sp.] DAN09434.1, [Myoviridae sp.] DAY66811.1 and [Siphoviridae sp.] DAI47842.1 | 5363000 |
| 38 | C4LG38-DECOY | (?)DEGAAVTAAGWVATRSGL(?)<br>Query 3 GAAVTAAGW 11<br>G A TA W<br>Sbjct 268 GTA-TADSW 275 | immunity-specific protein Beta371 [Corynephage beta] AAA32187.1; gp39 [Corynebacterium phage BFK20] a lytic phage of the industrial producer Brevibacterium flavum YP_001456769.1; putative major capsid protein [Corynebacterium phage P1201] YP_001468934.1; DNA annealing helicase and endonuclease [Podoviridae sp.] DAH75467.1 | 1515000 |

| | | | | |
|---|---|---|---|---|
| 39 | C4LHJ8-DECOY | (?)QKKMNRSSK(?)<br>Query 4 MNRSS 8<br>M SS<br>Sbjct 50 MTPSS 54 | immunity-specific protein Beta286 [Corynephage beta] AAA32186.1 Head-to-tail connector protein, podovirus-type [uncultured Caudovirales phage] CAB4160580.1; major capsid protein [Bacteriophage sp.] DAD57134.1; outer capsid protein [Human rotavirus C] AKJ87193.1 | 3119000 |
| 40 | C4LG40-DECOY | (?)TDGNIGILFEAISHCEATMR(?)<br>Query 13 SHCEA 17<br>SHC+A<br>Sbjct 171 SHCQA 175 | immunity-specific protein Beta201 [Corynephage beta] AAA32185.1; immunity-specific protein Beta286 [Corynephage beta] AAA32186.1; minor tail protein [Bacteriophage sp.] DAM09760.1; ankyrin repeat domain containing protein [Mimivirus sp.] QZX43547.2 | 1087000 |
| 41 | C4LI99-DECOY | (?)GLIMGGHVR (?)<br>Query 3 IMGG 6<br>+MGG<br>Sbjct 41 VMGG 44 | immunity-specific protein Beta241 [Corynephage beta] AAA32184.1; holin family protein [Bacteriophage sp.] UWG87962.1; ankyrin repeat-containing protein [Acanthamoeba polyphaga mimivirus] AKI78856.1; hypothetical protein LM1A4_033 [Leuconostoc phage 1-A4] YP_009168363.1; Phage terminase-like protein, large subunit [uncultured Caudovirales phage] CAB4148792.1 | 121740 |
| 42 | C4LGA6-DECOY | (?)GDAAVLCSGAIGVCALAEDR(?)<br>Query 12 GVC----ALAED 19<br>GV ALA+D<br>Sbjct 131 GVRQAYDALAQD 142 | immunity-specific protein Beta241 [Corynephage beta] AAA32184.1; Lysin motif [Myoviridae sp.] DAV60424.1; LysM [Siphoviridae sp.] DAS82107.1; | 46890 |
| 43 | C4LH32-DECOY | (?)TGVLDELLVVMGSNSREDNLK (?)<br>Query 4 LDELLVVMGSNS 15<br>L EL V G NS<br>Sbjct 288 LSELKTVTGTNS 299 | diphtheria toxin, partial [Corynephage beta] ABU25232.1; distal tail protein, partial [Siphoviridae sp.] DAZ56325.1; viral ankyrin 1 [Microplitis mediator bracovirus] AUO16837.1 | 81090 |
| 44 | C4LH04-DECOY | (?) VAPAVIPGK(?)<br>Query 6 IPGK 9<br>IPGK<br>Sbjct 437 IPGK 440 | Chain A, DIPHTHERIA TOXIN [Corynephage beta] 1DDT_A ; tape measure protein [Gordonia phage GiKK] UJQ86184.1; minor tail protein [Gordonia phage Gibbles] QDK01983.1; baseplate J-like protein [Bacteriophage sp.] UVY17948.1 | 870500 |
| 45 | C4LKI5-DECOY | (?)EGTLEADSLIR (?) | Small terminase subunit [Escherichia phage 500465-2] YP_009949266.1; hypothetical protein I5G61_gp44 [Mycobacterium phage Quesadilla] YP_009949800.1 | 533200 |
| 46 | C4LHC4-DECOY | (?)GMSSMSSASHQARGIMMASR (?)<br>Query 3 SSMSSA 8<br>SS SA<br>Sbjct 188 SSVASA 193<br>Query 2 MSSMSSASHQA 12<br>MS M QA<br>Sbjct 14 MS-MTIPRRQA 23<br><br>Query 7 SASHQARGIMM 17<br>S S QA M+<br>Sbjct 199 SQSLQAINTML 209 | immunity-specific protein Beta371 [Corynephage beta] AAA32187.1; immunity-specific protein Beta286 [Corynephage beta] AAA32186.1; cysteine protease [Myoviridae sp.] DAX51520.1 ;gag protein [Human immunodeficiency virus 1] QDB67569.1; gag protein [Human immunodeficiency virus 1] ALP13002.1; nonstructural polyprotein, partial [Norovirus GI] QEN27469.1 | 285000 |
| 47 | C4LLJ1-DECOY | (?)VQALRLQSR (?) | hypothetical protein [Bacteriophage sp.] UVX37815.1; Replication initiator A family protein [Siphoviridae sp.] DAK72172.1 | 875300 |
| 48 | C4LK38-DECOY | (?)GAVRTMK (?) | minor tail protein [Mycobacterium phage Apex] QFP96313.1; minor tail protein [Siphoviridae sp.] DAZ59018.1 | 44560 |
| 49 | C4LJU4-DECOY | (?)RACEAGADR (?)<br>Query 5 AGAD 8<br>AGAD<br>Sbjct 28 AGAD 31 | diphtheria toxin, partial [Corynephage beta] ABU25232.1; Spore cortex-lytic enzyme, lytic transglycosylase [Bacteriophage sp.] DAN11553.1; Spore cortex-lytic enzyme, lytic transglycosylase [Siphoviridae sp.] DAK95423.1; putative tail fiber protein [Corynebacterium phage phi673] YP_009639698.1; pol protein, partial [Human immunodeficiency virus 1] AAV59707.1, ABF00707.1; | 464700 |
| 50 | C4LJM2-DECOY | (?)ANAANREPSAGVVCGMQFTDGSNSR (?)<br>Query 18 FTDGSNS 24<br>F +GS S<br>Sbjct 140 FAEGSSS 146 | Chain A, DIPHTHERIA TOXIN [Corynephage beta] 1DDT_A; DNA gyrase subunit A [Bacteriophage sp.] DAI54469.1<br>CPXV219 protein [Cowpox virus] SNB54336.1 | 32490 |
| 51 | C4LIE4-DECOY | (?)ENLGGGHSTVR (?) | C40 family peptidase [Lactobacillus phage vB_Lga_AB1] QRD99546.1 | 61870 |
| 52 | C4LLN9-DECOY | (?)QNHGPVKSK (?)<br>Query 6 VKSK 9<br>V SK<br>Sbjct 523 VNSK 526 | Chain A, DIPHTHERIA TOXIN [Corynephage beta] 1DDT_A ; polymerase PB1 [Influenza A virus] QKY65769.1<br>helix-turn-helix DNA-binding domain protein [Streptomyces phage Soshi] QEQ94697.1 | 79340 |
| 53 | C4LGX7-DECOY | (?)DRSEGVAYMCNSSDAIR (?)<br>Query 12 SSDAI 16<br>SSD I<br>Sbjct 505 SSDSI 509 | Chain A, DIPHTHERIA TOXIN [Corynephage beta] 1DDT_A ; putative platelet-binding protein, minor tail fiber protein [Streptococcus phage 315.5] NP_795646.1 | 19860 |
| 54 | C4LIW4-DECOY | (?)AEILIAGKVVPLGMAILVSVAGLLIGFAAKGGAG SAKAWR (?)<br>Query 15 AI-LVSVAGLL--IGF 27<br>AI LV G L IGF<br>Sbjct 343 AIPLV---GELVDIGF 355<br><br>Query 1 IGFAA 5<br>IGFAA<br>Sbjct 353 IGFAA 357 | Chain A, DIPHTHERIA TOXIN [Corynephage beta] 1DDT_A; immunity-specific protein Beta241 [Corynephage beta] AAA32184.1; 3-hydroxybutyrate dehydrogenase [Acinetobacter phage MD-2021a CAH1085039.1; tail length tape measure protein [Pseudomonas phage Fc22] AYD80410.1 | 1397 |
| 55 | C4LID6-DECOY | (?)TRSLRLKAAEGMTK (?) | virion structural protein [Pseudomonas phage PA1C] QBX32339.1; head to tail connecting protein [Bacteriophage sp.] DAP98651.1 | 804900 |
| 56 | C4LIK4-DECOY | (?)DPASHNVIAIVTVFYFSHLAARGLIVAEVDKRML R (?) | hypothetical protein ACQ86_gp12 [Shigella phage SfMu] hypothetical protein ACQ86_gp12 [Shigella phage SfMu] YP_009152198.1; putative DNA polymerase I [Aeromonas phage LAh_7] YP_009998311.1 | 233,8 |
| 57 | C4LG92-DECOY | (?)YAGEGPDRADPGSFMGSDENR (?)<br>Query 12 GSFMG 16<br>GS MG<br>Sbjct 311 GSVMG 315 | Chain A, DIPHTHERIA TOXIN [Corynephage beta] 1DDT_A; hypothetical protein CPD7_23 [Clostridium phage CPD7] AZF89491.1 | 34970 |
| 58 | C4LH93-DECOY | (?)NAREQDLHSEMGFTNMDIGLLPTMLIRPLVAEL AAAGLQAGEGTGVRAAYDNAC(?) | immunity-specific protein Beta371 [Corynephage beta] AAA32187.1; hypothetical protein [Burkholderia phage FLC9] BDD79604.1; hypothetical protein [Yersinia phage YeP4] QCW23337.1 | 201,9 |
| 59 | C4LJ07-DECOY | (?)ESPVMDGEYAACKGEAR(?) | gp11 [Mycobacterium phage Brujita] YP_002241994.1; beta galactosidase small chain, partial [Bacteriophage sp.] UVY69219.1 | 61220 |
| 60 | C4LJR8-DECOY | (?)ERGNSMVDCTHEKQIK (?)<br>Query 12 EKQIK 16<br>EK IK<br>Sbjct 149 EKFIK 153 | diphtheria toxin, partial [Corynephage beta]ABU25232.1; Putative ribonuclease 3 [Satyrvirus sp.] AYV85794.1; Repressor protein CI [Siphoviridae sp.] DAM75527.1; gag protein [Human immunodeficiency virus 1] QDB67449.1 | 46070 |
| 61 | C4LGL4-DECOY | (?)GADIDTHTPSSGR (?)<br>Query 8 TPSS 11<br>TPSS<br>Sbjct 51 TPSS 54<br>Query 6 THTPSSGR 13<br>TH GR<br>Sbjct 448 THISVNGR 455 | immunity-specific protein Beta286 [Corynephage beta] AAA32186.1; Chain A, DIPHTHERIA TOXIN [Corynephage beta] 1DDT_A DNA polymerase I [Bacteriophage sp.] DAT67538.1; tail length tape measure protein [Streptococcus phage Javan617] QBX21901.1 | 302000 |
| 62 | C4LGM6-DECOY | (?)IEALDIPQR (?) | hypothetical protein UFOVP473_10 [uncultured Caudovirales phage] CAB4145518.1; ferrochetalase domain protein [Synechococcus phage S-P4] YP_009816077.1; cation:proton antiporter [Corynebacterium diphtheriae] WP_014307163.1 | 187900 |
| 63 | C4LJF1-DECOY | (?)GELASNVLDIGR (?) | hypothetical protein [Staphylococcus phage vB_SsapH-Golestan101-M] BCG66298.1; TPA: tail sheath protein [Bacteriophage sp.] DAH67593.1 | 424400 |
| 64 | C4LKU4-DECOY | (?)EGGFRDETGDGSSK (?) | baseplate hub + tail lysozyme [Prochlorococcus phage P-HM2] YP_004323394.1; tail fiber protein [Staphylococcus phage vB_SauS_IMEP5] ANM47011.1; Putative HNHc nuclease [Siphoviridae sp.] DAX94110.1 | 418100 |
| 65 | C4LGP9-DECOY | (?)GPGIITR (?) | putative internal virion protein [uncultured Mediterranean phage uvMED] BAQ86744.1; hypothetical protein DAC16_256 [Bacteroides phage DAC16] QIG63485.1; gp120, partial [Human immunodeficiency virus 1] CAD23653.1 | 494400 |
| 66 | C4LGW3-DECOY | (?)GAVLVEHK (? | Protein of unknown function (DUF2680) [Bacteriophage sp.] DAI11773.1; thymidylate synthase [Salmonella phage STWB21] QTJ63316.1; tail tape measure protein [Faecalibacterium phage FP_Epona] YP_009797164.1 | 764900 |
| 67 | C4LHA3-DECOY | (?)SISVPMTDSTDR (?)<br>Query 2 ISVP 5<br>ISVP<br>Sbjct 112 ISVP 115 | immunity-specific protein Beta201 [Corynephage beta] AAA32185.1; putative large terminase subunit [Gordonia phage GRU3] YP_009276095.1; pol protein, partial [Human immunodeficiency virus 1] AGF32915.1; | 47580 |
| 68 | C4LJ19-DECOY | (?)RCTPSQTDASPAGDSVIDK (?)<br>Query 8 DASPAGDSV 16<br>DA AG SV<br>Sbjct 61 DA--AGYSV 67 | Chain A, DIPHTHERIA TOXIN [Corynephage beta] 1DDT_A ; hypothetical protein [Siphoviridae sp.] DAT42697.1; Protein of unknown function (DUF935) [Bacteriophage sp.] UWH97483.1 | 86400 |
| 69 | C4LHJ6-DECOY | (?)DDDFMVLVDR (?) | hypothetical protein AXJ17_gp48 [Lactobacillus phage LfeSau] YP_009223148.1; holin [Bacteriophage sp.] DAJ24026.1; hypothetical protein IPP12_00004 [Streptococcus phage IPP12] APD21812.1 | 11900 |
| 70 | C4LJ63-DECOY | (?)GDSSQMRNTDGAR (?) | hypothetical protein DIDNDMLP_00331 [Klebsiella phage KP13-7] UYL05316.1; oxidoreductase [Vibrio phage MZH0603] QGT52232.1; polyprotein, partial [Rhinovirus B] ALT20857.1 | 111741 |
| 71 | Q8WXH0-6-DECOY | (?)MCSWRAPCCDSPIPLMVMSSGSPMESRSAESIAV SR (?)<br>Query 10 DSPIPLMVMSS 20<br>D PI M SS<br>Sbjct 44 DAPITRMTPSS 54 | immunity-specific protein Beta286 [Corynephage beta]AAA32186.1; immunity-specific protein Beta371 [Corynephage beta] AAA32187.1; Chain A, DIPHTHERIA TOXIN [Corynephage beta] 1DTP_A; RNA chaperone [Aeromonas phage BUCT695] UIW10504.1; hypothetical protein UFOVP731_20 [uncultured Caudovirales phage] CAB4160882.1; terminase [Gordonia phage RobinSparkles] AXH46463.1; hypothetical protein L3Y22_gp141 [Gordonia phage Octobien14] YP_010246342.1; stabilization protein [Podoviridae sp.] DAK85409.1 | 38410 |
| 72 | H0YB79-DECOY | (?)SGDQTRGTGPDCEWQFCSMVWSMPWR(?)<br>Query 11 DCE------WQ 15<br>DC+ WQ<br>Sbjct 156 DCDLWPEYRWQ 166 | immunity-specific protein Beta201 [Corynephage beta] AAA32185.1; DNA polymerase [Corynebacterium diphtheriae] CAB0562474.1;putative lysin [Corynebacterium phage P1201] YP_001468954.1; hypothetical protein HOU90_gp082 [Lactobacillus phage Lpa804] YP_009818229.1; serine protease inhibitor-like SPI-1 protein [Monkeypox virus] QNI40044.1; DNA polymerase I [Siphoviridae sp.] DAS48459.1; | **2.62614E7** |
| 73 | Q13936-25-DECOY | (?)LGFEKR(?)<br>Query 2 GFEKR 6<br>GFEKR<br>Sbjct 156 GFEKR 160 | iron transport system exported solute-binding protein [Corynebacterium diphtheriae] CAB1034823.1; putative holin [Corynebacterium phage P1201] YP_001468955.1; DNA primase domain protein [Nonlabens phage P12024L] YP_006560423.1; Restriction_endonuclease_like domain containing protein [uncultured Caudovirales phage] CAB5220562.1; serine protease [Siphoviridae sp.] DAT18916.1; tail fiber [Bacillus phage vB_BanS-Thrax4] UUV46710.1 | **3.03223E7** |
| 74 | K4DI89-DECOY | (?)GQKDTDPLGATSDQRPNPENSRR(?) | immunity-specific protein Beta201 [Corynephage beta] AAA32185.1; tail protein [Alteromonas phage vB_AmaS-R9Y2] UNA02106.1; Terminase large subunit [Bacteriophage sp.] DAO64006.1 | 101332 |
| 75 | P27635-DECOY | (?)QLGALPQEK(?)<br>Query 2 LGALP 6<br>LGALP<br>Sbjct 139 LGALP 143 | immunity-specific protein Beta286 [Corynephage beta] AAA32186.1; replicative DNA helicase [Pseudomonas phage vB_Pae_CF23a] QBI79904.1; Integrase [Bacteriophage sp.] DAS11593.1; MC091L [Molluscum contagiosum virus] QHW16479.1; polymerase [Hepatitis B virus] BAO31728.1 | 611800 |
| 76 | Q16720-DECOY | (?)QNDNGSSCER(?) | terminase DNA packaging enzyme large subunit [Acinetobacter phage SH-Ab 15599] AXF41497.1; envelope glycoprotein, partial [Human immunodeficiency virus 1] AWY93263.1; membrane glycoprotein UL119 [Human betaherpesvirus 6A] APO39220.1 | 2914000 |
| 77 | Q96AC1-DECOY | (?)FKYTHGSTGR(?) | virion morphogenesis family protein [Bacteriophage sp.] UVY37676.1; tail component [Phage sp. ctvTz5] DAE21567.1 | 3581000 |

| | | | | |
|---|---|---|---|---|
| 78 | P42285-DECOY | (?)RRGDVATGSGGEAR (?)<br>Query 2 RGDVATGSGGE 12<br>R + TGSGGE<br>Sbjct 56 RNKISTGSGGE 66 | SpaH/EbpB family LPXTG-anchored major pilin [Corynebacterium diphtheriae] WP_088251462.1; putative DNA primase [Corynebacterium phage P1201] YP_001468956.1; hypothetical protein [Klebsiella phage pKp383] UVD41553.1; head to tail adaptor [Myoviridae sp.] DAR38668.1; Phosphoribosylformylglycinamidine synthase [Acinetobacter phage MD-2021a] CAH1079775.1; helicase-primase subunit BBLF2/3 [Human gammaherpesvirus 4] ATO60069.1 | **1.33528E7** |
| 79 | V9GZ50-DECOY | (?)AMEAFPPEDIR (?)<br>Query 2 MEAF-PPEDIR 11<br>M AF P EDIR<br>Sbjct 68 MTAFGPQEDIR 78 | tRNA dihydrouridine synthase DusB [Corynebacterium diphtheriae] CAB0959088.1; putative portal [Corynebacterium phage P1201] YP_001468930.1; catalase HPII [Acinetobacter phage MD-2021a] CAH1069987.1; portal vertex protein [Bacteriophage sp.] DAR31282.1; pol protein, partial [Human immunodeficiency virus 1] QDB72360.1 | **1.015E7** |
| 80 | Q8IX12-DECOY | (?)CHGAPNAMDYFADSK(?)<br>Query 12 ADSK 15<br>ADSK<br>Sbjct 208 ADSK 211 | immunity-specific protein Beta371 [Corynephage beta] AAA32187.1; endolysin [Vibrio phage phiKT1028] UVD32192.1; DUF2479 domain-containing protein [Lactococcus phage bIL285] NP_076627.1; bi-component leukocidin LukMF' subunit M [Staphylococcus prophage phiPV83] NP_061649.1 | 702900 |
| 81 | O95169-DECOY | (?)NLLPALPGGK(?)<br>Query 2 LLPALPG 8<br>LLP PG<br>Sbjct 433 LLPTIPG 439 | Chain A, DIPHTHERIA TOXIN [Corynephage beta] DDT_A ; head-to-tail adaptor [Mycobacterium phage Donny] QBI96477.1; polyprotein precursor [Hepatitis C virus subtype 6o] ABP88846.1; | 1334000 |
| 82 | P05067-2-DECOY | (?)DGHGDLGSADMQR (?) | hypothetical protein [Bacteriophage sp.] DAQ96413.1; hypothetical protein ORF022 [Pseudomonas phage PA11] YP_001294615.1; hypothetical protein SEA_ANDRIS_60 [Streptomyces phage Andris] WAB08764.1 | 8.324,40 |
| 83 | A8MX34-DECOY | (?)SNLRRSNMYENQLPVDSSCPTK (?)<br>Query 15 VDSS 18<br>VDSS<br>Sbjct 6 VDSS 9 | Chain A, DIPHTHERIA TOXIN [Corynephage beta] 1DDT_A ; Integrase [Bacteriophage sp.] DAI95817.1; putative DNA primase [Brevibacillus phage Sundance] YP_009194183.1 | 780600 |
| 84 | Q8WXH0-2-DECOY | (?)TGGGTPAALSQEEDNSTPMFAASAGR(?)<br>Query 5 TPAALS 10<br>T AALS<br>Sbjct 300 TTAALS 305 | diphtheria toxin, partial [Corynephage beta]BAB03348.1; putative scaffolding protein [Roseobacter phage CRP-7] QBQ72849.1; ORF1a polyprotein, partial [Severe acute respiratory syndrome coronavirus 2] UNO74509.1 | 1164000 |
| 85 | Q9H078-DECOY | (?)DDGALTPTLK(?)<br>Query 1 DDGALTPTL 9<br>DDGAL TL<br>Sbjct 179 DDGALS-TL 186 | immunity-specific protein Beta241 [Corynephage beta] AAA32184.1; DNA replication/repair protein RecF [Corynebacterium diphtheriae] RNF45856.1; putative tail measure protein [Corynebacterium phage P1201] YP_001468947.1; N-6 DNA Methylase [Acinetobacter phage MD-2021a] CAH1090743.1; major capsid protein [Siphoviridae sp.] DAP09927.1; GDSL-like Lipase/Acylhydrolase [Siphoviridae sp.] DAQ98621.1 | **1.38893E7** |
| 86 | K7EJZ9-DECOY | (?)ELGVGGEEIR(?)<br>Query 1 ELG----------VGGEE 8<br>ELG VG EE<br>Sbjct 105 ELGLSLTEPLMEQVGTEE 122 | Chain A, DIPHTHERIA TOXIN [Corynephage beta] 1DDT_A ; active helicase ring shaped helicase [Siphoviridae sp.] DAW75510.1; RNA polymerase-ADP-ribosyltransferase Alt [Vibrio phage phi-Grn1] ALP46977.1; ParB-like nuclease domain [Bacteriophage sp.] UVY12958.1 | 474600 |
| 87 | Q96PU5-7-DECOY | (?)GADLRCGGLASR(?)<br>Query 7 GGLAS 11<br>GG AS<br>Sbjct 43 GGIAS 47 | immunity-specific protein Beta241 [Corynephage beta] AA32184.1; hypothetical protein UFOVP626_20 [uncultured Caudovirales phage] CAB4153688.1; pentapeptide repeat protein [Bacteriophage sp.] DAW94266.1; pentapeptide repeat protein [Siphoviridae sp.] DAN36313.1; lysozyme [Siphoviridae sp. ct86u1] DAF58635.1 | 1617860 |
| 88 | P13671-DECOY | (?)KPLVVTK(?) | major capsid protein [Bacteriophage sp.] DAW74662.1; hypothetical protein [uncultured Mediterranean phage uvMED] BAQ84573.1; putative tail protein 4 [Salmonella phage S144] QMV47845.1 | 54020 |
| 89 | B4DXL1-DECOY | (?)GEAVEIAPR(?) | endolysin [Caudovirales sp. ctrNG92] DAF49189.1; hypothetical protein SS1_02 [Cronobacter phage vB_CsaP_Ss1] AIK67506.1 | 99240 |
| 90 | Q9P2N6-6-DECOY | (?)VLEIMMHK(?) | envelope glycoprotein [Human immunodeficiency virus 1] AGC81117.1; MC133 [Molluscum contagiosum virus subtype 1] AYO88805.1; tail tubular protein [Bacteriophage sp.] UVN00301.1 | 1097000 |
| 91 | Q8NDM7-2-DECOY | (?)MITEIEK(?) | hypothetical protein SPFM1_00111 [Salmonella phage SPFM1] VFR10297.1; ankyrin repeat protein [Fadolivirus 2] QKF94907.1 | 1643000 |
| 92 | O15541-DECOY | (?)ENCLTPK(?) | homing endonuclease [Bacteriophage sp.] DAJ12705.1; Hemolysin [Bacteriophage sp.] DAX10491.1; hemolysin [Siphoviridae sp.] DAZ61289.1 | 1811000 |
| 93 | H0Y872-DECOY | (?)NPSEVGLK(?)<br>Query 4 EVGLK 8<br>+VGLK<br>Sbjct 245 DVGLK 249 | immunity-specific protein Beta371 [Corynephage beta] AAA32187.1; immunity-specific protein Beta201 [Corynephage beta] AAA32185.1; envelope glycoprotein [Human immunodeficiency virus 1] ADG57029.1; LptA [Bacteriophage sp.] DAN84629.1; large polymerase protein [Measles morbillivirus] UTJ94268.1 | 1382000 |
| 94 | P78415-DECOY | (?)LAGAVVIR(?)<br>Query 2 AGAVV 6<br>AG VV<br>Sbjct 77 AGGVV 81 | Chain A, DIPHTHERIA TOXIN [Corynephage beta] 1DDT_A ; NeuD/PglB/VioB family sugar acetyltransferase protein [Bacillus phage PK1] UKL29997.1; gp120, partial [Human immunodeficiency virus 1] AAC55414.1; tapemeasure protein [Gordonia phage Untouchable] P_009853691.1 | 4768000 |
| 95 | Q96RD6-DECOY | (?)NWKSPKGPGSMMASPGADERDMECR(?)<br>Query 9 GSMMASPG-AD 18<br>GS M G AD<br>Sbjct 311 GSVM---GIAD 318 | Chain A, DIPHTHERIA TOXIN [Corynephage beta] 1DDT_A; immunity-specific protein Beta241 [Corynephage beta] AAA32184.1; leucyl-tRNA synthetase [Acinetobacter phage MD-2021a] CAH1089691.1; putative nuclease [uncultured Mediterranean phage uvMED] BAR30589.1 | 22370 |
| 96 | O15375-DECOY | (?)QLTECVLKCMECDAKR(?) | 4Fe-4S dicluster domain protein [Bacteriophage sp.] UWI17486.1; Polyferredoxin [Siphoviridae sp.] DAX98805.1; pol protein, partial [Human immunodeficiency virus 1] AEN04404.1 | 162400 |
| 97 | Q9UPN3-4-DECOY | (?)DMTQMFCEEPKGKSPANTDWT QGMLGK(?) | Replication associated protein [Microviridae sp.] DAP34873.1; hypothetical protein JO84_gp126 [Aureococcus anophagefferens virus] YP_009052423.1 | 29910 |
| 98 | C9JGX5-DECOY | (?)RQQLGHMQEHEK(?) | holin gp22 [Burkholderia phage KS9] YP_003090198.1; holin [Myoviridae sp.] DAF79651.1; holin family protein [Escherichia phage ESSI2_ev040] YP_009877496.1; holin [Yersinia phage vB_YenM_324] UKL54209.1; Bacteriophage lambda, GpH, tail tape measure, C-terminal [uncultured Caudovirales phage] CAB4136554.1 | 497200 |
| 99 | F8W840-DECOY | (?)SPLLSTR(?) | hypothetical protein [Bacteriophage sp.] DAR58743.1; Uncharacterised protein [Acinetobacter phage MD-2021a] CAH1091257.1; tail protein [Siphoviridae sp.] DAI44530.1 | 1600000 |
| 100 | H0YBZ5-DECOY | (?)MSKCSLNQECTCSVVEDDQMQRYK(?)<br>Query 7 NQECTCSVV 15<br>NQ VV<br>Sbjct 185 NQ----TVV 189 | immunity-specific protein Beta241 [Corynephage beta] AAA32184.1; prohead core scaffold and protease [Prochlorococcus phage P-SSM4] YP_214667.1; hypothetical protein SPFM8_00010 [Salmonella phage SPFM8] VFR11642.1 | 18840 |
| 101 | Q9Y493-2-DECOY | (?)GCSSAEHSLGLAEQR(?)<br>Query 8 SLGLAE 13<br>SL AE<br>Sbjct 137 SLPFAE 142<br><br>Query 12 AEQR 15<br>AEQR<br>Sbjct 175 AEQR 178<br><br>Query 11 LAEQR 15<br>LA QR<br>Sbjct 93 LATQR 97 | Chain A, DIPHTHERIA TOXIN [Corynephage beta] 1DDT_A ; immunity-specific protein Beta201 [Corynephage beta] AAA32185.1 ; immunity-specific protein Beta241 [Corynephage beta] AAA32184.1; immunity-specific protein Beta286 [Corynephage beta] AAA32186.1; oligopeptide ABC transporter ATP-binding protein [Acinetobacter phage MD-2021a] CAH1092059.1; capsid and scaffold protein [Pseudomonas phage VSW-3] YP_009596178.1; phage tail tape measure protein [Enterococcus phage Entfac.YE1] BCR37048.1 | 2700000 |
| 102 | J3QQX5-DECOY | (?)VEISKCAPLK(?) | endoribonuclease [Bacteriophage sp.] DAT26342.1; Integrase [Bacteriophage sp.] DAG76052.1 | 1455000 |
| 103 | Q9GZX6-DECOY | (?)TASESGLEDKLK(?) | DNA-polymerase catalytic subunit [Staphylococcus phage vB_StaM_SA1] QPI17093.1; ABC transporter [Bacteriophage sp.] UWI29106.1 | 304200 |
| 104 | P27037-DECOY | (?) SMNGSMQMTKIPRK(?)<br>Query 3 NGSM 6<br>NG M<br>Sbjct 52 NGAM 55<br><br>Query 2 MNGSMQ 7<br>M+ M+<br>Sbjct 27 MDNNME 32 | immunity-specific protein Beta241 [Corynephage beta] AAA32184.1; immunity-specific protein Beta286 [Corynephage beta] AAA32186.1; tape measure protein [Streptococcus phage SW19] AYP29683.1; terminase large subunit [Lactococcus phage P118] YP_009035824.1 | 1810000 |
| 105 | Q96NY9-DECOY | (?)NNLLPGHSK(?) | tail length tape measure protein [Escherichia phage vB_KppS-Ant] CAD5235972.1; hypothetical protein HSE3_gp042 [Klebsiella phage vB_KleS-HSE3] QIN94990.1 | 166100 |
| 106 | Q6P5Z2-DECOY | (?)QFSFAMIYEDARK(?) | terminase family protein [Escherichia phage ST0] YP_009608498.1; minor capsid protein [Bacteriophage sp.] DAH88091.1 | 496300 |
| 107 | 9BTW9-DECOY | (?)QAGLGLNFGSPG(?) | UvrD-like DNA helicase, C-terminal [uncultured Caudovirales phage] CAB4165171.1; putative internal viron protein [Pelagibacter phage Skadi-5 EXVC105P] UWJ03751.1 | 85410 |
| 108 | Q8NGC2-DECOY | (?)LDSSLPGSR(?) | hypothetical protein HWB33_gp25 [Leptospira phage LE3] YP_009835417.1 ; envelope glycoprotein, partial [Human immunodeficiency virus 1] AIY68095.1; | 9680880 |
| 109 | Q6WBX8-1-DECOY | (?)AQKKPAK(?) | Cell wall hydrolase autolysin [Siphoviridae sp.] DAZ28878.1; N-acetylmuramoyl-L-alanine amidase [Bacteriophage sp.] UVX76119.1; large terminase [Siphoviridae sp.] DAN00483.1; peptidoglycan hydrolase [Siphoviridae sp.] DAJ77011.1 | 119500 |
| 110 | Q13360-DECOY | (?)LSFTAEVK(?)<br>Query 2 SFTAEV 7<br>SFTA++<br>Sbjct 323 SFTADI 328 | immunity-specific protein Beta371 [Corynephage beta] AAA32187.1; hypothetical protein [Burkholderia phage FLC9] BDD79536.1; ATP-dependent DNA ligase [Enterobacter phage phiEap-1] YP_009196347.1 | 2850000 |
| 111 | Q13216-DECOY | (?)GLLSIWEESR(?)<br>Query 3 LSIWEE 8<br>LS WE+<br>Sbjct 3 LSQWEQ 8 | immunity-specific protein Beta286 [Corynephage beta] AAA32186.1; minor tail protein [Bacteriophage sp.] UWI24341.1; envelope glycoprotein [Human immunodeficiency virus 1] ADF83990.1 | 7046000 |
| 111 | Q13216-DECOY | (?)GLLSIWEESR(?)<br>Query 3 LSIWEE 8<br>LS WE+<br>Sbjct 3 LSQWEQ 8 | immunity-specific protein Beta286 [Corynephage beta] AAA32186.1; minor tail protein [Bacteriophage sp.] UWI24341.1; envelope glycoprotein [Human immunodeficiency virus 1] ADF83990.1 | 7046000 |
| 112 | Q9H3S7-DECOY | (?)ETCTNASKR(?) | antirepressor protein [Bacteriophage sp.] DAU81194.1; envelope glycoprotein [Human immunodeficiency virus 1] UST15839.1; envelope glycoprotein [Human immunodeficiency virus 1] AXP18759.1; envelope glycoprotein [Human immunodeficiency virus 1] AAC40596.1 | 672900 |
| 113 | C4LHQ5-DECOY | (?)DNMLWPVGNEVCVCEPGQPEFANASSR(?)<br>Query 21 FANASS 26<br>FA SS<br>Sbjct 140 FAEGSS 145 | Chain A, DIPHTHERIA TOXIN [Corynephage beta] 1DDT_A ; lysin A [Gordonia phage Schmidt] YP_010050961.1 tail terminator [Arthrobacter phage Lucy] AOQ28284.1; hypothetical protein CM02_gp058 [Mycobacterium phage Gadjet] YP_009011289.1; putative plasmid partitioning and stability protein StbB [Acinetobacter phage MD-2021a] CAH1090786.1 | 324210 |
| 114 | C4LLQ2-DECOY | (?)SLESAAR (?) | Tail tape measure [Myoviridae sp.] DAK10548.1; tail tape measure [Siphoviridae sp. ctCUc43] DAE19141.1; minor capsid protein [Siphoviridae sp.] DAN72296.1 | 608926 |
| 115 | C4LL96-DECOY | (?)RPMSGR(?) | tail tape measure [Siphoviridae sp.] DAU85324.1; tail protein [Bacteriophage sp.] DAG24887.1 | 39003,4 |
| 116 | C4LIM9-DECOY | (?)AQTNGGR(?) | DNA polymerase [Bacillus phage JBP901]; YP_009149176.1; Cpl 1 lysin [Siphoviridae sp. ctC6Q17]; DAE25693.1 | 19989 |
| 117 | C4LGP0-DECOY | (?)NAAAGSGR(?)<br>Query 2 AAAG 5<br>AAAG<br>Sbjct 122 AAAG 125 | immunity-specific protein Beta241 [Corynephage beta] AAA32184.1 ; structural protein [Pseudomonas phage AN14] ANO57389.1 ,envelope glycoprotein B [Human alphaherpesvirus 3] AKE13368.1; | 13124,4 |
| 118 | C4LGQ5-DECOY | (?)EVALGVR (?)<br>Query 2 VALG 5<br>VALG<br>Sbjct 116 VALG 119 | immunity-specific protein Beta286 [Corynephage beta] AAA32186.1; tail fibers protein [Klebsiella phage Kp_Pokalde_001] QWT56636.1; large terminase [Siphoviridae sp.] DAJ71453.1 | 657077 |
| 119 | C4LHJ0-DECOY | (?)AICEIAR(?)<br>Query 2 ICEI 5<br>+CEI<br>Sbjct 132 VCEI 135 | immunity-specific protein Beta286 [Corynephage beta] AAA32186.1; repressor protein [Siphoviridae sp.] DAT48034.1; envelope glycoprotein [Human immunodeficiency virus 1] AQL57337.1, DNA polymerase family B [Bacteriophage sp.] UVX61673.1 | 1709920 |

| | | | | |
|---|---|---|---|---|
| 120 | C4LJE9-DECOY | (?)LAPAVISK (?) | tail completion protein [Escherichia phage vB_EcoS-569R6] URC09581.1; Tape measure domain protein [Siphoviridae sp.]; DAQ02495.1 | 4767950 |
| 121 | C4LGZ4-DECOY | (?)KVSGGLHK(?) | gyrase subunit B [Acinetobacter phage MD-2021a] CAH1093311.1; DNA TOPOISOMERASE IV, B SUBUNIT [Siphoviridae sp.] DAL56275.1 | 832793 |
| 122 | C4LGW3-DECOY | (?)GAVLVEHK(?) | thymidylate synthase [Salmonella phage STWB21] QTJ63316.1; lysozyme [Escherichia phage Ecwhy_1] QAY00471.1 | 764924 |
| 123 | C4LLM7-DECOY | (?)LAGPVASIR (?)<br>Query 4 PVASI 8<br>PV SI<br>Sbjct 136 PVTSI 140<br><br>Query 2 AGPV 5<br>AGPV<br>Sbjct 280 AGPV 283 | immunity-specific protein Beta371 [Corynephage beta] AAA32187.1; immunity-specific protein Beta286 [Corynephage beta] AAA32186.1; head-to-tail adaptor [Mycobacterium phage prophi88-1] QSM02800.1; gag protein, partial [Human immunodeficiency virus 1] ASV70837.1; | 430329 |
| 124 | C4LIT9-DECOY | (?)AKGVGSGTSR(?) | Major capsid protein [Siphoviridae sp.] DAU00215.1; virion RNA polymerase [Vibrio phage vB_VspP_pVa5] YP_009876144.1; Chromatin remodeling complex ATPase [Siphoviridae sp.] DAG55609.1 | 644664 |
| 125 | C4LGX3-DECOY | (?)AAKEAGDRR (?)<br>Query 1 AAKEA 5<br>AA EA<br>Sbjct 123 AAGEA 127<br><br>Query 1 AAKEA 5<br>AA EA<br>Sbjct 109 AALEA 113 | immunity-specific protein Beta241 [Corynephage beta] AAA32184.1; PolyVal ADP-Ribosyltransferase [Myoviridae sp.] DAR46956.1; ATPase [Podoviridae sp.] DAS28226.1; | 644731 |
| 126 | C4LHE6-DECOY | (?)LGGSLSTDVR(?)<br>Query 2 GGSLS 6<br>GGSLS<br>Sbjct 221 GGSLS 225<br><br>Query 1 LGGSL 5<br>LG SL<br>Sbjct 106 LGLSL 110 | Diphtheria toxin homolog CRM228 fragment A P00589.2 ; Chain A, DIPHTHERIA TOXIN [Corynephage beta] 1DDT_A ; peptidase G2 [Staphylococcus phage 37] YP_240092.1; hypothetical protein Unbinned3138contig1002_36 [Prokaryotic dsDNA virus sp.] QDP52158.1; bacterial toxin [Siphoviridae sp.] DAW74264.1 | 1004,5372 |
| 127 | C4LLU3-DECOY | (?)LASTPVSPTAR(?)<br>Query 2 ASTP 5<br>ASTP<br>Sbjct 57 ASTP 60 | immunity-specific protein Beta241 [Corynephage beta] AAA32184.1 head-tail preconnector protease C / scaffolding domain Nu3 [Vibrio phage HY01] QJT70744.1; [Molluscum contagiosum virus subtype 1] NP_044065.1; envelope glycoprotein gB [Human gammaherpesvirus 4] AHA36404.1 | 679494 |
| 128 | C4LG91-DECOY | (?)VLVGGTEPEGR(?) | bacterial toxin [Siphoviridae sp.] DAI52742.1; phage related lysozyme [Acidithiobacillus phage AcaML1] AFU62909.1; DNA methylase [Gordonia phage Leonard] YP_010002224.1; | 538396 |
| 129 | C4LH59-DECOY | (?)AEMEVLDSTPR(?) | HNH endonuclease [Klebsiella phage JD001] YP_007392863.1; tail tube initiator protein [Escherichia phage PO103-1] UTQ80263.1; | 182151 |
| 130 | C4LHA3-DECOY | (?)SISVPMTDSTDR (?)<br>Query 2 ISVP 5<br>ISVP<br>Sbjct 112 ISVP 115 | immunity-specific protein Beta201 [Corynephage beta] AAA32185.1; putative large terminase subunit [Gordonia phage GRU3] YP_009276095.1; Putative ATP dependent Clp protease [Siphoviridae sp.] DAE93020.1; L2 protein [Human papillomavirus 174] CCV02863.1; pol protein, partial [Human immunodeficiency virus 1] AGF32915.1; | 47577,1 |
| 131 | C4LIX0-DECOY | (?)GFDDWDPTAPEYDSQE(?)<br>Query 1 GFDDW 5<br>GF DW<br>Sbjct 108 GFGDW 112 | immunity-specific protein Beta371 [Corynephage beta] AAA32187.1; major capsid protein [Arthrobacter phage KeAlii] UDL14613.1; portal protein [Bacteriophage sp.] UWG67114.1; AAA ATPase [Pseudomonas phage MiCath] WAX22444.1; putative DNA replication protein [Streptococcus satellite phage Javan462] QBX10613.1; polyprotein [Hepatitis C virus subtype 6m] WAX22444.1 | 120,332 |
| 132 | C4LIK1-DECOY | (?)SSMSPLYPCAMDTPS (?) | Thymidylate synthase [Siphoviridae sp.] DAI31909.1; putative queuine tRNA-ribosyltransferase [Pantoea phage vB_PagS_MED16] AZS06278.1; hypothetical protein Unbinned6284contig1001_19 [Prokaryotic dsDNA virus sp.] QDP50463.1 | 6251020 |
| 133 | C4LHH1-DECOY | (?)TELHNSEVLTDAHTHVSIATQATMPERITVDYSVPKLEAR(?)<br>Query 9 TDAHTHV 15<br>TD H+<br>Sbjct 48 TDGYVHI 54 | immunity-specific protein Beta241 [Corynephage beta] AAA32184.1; immunity-specific protein Beta371 [Corynephage beta] AAA32187.1; tail sheath protein [Siphoviridae sp. ctsf32] DAD71446.1; glyoxylase family protein [Escherichia phage Mansfield] QEG09891.1; RusA-like Holliday junction resolvase [Propionibacterium phage PFR1] YP_009287723.1 | 2979,37 |
| 134 | C4LIX0-DECOY | (?)FEVSNLWADENAPVLNFNVMVAYRMRAGHSRVYGFHTIR(?) | | 120,332 |
| 135 | P16930-2-DECOY | GIPMTRR | major capsid protein E [Bacteriophage sp.] UVY27154.1; major capsid protein [Caudovirales sp.] DAH50489.1; portal protein [Bacteriophage sp.] DAQ28708.1 | **1,91E+12** |
| 136 | P50851-DECOY | (?)LFLDYLLVQK(?)<br>Query 2 FLDYLLVQ 9<br>FL YLLVQ<br>Sbjct 350 FLIYLLVQ 357<br><br>Query 1 LFLD----YL----------LVQK 10<br>LFLD YL LVQK<br>Sbjct 181 LFLDEPTTYLDLATSVEILELVQK 204 | iron ABC transporter [Corynebacterium diphtheriae subsp. lausannense] OLN15681.1; MULTISPECIES: iron chelate uptake ABC transporter family permease subunit [unclassified Corynebacterium]WP_023025753.1; DNA polymerase B [Bacteriophage sp.] DAD57058.1; DNA polymerase B [Podoviridae sp.] DAG68826.1; portal protein [Myoviridae sp.] DAY09267.1; DNA polymerase I [Morganella phage MmP1] YP_002048647.1 | **1,72E+12** |
| 137 | H0YFK4-DECOY | (?)NQLNRDVNSSLLAISDFLLDAMGLQLVLEGLGDVRNDSEMLAAILGEVAK (?)<br>Query 5 RDVNSSLLAISDFLLD 20<br>R V SSL I LD<br>Sbjct 193 RSVGSSLSCIN---LD 205 | Chain A, DIPHTHERIA TOXIN [Corynephage beta] 1DDT_A; immunity-specific protein Beta286 [Corynephage beta] AAA32186.1; Diphtheria toxin homolog CRM228 P00589.2; immunity-specific protein Beta201 [Corynephage beta] AAA32185.1; immunity-specific protein Beta241 [Corynephage beta] AAA32184.1; immunity-specific protein Beta371 [Corynephage beta] AAA32187.1; putative tape measure protein [Corynebacterium phage phi674] YP_009639749.1; tail tape measure [Siphoviridae sp.] DAG85167.1; RIIA lysis inhibitor [Shewanella sp. phage 1/40] YP_009104011.1; terminase large subunit [Escherichia phage grams] YP_009901824.1 | **1,34E+12** |
| 138 | Q5T215-DECOY | (?)ALTLLVGCAMNEIQEPLFSRQDVVVEGGGVISLCEMPLLQLPLAEMGCQLPDGLYREK (?)<br>Query 12 LQLPLAE 18<br>L LP AE<br>Sbjct 136 LSLPFAE 142<br><br>Query 14 LPLAEMGCQLP 24<br>LP A G LP<br>Sbjct 427 LPIA--GVLLP 435<br><br>Query 19 MGCQLPDG 26<br>MG DG<br>Sbjct 314 MG--IADG 319 | Chain A, DIPHTHERIA TOXIN [Corynephage beta]1DDT_A; immunity-specific protein Beta371 [Corynephage beta] AAA32187.1; immunity-specific protein Beta286 [Corynephage beta]; AAA32186.1; ATP-dependent endonuclease [Corynebacterium diphtheriae] CAB0547077.1; lysis accessory protein Rz1 [Klebsiella phage vB_KpnS-MUC-5] UMW87868.1; repressor domain protein [Siphoviridae sp.] DAV94682.1; repressor domain protein [Bacteriophage sp.] DAV53220.1; Large Terminase [Siphoviridae sp.] DAL98001.1; RNA-dependent RNA polymerase, partial [Lihan tick virus] AYV61039.1 | **1,02E+12** |
| 139 | Q8N539-2-DECOY | (?)QNAAAPK (?) | endolysin; inhibits RNA polymerase [Ralstonia phage RSB3] YP_008853912.1; ORF1ab polyprotein, partial [Severe acute respiratory syndrome coronavirus 2] UNH99369.1; minor tail protein [Bacteriophage sp.] UVY05131.1 | 7.095,55 |
| 140 | Q9UPU7-DECOY | (?)DPPLMK (?) | DNA polymerase [Mycobacterium phage Myrna] YP_002225071.1; head to tail connecting protein [Bacteriophage sp.] DAJ09887.1; BVRF1 [Human gammaherpesvirus 4] AWG88704.1 | 42.713,10 |
| 141 | Q9H4A3-DECOY | (?)QAWAPK (?) | LysM-like endolysin [Gordonia phage Skog] YP_010059406.1; bacterial toxin [Siphoviridae sp.] DAF12906.1; Pr gag-pro [Human T-cell leukemia virus type I] NP_057861.1; gag/pro/pol precursor [Simian T-lymphotropic virus 1] QDF62685.1; | 7.061,64 |
| 142 | D3DPQ1-DECOY | (?)DPPPFK (?) | K15 [Human gammaherpesvirus 8] UQT64912.1; DNA ligase [Stenotrophomonas phage YB07] UMO77365.1 | 41.559,20 |
| 143 | Q4AC94-4-DECOY | (?)LLPTMLIRPLVAELAAKLAIEK(?)<br>Query 9 PLVAEL 14<br>PLV EL<br>Sbjct 345 PLVGEL 350<br><br>Query 1 LLPT 4<br>LLPT<br>Sbjct 433 LLPT 436 | Chain A, DIPHTHERIA TOXIN [Corynephage beta]1DDT_A; portal protein [Siphoviridae sp.] DAT31011.1 | 10428 |
| 144 | Q96RL7-DECOY | (?)GALSLIK(?)<br>Query 1 GALS 4<br>GALS<br>Sbjct 330 GALS 333 | diphtheria toxin, partial [Corynephage beta] ABU25232.1; putative recT protein [Streptococcus phage phi891591] AGF87579.1; Ribonucleotide reductase small subunit [Lymphocystis disease virus 1] NP_078636.1 | 25.013,80 |
| 145 | Q5STU3-DECOY | (?)KVVVEK(?) | nonstructural polyprotein [Norovirus GII] QVV57737.1 | 9.696,74 |
| 146 | B4DT16-DECOY | (?)VHKAAAK(?) | envelope glycoprotein, partial [Human immunodeficiency virus 1] ADG26538.1; helicase-primase primase subunit [Saimiriine alphaherpesvirus 1] YP_003933788.1 | 47.799,80 |
| 147 | E9PJG4-DECOY | (?)DSPIAPK(?) | pectin lyase fold/virulence factor [Vibrio phage 2.275.O._10N.286.54.E11] AUS02986.1; RNA ligase [Gordonia phage ChisanaKitsune] QZE10792.1 | 9.495,71 |
| 148 | E7EPZ0-DECOY | (?)SSASGPPK(?) | hypothetical protein LU11_gp039 [Pseudomonas phage Lu11]; phosphoprotein [Lyssavirus rabies] ALC74030.1 | 13.988,70 |
| 149 | Q03933-2-DECOY | (?)ENFPNK(?) | recombinase [Staphylococcus phage vB_SauM-V1SA22] UVD42777.1 envelope glycoprotein [Human immunodeficiency virus 1] AKN11060.1; | 4372880 |
| 150 | Q15878-3-DECOY | (?)LNFLKK(?) | Papain-like cysteine protease AvrRpt2 [Myoviridae sp.] DAR18547.1; replicase [Potato virus S] QYY48756.1 | 675619 |
| 151 | Q6PIV7-DECOY | (?)KTCGMPK(?) | phenylacetaldehyde dehydrogenase [Acinetobacter phage MD-2021a] CAH1090655.1; putative Zn-dependent protease [Siphoviridae sp.] DAI35237.1 | 434507 |
| 152 | Q13765-DECOY | (?)EPFMPK(?) | methionyl-tRNA synthetase [Siphoviridae sp. ctBLh2] DAF45288.1 ; X protein [Hepatitis B virus] AVK93882.1 | 363093 |
| 153 | P29122-6-DECOY | (?)VAEYGNK(?) | terminase large subunit [Synechococcus phage S-P4] YP_009816042.1; tail tape measure protein [Siphoviridae sp.] DAQ09619.1 | 226850 |
| 154 | C9JUK0-DECOY | (?)VRLGAHK(?) | putative baseplate assembly protein [Bacteriophage sp.] DAV87054.1; DNA polymerase I [Microbacterium phage SBlackberry] UAW08794.1 | 221001 |
| 155 | Q5VZK9-DECOY | (?)ELYADCESLCHYLFQFLWDNPGYSYLCAEAK(?)<br>Query 4 ADCES 8<br>AD ES<br>Sbjct 171 ADIES 175<br><br>Query 7 YLCAE--AK 13<br>YL AE AK<br>Sbjct 73 YLIAENLAK 81 | immunity-specific protein Beta371 [Corynephage beta] AAA32187.1; immunity-specific protein Beta286 [Corynephage beta]AAA32186.1; hypothetical protein [Enterococcus phage EF1] ASZ76815.1; hypothetical protein [Burkholderia phage FLC9] BDD79743.1; NLR family CARD domain-containing protein-inhibition, Muti-domain, Innate Immunity, Phosphorylation [Myoviridae sp.] DAG66881.1 | 396,015 |
| 156 | O60563-2-DECOY | (?)VLPSIEK(?) | antirepressor protein [Staphylococcus phage phiSP119-2] AZB66758.1; antirepressor protein, partial [Siphoviridae sp.] DAN68257.1 | 28964,8 |
| 157 | E7ESB2-DECOY | (?)LEDLGEI(?) | nicotinamide phosphoribosyltransferase [Serratia phage 4S] QPI13761.1; tail tape measure [Bacteriophage sp.] DAS25206.1 | 11007,2 |
| 158 | P49006-DECOY | (?)GLHKIVK(?) | terminase large subunit [Shigella phage A2] UVD36852.1; terminase large subunit [Enterobacteria phage RB27] YP_009102369.1 ; capsid protein, partial [Human immunodeficiency virus 1] AGR65491.1; Chain A, AMC016 gp120 [Human immunodeficiency virus 1] 7RSO_A | 40493,7 |
| 159 | A8MVR0-DECOY | (?)GSNGNPPR(?) | YomR [Salmonella phage SPFM4] VFR10648.1; hypothetical protein [Proteus phage 7] QNN97587.1 | 205347 |
| 160 | Q5W026-DECOY | (?)KKADQGR(?) | Pectate lyase [Bacteriophage sp.] UWD61269.1; putative baseplate hub subunit and tail lysozyme [Synechococcus phage S-N03] QIN96785.1; tail tape measure protein [Lactobacillus phage JNU_P5] QHJ74756.1 | 355174 |

| | | | | |
|---|---|---|---|---|
| 161 | P61081-DECOY | (?)AASSINSR(?)<br>Query 1 AASSI 5<br>AA SI<br>Sbjct 302 AALSI 306 | Chain A, DIPHTHERIA TOXIN [Corynephage beta] 1DDT_A ; Diphtheria toxin homolog CRM228; P00589.2; hypothetical protein BA720P3_00003 [Bifidobacterium phage BA720P3] WAX05478.1; hyaluronidase [Streptococcus phage Javan548] QBX30522.1; outer capsid spike protein [Rotavirus A] AAX15951.2 | 9406 |
| 162 | H0YH26-DECOY | (?)ACCAPSHK(?) | head-to-tail adaptor [Microbacterium phage Phinky] QNJ55529.1; KID repeat-containing family protein [Clostridium phage phiCp-A] QTZ82792.1; gp120, partial [Human immunodeficiency virus 1] CAD22592.1 | 51380,2 |
| 163 | A6NNK5-DECOY | (?)QQTNTVK(?)<br>Query 2 QTNTVK 7<br>QT VK<br>Sbjct 186 QTVVVK 191 | immunity-specific protein Beta241 [Corynephage beta] AAA32184.1 ; tail tape measure [Siphoviridae sp.] DAP80073.1 ;K1 glycoprotein, partial [Human gammaherpesvirus 8]; L1 protein [Human papillomavirus] AYA94244.1; gag protein, partial [Human immunodeficiency virus 1] AYE55702.1 | 512909 |
| 164 | Q13936-9-DECOY | (?)KFLRANVTAGKR(?) | ribosylnicotinamide kinase [Caulobacter phage CcrBL9] YP_009809998.1; Rz lysis protein [Siphoviridae sp.] DAQ73662.1; Phage outer membrane lytic protein Rz; Endopeptidase [Escherichia phage Evi] VUF53346.1; neuraminidase [Influenza B virus (B/Tennessee/25/2017)] ASK80599.1 | 22031,8 |
| 165 | K7EIU7-DECOY | (?)KCQGQVR(?) | minor capsid protein [Streptococcus phage Javan554] QBX30550.1; putative ribonucleoside-diphosphate reductase 1 subunit beta [Vibrio phage 70E35.5a] QZI88703.1; gag protein, partial [Human immunodeficiency virus 1] AEE00158.1; | 1282420 |
| 166 | Q8WZ42-7-DECOY | (?)VVKYPRR(?) | repressor protein [Staphylococcus phage phiSP119-2] AZB66754.1; spike protein, partial [Infectious bronchitis virus] QWM97405.1; tegument host shutoff protein [Human alphaherpesvirus 1] | 843513 |
| 167 | Q9BX84-DECOY | (?)SEPQGGVR(?) | antirepressor protein [Myoviridae sp.] DAQ89910.1; protein of unknown function (DUF4376) [Bacteriophage sp.] DAL77723.1 | 16998,6 |
| 168 | I3L1Q4-DECOY | (?)AVSNQGVR(?) | tape measure protein [Salmonella phage SSU5] YP_006906647.1; tail length tape measure protein [Myoviridae sp.] DAP62309.1 | 48677,4 |
| 169 | F8W6H5-DECOY | (?)LDDSPER(?) | tail fiber protein [Escherichia phage KW1E_UTAR] QYA57228.1; Thioredoxin, phage-associated [Yersinia phage fPS-65] YP_010091396.1 | 17799 |
| 170 | M0R0L0-DECOY | (?)RKPGRTPDIR(?) | RNA ligase [Lactobacillus phage LpeD] YP_009835354.1; transcriptional regulator [Klebsiella phage phiKO2] YP_006625.1; DNA repair protein [uncultured Mediterranean phage uvMED] BAQ85177.1; | 1866360 |
| 171 | Q9HBX9-5-DECOY | (?)VAGDAENR(?) | CI repressor [Bacteriophage sp.] UWI00938.1; gag, partial [Human immunodeficiency virus 1] CAA76042.1 | 30180 |
| 172 | Q9UGK8-2-DECOY | (?)NGATPDEK(?) | portal protein [Haemophilus phage SuMu] YP_007002935.1; reverse transcriptase, partial [Human immunodeficiency virus 2] QTZ20165.1 | 108850 |
| 173 | Q96GN5-4-DECOY | (?)KDMAKDK(?) | Heat shock protein 60 [Podoviridae sp.] DAV62240.1; hypothetical protein [Bacteriophage sp.] UVX47616.1; chromosome segregation ATPase [Myoviridae sp.] DAT34135.1; hypothetical protein SpyM3_0935 [Streptococcus phage 315.2] NP_795472.1 | 7762,74 |
| 174 | H0Y353-DECOY | (?)SLDCESNDVKCR(?) | baseplate protein [Myoviridae sp.] DAY64649.1; DNA polymerase [Providencia phage vB_PreS_PR1] YP_009599153.1; polyprotein [rhinovirus A82] ACK37416.1 | 50487,8 |
| 175 | Q5T1R4-2-DECOY | (?)KVESSPAR(?) | putative endonuclease [Pseudomonas phage PAK_P4] YP_008859223.1; tail length tape-measure protein 1 [Pseudomonas phage AUS531phi]; | 450911 |
| 176 | Q8TBC5-DECOY | (?)MHDEPSR(?) | terminase large subunit [Bacteriophage sp.] DAH34503.1; terminase [Mycobacterium phage Bud] QBI98480.1; holin [Myoviridae sp.] DAI38574.1 | 212479 |
| 177 | P15884-DECOY | (?)QPTTPKPK(?) | hypothetical protein [Helicobacter phage DeM53M] ANT43230.1; Glycophorin A [Myoviridae sp.] DAE89502.1; nuclear antigen EBNA-3C [Human gammaherpesvirus 4] QCF48505.1 | 104253 |
| 178 | F5GXI6-DECOY | (?)AGAGSPTSQK(?)<br>Query 5 SPTSQK 10<br>SP QK<br>Sbjct 7 SPSGQK 12 | immunity-specific protein Beta241 [Corynephage beta] AAA32185.1; helix-turn-helix domain protein [Bacteriophage sp.] DAO38512.1; pol protein, partial [Human immunodeficiency virus 1] ALX35585.1; Full=Regulatory protein E2 [human papillomavirus 20] P50766.1 | 89677,4 |
| 179 | O95757-DECOY | (?)YKQDTQK(?) | Morphogenesis protein 1 wall, phi29, hydrolase, infection [Podoviridae sp.] DAL10244.1; hypothetical protein NVP1123O_42 [Vibrio phage 1.123.O._10N.286.48.F3] AUR89471.1 | 38620,1 |
| 180 | Q9H5I5-DECOY | (?)TIKHRGAK(?) | type I neck protein [Bacteriophage sp.] DAW91072.1; E1 protein [human papillomavirus 43] CAD1814277.1; E1 [human papillomavirus 91] CAD1807067.1; E1 [human papillomavirus 40] ALT54708.1; early protein [Human papillomavirus type 7] NP_041856.1; E1 [human papillomavirus 42] ACX32353.1 | 7600,47 |
| 181 | Q5T2R2-DECOY | (?)LDAPEPFK(?)<br>Query 1 LDAPEP 6<br>LD EP<br>Sbjct 271 LDLKEP 276 | immunity-specific protein Beta286 [Corynephage beta] AAA32186.1; virion structural protein [Serratia phage vB_SmaM_ 2050HW] YP_009833727.1; N-acetylmuramoyl-L-alanine amidase [Siphoviridae sp. ct1SN28] DAF84806.1; DNA-directed RNA polymerase [Siphoviridae sp.] DAZ05373.1 | 67472,5 |
| 182 | Q9H4M7-2-DECOY | (?)INANDLEK(?) | holin protein [Bacteriophage sp.] DAP58957.1; VP2 [Epizootic hemorrhagic disease virus 2] QDO66979.1; | 87462,2 |
| 183 | H7C5A9-DECOY | (?)GLAENPLSK(?)<br>Query 4 ENPLS 8<br>ENPLS<br>Sbjct 70 ENPLS 74 | Chain A, DIPHTHERIA TOXIN [Corynephage beta] 1DDT_A ; Diphtheria toxin homolog CRM228 P00589.2 ; replication protein O [Acinetobacter phage vB_AbaM_IME284] AYP68959.1; virulence associated protein E [Siphoviridae sp.] DAF76450.1 | 22382 |
| 184 | F5H7F0-DECOY | (?)KRHDASSK(?) | portal protein [Vibrio phage 1.027.O._10N.286.54.B8] AUR82708.1; hemagglutinin, partial [Influenza A virus (A/turkey/Italy/12vir6607-9/2012(H5N2))] AFU83118.1 | 29773,7 |
| 185 | EGVGQGTRK | (?)EGVGQGTRK(?) | lysozyme [Podoviridae sp.] DAN41822.1; tape measure protein [Pseudomonas phage PARCL1pr] QXM18731.1; DNA methyltransferase [Mycobacterium phage Mundrea] YP_010063246.1 | 29066,8 |
| 186 | Q6P4G0-DECOY | (?)AHTGYQQK(?) | Portal protein, Peptidoglycan hydrolase gp4, portal, tailspike, adhesin, VIRAL.5A [Podoviridae sp. ct6BA50] DAG05699.1; HNH endonuclease [Streptococcus phage phi3396] YP_001039949.1 | 14267,6 |
| 187 | O95772-DECOY | (?)IIVCVCVK(?) | DNA polymerase subunit [Aeromonas phage Aes123] AFN69891.1 ; 19.3 kDa MHC class I antigen-binding glycoprotein precursor [Human mastadenovirus B] AKQ98425.1; neuraminidase, partial [Influenza A virus (A/Denmark/33/2008(H1N1))] ACI25642.1 | 8895,49 |
| 188 | P45379-7-DECOY | (?)LLLRKHR(?) | immunity-specific protein Beta241 [Corynephage beta] AAA32184.1; putative ATPase [Corynebacterium phage phi16] APQ42604.1; Putative regulator of cell autolysis [Myoviridae sp.] DAT61446.1 | 3308,75 |
| 189 | G3V5X4-DECOY | (?)AGMAVSEFK(?) | putative exonuclease [Serratia phage vB_SmaS_Rovert] QPX75000.1; virion structural protein [Bacillus phage Spock] YP_008770325.1 | 6626,28 |
| 190 | Q99956-DECOY | (?)GVTVVEQSK(?)<br>Query 4 VVEQSK 9<br>VV+ SK<br>Sbjct 5 VVDSSK 10 | Chain A, DIPHTHERIA TOXIN [Corynephage beta] 1DDT_A; immunity-specific protein Beta286 [Corynephage beta] AAA32186.1; DNA gyrase/topoisomerase IV, subunit A [Bacillus phage vB_BanS_MrDarsey] UGO47949.1 | 12190,6 |
| 191 | B4DVD1-DECOY | GVLSQDISK | hypothetical protein [Enterococcus phage VPE25] SCO93501.1; | 12596,7 |
| 192 | F2Z2S2-DECOY | (?)IGVLLEMR(?)<br>Query 1 IGVL 4<br>IGVL<br>Sbjct 509 IGVL 512<br>Query 2 GVLL 5<br>GVLL<br>Sbjct 431 GVLL 434 | Chain A, DIPHTHERIA TOXIN [Corynephage beta] 1DDT_A; tail protein [Acinetobacter phage vB_AbaS_TRS1] YP_009289768.1; DNA polymerase [Pectobacterium phage DU_PP_V] YP_009795303.1 | 72894,7 |
| 193 | Q9UJF2-2-DECOY | (?)QIAQAEPAK(?) | tail tape measure protein [Bacteriophage sp.] DAW73697.1; putative terminase large subunit [uncultured Caudovirales phage] ASN69115.1 | 419378 |
| 194 | Q9UBK2-9-DECOY | (?)LLKIPNHK(?) | histone deacetylase [Satyrvirus sp.] AYV85460.1; RelB antitoxin [Bacteriophage sp.] UWG11619.1; glycoprotein [Lymphocytic choriomeningitis mammarenavirus] ACV72558.1 | 4533,96 |
| 195 | B4DWY7-DECOY | (?)AQGGSGPLGPK(?)<br>Query 2 QGGSG 6<br>QG SG<br>Sbjct 411 QGESG 415 | Chain A, DIPHTHERIA TOXIN [Corynephage beta] 1DDT_A; putative lectin-like domain protein [Brevundimonas phage vB_BpoS-Gurke] UTC28405.1; tail protein [Siphoviridae sp.] DAR16869.1 | 5261,2 |
| 196 | F6X5S1-DECOY | (?)LSMGESFAK(?)<br>Query 5 ESFAK 9<br>E FAK<br>Sbjct 7 EQFAK 11 | Diphtheria toxin homolog CRM228 P00589.2 Dna polymerase B [Bacteriophage sp.] DAH90912.1; iron-sulfur cluster biosynthesis protein [Marseillevirus LCMAC101] QBK86171.1 | 8303,8 |
| 197 | B4DZ46-DECOY | (?)TAQTGKQIK(?)<br>Query 1 TAQTGK 6<br>TAQ GK<br>Sbjct 153 TAQRGK 158 | immunity-specific protein Beta241 [Corynephage beta] AAA32184.1; major capsid protein [Streptococcus phage SOCP] AID18053.1; site-specific recombinase [Lactobacillus phage JNU_P10] QHJ75141.1 | 17296,8 |
| 198 | J3QLA7-DECOY | (?)RLVGMEKK(?)<br>Query 3 VGMEK 7<br>VG EK<br>Sbjct 146 VGTEK 150 | diphtheria toxin, partial [Corynephage beta] ABU25232.1; tail assembly chaperone protein [Bacteriophage sp.] UVN10603.1; vpu protein [Human immunodeficiency virus 1] QJQ91397.1 | 47981,8 |
| 199 | E7EX90-DECOY | (?)AFIAITTDK(?) | holin [Siphoviridae sp.] DAF25580.1; E2 protein [Human papillomavirus] AYA94267.1 | 87811,1 |
| 200 | Q14721-DECOY | (?)ILLRREGK(?) | hypothetical protein BPABA456_00410 [Acinetobacter phage YMC/09/02/B1251] YP_007010622.1; gag protein, partial [Human immunodeficiency virus 1] AYE55620.1 | 7973,08 |
| 201 | Q9Y493-4-DECOY | (?)LPLRLYSK(?) | tail family protein [Streptococcus phage phiNIH1.1] NP_438155.1; putative tail tubular protein B [Pseudomonas phage LKA1] YP_001522886.1 | 54345,9 |
| 202 | Q9Y2L9-DECOY | (?)TVMAFPLPK(?) | phage excisionase [Escherichia coli O145:H28] [Stx2a-converting phage Stx2_14040] BCI48934.1; hypothetical protein [Bacteriophage sp.] UVY34807.1; polyprotein [Dengue virus type 4] UCQ65263.1 | 106471 |
| 203 | Q92824-DECOY | (?)EPGTKPLIR(?)<br>Query 3 GTKP 6<br>GTKP<br>Sbjct 22 GTKP 25 | Chain A, DIPHTHERIA TOXIN [Corynephage beta] 1DDT_A; metallophosphatase domain protein [Bacteriophage sp.] DAO98194.1; chitin synthase regulator [Bacteriophage sp.] DAH08503.1; envelope glycoprotein E [Human alphaherpesvirus 3] AQT34120.1 | 37247,8 |
| 204 | Q04760-DECOY | (?)AVDGKEENR(?) | Rep protein [Bacteriophage sp.] DAS63005.1; Portal [Bacteriophage sp.] DAY39946.1; envelope glycoprotein [Human immunodeficiency virus 1] AHC00335.1 | 6773,19 |
| 205 | M0R0S4-DECOY | (?)IIVLLEVPK(?) | restriction enzyme [Bacteriophage sp.] DAW87427.1; envelope glycoprotein, partial [Human immunodeficiency virus 1] AWF48941.1 | 6347,85 |
| 206 | P0CG42-DECOY | (?)FKNCDESK(?) | terminase large subunit [Bacteriophage sp.] DAT14571.1; | 10534,8 |
| 207 | P19113-DECOY | (?)GVSLQDAPIK(?)<br>Query 2 VSLQDAPI 9<br>VS DAPI<br>Sbjct 41 VSI-DAPI 47 | immunity-specific protein Beta286 [Corynephage beta] AAA32186.1; phage tape measure protein [uncultured Mediterranean phage uvMED] BAR35206.1; Cell wall hydrolase autolysin [Siphoviridae sp.] DAL93499.1 | 68174,7 |
| 208 | Q5T8R8-DECOY | (?)LRALSIHLK(?)<br>Query 3 ALSI 6<br>ALSI<br>Sbjct 303 ALSI 306 | Chain A, DIPHTHERIA TOXIN [Corynephage beta] 1DDT_A; integrase [Alteromonas phage P24] AZU97334.1; tail length tape-measure protein [Streptococcus phage Javan199] QBX15678.1; polymerase [Hepatitis B virus] ACO05395.1 | 18557,8 |
| 209 | Q09MP3-DECOY | (?)LVLTAVKVPK(?) | replicase [Lactobacillus phage ATCC8014] YP_007236720.1; DNA-binding phosphoprotein (1), partial [Monkeypox virus] UZG89539.1; putative DNA-binding phosphoprotein [Vaccinia virus] YP_232938.1 | 6271,76 |
| 210 | Q9H3S1-2-DECOY | (?)HDGSVSRKLLR(?)<br>Query 5 VSRKL 9<br>VSRKL<br>Sbjct 4 VSRKL 8<br>Query 3 GSVSRKLLR 11<br>GSVSR LLR<br>Sbjct 520 GSVSR-LLR 527 | diphtheria toxin, partial [Corynephage beta] ABU25232.1; hypothetical protein FRC0104_02371 [Corynebacterium diphtheriae] CAB0765614.1; putative transcriptional regulator, partial [Siphoviridae sp.] DAU44897.1; anaerobic ribonucleoside reductase large subunit [Lactococcus phage P1048] 009905618.1; Dna polymerase B [Siphoviridae sp.] DAZ26473.1 | **3,70E+12** |

| | | | | |
|---|---|---|---|---|
| 211 | Q16854-4-DECOY | (?)DMLGYPHLK(?) | glycosyltransferase [Brevibacterium phage LuckyBarnes] 009792215.1 ; terminase large subunit [Enterococcus phage EfV12-phi1] YP_009814808.1 ; hemagglutinin precursor [Influenza A virus (A/equine/Santiago/1/1985(H3N8))] AAA43110.1 | 7919,13 |
| 212 | P07288-2-DECOY | (?)LSYASNCFK(?) | 6-pyruvoyl tetrahydropterin synthase [Bacteriophage sp.] DAP29135.1; replication initiator protein [Microviridae sp.] WAE43772.1; leucine-rich repeat protein [Siphoviridae sp.] DAU15445.1 | 4396,55 |
| 213 | Q5T987-DECOY | (?)IRHVQGKMK(?) | terminase large subunit [Salinibacter phage M8CRM-1] YP_009639469.1; head morphogenesis [Lactobacillus phage LBR48] YP_009168589.1 | 342943 |
| 214 | Q9P2N4-4-DECOY | (?)VLANSKEKLK(?) | tail length tape-measure protein 1 [Klebsiella phage vB_KpnS_ZX4] YP_010054541.1; DNA binding protein [Bacillus phage YungSlug] QKE56345.1 | 101983 |
| 215 | Q8N9W6-4-DECOY | (?)LMLSMGVQDK(?) | hypothetical protein Javan23_0034 [Streptococcus phage Javan23] QBX15999.1; holin [Mycobacterium phage ZoeJ] YP_009032424.1; tail length tape-measure protein 1 [Klebsiella phage vB_KpnS_ZX4] YP_010054541.1 | 8525,44 |
| 216 | Q6ZSC3-DECOY | (?)MDLTDTSLNK(?) | GTP cyclohydrolase I [Bacteriophage sp.] UVX53378.1; putative pentapeptide repeat protein [Staphylococcus phage vB_SauM_VL10] WAX23382.1; hemagglutinin [Influenza A virus (A/Texas/83/2014(H3N2))] APB91074.1 | 1133600 |
| 217 | Q14833-5-DECOY | (?)FDAILGAEPDK(?) | hypothetical protein FDJ60_gp52 [Pseudomonas phage JBD67] YP_009625975.1; tape measure protein [Gordonia phage Margaret] AWY05199.1; immunity repressor [Gordonia phage Fairfaxidum] YP_009822332.1 | 268862 |
| 218 | Q13936-32-DECOY | (?)SDLMLKQNSK(?) | portal protein [Bacteriophage sp.] UWG25738.1; repressor domain protein [Siphoviridae sp.] DAV34810.1; hemagglutinin [Influenza A virus] QWO70866.1 | 76787,2 |
| 219 | J3QST1-DECOY | (?)SWEDAKSASGR(?)<br>Query 2 WEDAKSASG 10<br>W D K ASG<br>Sbjct 139 WADRKEASG 147<br><br>Query 2 WEDAK 6<br>WE AK<br>Sbjct 153 WEQAK 157 | immunity-specific protein Beta201 [Corynephage beta] AAA32185.1 ; Chain A, DIPHTHERIA TOXIN [Corynephage beta] 1DDT_A; Diphtheria toxin homolog CRM228 P00589.2; Muramidase (flagellum-specific) [Siphoviridae sp.] DAT93538.1; tail tube protein [Bacteriophage sp.] DAV39086.1; putative portal protein [Pseudomonas phage VB_PaeS_VL1] UGV19880.1; aggregation promoting factor [Enterococcus phage vB_EfaM_Ef2.3] QBZ70101.1; transglycosylase [Enterococcus phage EF24C] YP_001504119.1 | 15636,9 |
| 220 | P22455-2-DECOY | (?)NAVLRQLEPR(?) | putative tail spike protein [Klebsiella phage Geezett] QXV72116.1; portal vertex of the head [Klebsiella phage Kp_GWPR59] WAW44582.1; portal protein [Bacteriophage sp.] DAL66737.1 | 260571 |
| 221 | M0R0L0-DECOY | (?)NNDIDFDICK(?) | ATPase [Bacteriophage sp.] DAD57711.1; DNA polymerase [Siphoviridae sp.] DAS43076.1; baseplate wedge initiator [Synechococcus phage ACG-2014f]; AIX31745.1; Cytosine specific methyltransferase [Siphoviridae sp. ctxYv12] DAF45729.1 | 1866360 |
| 222 | O15354-DECOY | (?)MLFFPTIPDTR(?)<br>Query 5 PTIP 8<br>PTIP<br>Sbjct 435 PTIP 438 | Chain A, DIPHTHERIA TOXIN [Corynephage beta] 1DDT_A; Diphtheria toxin homolog CRM228 P00589.2; 6b protein [Infectious bronchitis virus] ADV71775.1; hypothetical protein HOU32_gp177 [Dickeya phage vB_DsoM_JA11] YP_009813622.1 | 4767800 |
| 223 | G8JLB2-DECOY | (?)VMGSQCMPLLSR(?) | hypothetical protein [Bacteriophage sp.] DAF02146.1; hypothetical protein [Propionibacterium phage pa6919-4] AUV62324.1; envelope glycoprotein, partial [Human immunodeficiency virus 1] AAO24041.1; large S protein [Hepatitis B virus] ADF06177.1; | 3233740 |
| 224 | K4DIA0-DECOY | (?)RDASFPETAMGR(?) | Baseplate J like protein [Bacteriophage sp.] DAR09938.1; Baseplate J like protein [Siphoviridae sp.] DAG66204.1; ATP dependent DNA helicase [Siphoviridae sp.] DAT77930.1; N acetylmuramoyl L alanine amidase endolysin [Myoviridae sp.] DAS42540.1; pol protein, partial [Human immunodeficiency virus 1] UVF23472.1 | 1483240 |
| 225 | Q6PJT7-DECOY | (?)LLLGGVPLRRGSAR(?)<br>Query 1 LLLGGV 6<br>LL GG+<br>Sbjct 23 LLKGGI 28 | immunity-specific protein Beta201 [Corynephage beta] AAA32185.1 ; head decoration [Corynebacterium phage Darwin] YP_009620270.1; Replication initiation factor [Inoviridae sp.] DAR73470.1; endonuclease [Gordonia phage OneUp] YP_009274525.1; pectate lyase superfamily protein [Klebsiella phage GADU21] UOX39351.1 | 1031130 |
| 226 | O43768-3-DECOY | (?)RVPVVKESVWKPR(?) | HNH nuclease [uncultured Caudovirales phage] CAB4166728.1; RNA dependent RNA polymerase [Siphoviridae sp.] DAQ07376.1; terminase [Mycobacterium phage Ekdilam] YP_009952058.1; DNA primase [Salmonella phage Chi] YP_008058159.1 | 1791730 |
| 227 | H3BMZ9-DECOY | (?)VAGSGIWRVLRLPR(?)<br>Query 9 VLRLP 13<br>VL LP<br>Sbjct 135 VLSLP 139 | Chain A, DIPHTHERIA TOXIN [Corynephage beta] 1DDT_A; immunity-specific protein Beta371 [Corynephage beta] AAA32187.1; Diphtheria toxin homolog CRM228 P00589.2;<br>portal protein [Achromobacter phage JWX] YP_009196194.1; tail protein [Caudovirales sp.] DAR60063.1; DNA mismatch endonuclease [Siphoviridae sp.] DAJ68944.1 | 5564380 |
| 228 | Q0VDD8-DECOY | (?)AHVFSIHHKVMFR(?) | hypothetical protein [Bacteriophage sp.] DAW92603.1; hypothetical protein [Klebsiella phage vB_KpnM_VAC13] UJB55111.1; putative portal protein [Acinetobacter phage BS46] QEP53201.1 | 2064110 |
| 229 | F5H6V4-DECOY | (?)NIVLVKAISTAVSLKR(?) | holin [Siphoviridae sp.] DAL45079.1; tail tape measure [Siphoviridae sp.] DAO32156.1; Chain A, Nucleoprotein [Human parainfluenza 3 virus (strain NIH 47885)] 7EV8_A | 3679870 |
| 230 | Q2PPJ7-3-DECOY | FASVPFALK<br>Query 2 ASVPF-ALK 9<br>AS PF A+K Sbjct<br>57 ASTPFEAMK 65 | immunity-specific protein Beta241 [Corynephage beta] AAA32184.1 | **1,08E+12** |
| 231 | Q96S53 | (?)QDLMGGK(?)<br>Query 2 DLMGG 6<br>DLMGG<br>Sbjct 132 DLMGG 136<br>**Homo sapiens**<br>Query 1 QDLMGGK 7<br>QDLMGGK<br>Sbjct 406 QDLMGGK 412 | immunity-specific protein Beta201 [Corynephage beta] AAA32185.1; glycerol-3-phosphate dehydrogenase/oxidase [Corynebacterium simulans] WP_062036793.1; RNA-dependent RNA polymerase [Bacteriophage sp.] DAN49619.1 ; membrane-bound protein [Bacillus phage vB_BthM-Goe5] AZF89335.1; holin [Siphoviridae sp.] DAE48563.1; Leucine-rich repeat [Bacteriophage sp.] UWD75692.1; rev protein [Human immunodeficiency virus 2] BAM76160.1; dual specificity testis-specific protein kinase 2 isoform 1 [Homo sapiens] NP_009101.2 | **1,51E+12** |
| 232 | P61018-2-DECOY | (?)GEVPNAIDILR(?) | immunity-specific protein Beta286 [Corynephage beta] AAA32186.1; HNH endonuclease [Pseudomonas phage vB_PsyM_KIL1] YP_009276041.1; Terminase large subunit [Siphoviridae sp.] DAZ28584.1; nonstructural protein 1 [Influenza A virus (A/chicken/Vietnam/29/2007(H5N1))] ACB70641.1 | 60860,9 |
| 234 | A2RUS2-DECOY | (?)MVSFSQSQKR(?) | Phage major capsid protein, HK97 [uncultured Caudovirales phage] CAB4142186.1; terminase [Bacteriophage sp.] UVM81712.1; gene 34 protein [Shigella phage Sf6] NP_958210.1 | 485446 |
| 235 | P00740-2-DECOY | (?)KAQMNYNYR(?) | tail tape measure protein [Siphoviridae sp.] DAO66189.1; hypothetical protein [Bacteriophage sp.] UWI30234.1; Lysozyme-like Peptidoglycan-binding protein [Myoviridae sp.] DAS80482.1; polymerase PB1 [Influenza A virus] UUB81356.1; antitoxin [Siphoviridae sp.] DAZ76960.1 | 293712 |
| 236 | Q6PIV2-DECOY | (?)DPPKKVASIPR(?) | tail tape measure [Siphoviridae sp.] DAI07704.1; stabilization protein [Bacteriophage sp.] DAV95370.1; stabilization protein [Podoviridae sp.] DAK43789.1; L2 [Human papillomavirus 202] AKP16352.1 | 5885,14 |
| 237 | P13942-5-DECOY | (?)NEPLPLSSDSSPILR(?)<br>Query 1 NEPLPLSSDS 10<br>NE SSDS<br>Sbjct 502 NE---ISSDS 508<br><br>Query 4 LPLSSDSS 11<br>LP SS<br>Sbjct 138 LPFAEGSS 145 | Chain A, DIPHTHERIA TOXIN [Corynephage beta] 1DDT_A; RNA-directed RNA polymerase [ssRNA phage SRR7976299_2] DAD52667.1; head protein [Edwardsiella phage MSW-3] YP_007348919.1 | 279570 |
| 238 | A2AAS7-DECOY | (?)LEGNMFFPFSFCPSLTVEPIMVVQRESPMLGAST CPTWKILATLRFGLVQPPLMSR(?) | immunity-specific protein Beta371 [Corynephage beta] AAA32187.1; hypothetical protein [Bacteriophage sp.] UVM92066.1; DNA primase [Siphoviridae sp.] DAS85355.1; thioredoxin [Escherichia phage UPEC01] DAE03409.1; DNA primase, catalytic core [Siphoviridae sp. ctTXt1] DAE03409.1; putative transcriptional regulator [Siphoviridae sp.] DAW46092.1; holin family protein [Bacteriophage sp.] UVX92009.1; late protein L2 [human papillomavirus 52] AEI61572.1 | **1,08E12** |
| 239 | Q6ZUM4-4-DECOY | (?)AISSKIEKKSPK(?) | immunity-specific protein Beta371 [Corynephage beta] AAA32187.1; ABC transporter permease [Corynebacterium diphtheriae] CAB0632219.1; dsdna helicase [Bacteriophage sp.] UOF79517.1; toxin [Siphoviridae sp.] DAI78401.1; DNA-binding protein [Siphoviridae sp.] DAT22249.1 | **3,19E12** |
| 240 | Q93084-2-DECOY | (?)LGKQNNMQGVSRVWVRVSK(?) | immunity-specific protein Beta286 [Corynephage beta] AAA32186.1; immunity-specific protein Beta241 [Corynephage beta] AAA32184.1; hypothetical protein AOT42_10540 [Corynebacterium diphtheriae bv. gravis]OKY22735.1; G/U mismatch-specific DNA glycosylase [Bacteriophage sp.] DAY57160.1; portal protein [Siphoviridae sp.] DAU75319.1; ORF39 [Human alphaherpesvirus 3] AON76579.1 | **1,38E12** |
| 241 | A6NJ46,A6NJ46-2,P45379-7-DECOY | (?)LLLRKHR(?) | immunity-specific protein Beta286 [Corynephage beta] AAA32186.1; putative ATPase [Corynebacterium phage phi16] APQ42604.1; Putative regulator of cell autolysis [Myoviridae sp.] DAT61446.1; replication initiator protein [Microviridae sp.] AXL14850.1; WhiB family transcriptional regulator [Corynebacterium sp. HMSC08A12] WP_070822393.1 | 3308,75 |
| 242 | P52306-4-DECOY | (?)SKCEFIFSEK(?) | immunity-specific protein Beta201 [Corynephage beta] AAA32185.1 ; complement control protein homologue [Saimiriine gammaherpesvirus 2] NP_040206.1; RNA dependent RNA polymerase [Picornaviridae sp.] URG14375.1; putative resolvase [Ralstonia phage Rs551] YP_009786106.1 | 7696,34 |
| 243 | Q9BQI5-4-DECOY | (?)NCDPTIGGLDGR(?) | immunity-specific protein Beta286 [Corynephage beta] AAA32186.1; DNA polymerase [Pelagibacter phage HTVC200P] AXH71585.1; | 6139,32 |
| 244 | Q9Y2G1-DECOY | (?)IENLGGEAQMR(?) | immunity-specific protein Beta241 [Corynephage beta] AAA32184.1; putative structural lysozyme [Klebsiella phage N1M2] QGH72084.1 ; head-closure protein [Vibrio phage 1.182.O._10N.286.46.E1] AUR93054.1 | 7784,83 |
| 245 | J3KPZ3-DECOY | (?)LGANCPFVHLTIRSEGEGSIVCIASPHLLEKGRIPG LVPLCNPVDAVAMK(?)<br>Query 13 RSEGEGSIVCI 23<br>RS G GS CI<br>Sbjct 218 RSVG-GSLSCI 227<br><br>Query 16 GEGSIVCIA 24<br>G GS+ IA<br>Sbjct 334 GIGSVMGIA 342<br><br>Query 16 GEGSIVCIA 24<br>G GS+ IA<br>Sbjct 309 GIGSVMGIA 317 | Diphtheria toxin homolog CRM228 P00589.2; Chain A, DIPHTHERIA TOXIN [Corynephage beta] 1DDT_A; glycosyltransferase [Streptomyces phage Gilson] YP_009842485.1; E4 protein [Human papillomavirus] AYA93888.1; tail length tape measure protein [Staphylococcus phage B166] YP_009204078.1 | 17180,8 |
| 246 | E5RGL8-DECOY | (?)FPRPGNPYCGK(?)<br>Query 6 NPYCGK 11<br>NP GK<br>Sbjct 71 NPLSGK 76 | Chain A, DIPHTHERIA TOXIN [Corynephage beta] 1DDT_A; Diphtheria toxin homolog CRM228 P00589.2; Major tail protein [Bacteriophage sp.] DAX25031.1; glycoprotein [Human respiratory syncytial virus B] BBD49505.1 | 13271,1 |
| 247 | K7EJU0-DECOY | (?)CGQNPSKAAFR(?) | DNA polymerase I [Streptomyces phage TP1604] YP_009200163.1; YP_009200163.1 | 21303,9 |
| 248 | H3BTZ6-DECOY | (?)IGGTTQGSEQIR(?) | immunity-specific protein Beta371 [Corynephage beta] AAA32187.1; tape measure protein [Bacillus phage vB_BanS_Nate] UGO51069.1; nucleoprotein [Tacaribe mammarenavirus] QLB38570.1 | 14574,8 |
| 249 | F5H0U9-DECOY | (?)TQIDAKWETR(?) | immunity-specific protein Beta371 [Corynephage beta] AAA32187.1; Dihydrofolate reductase [Phage NCTB] SBV38338.1; putative YadA-like protein [Aeromonas phage D3] QDJ97161.1 | 40995,9 |
| 250 | Q30KQ5-DECOY | (?)MVEGATLSRQR(?) | glutamine amidotransferases class-II [Bacteriophage sp.] UVY01503.1; major capsid protein [Human betaherpesvirus 5] ARX80298.1 | 67379,7 |

| | | | | |
|---|---|---|---|---|
| 251 | H7C4J8-DECOY | (?)IAVGDMVVAYSK(?)<br>Query 1 IAVGDMVVA 9<br>IA + VA<br>Sbjct 333 IALSSLMVA 341<br><br>Query 7 VVAYSK 12<br>VV SK<br>Sbjct 5 VVDSSK 10 | Chain A, DIPHTHERIA TOXIN [Corynephage beta] 1DDT_A; Diphtheria toxin homolog CRM228 P00589.2; Terminase [Bacteriophage sp.] DAY34796.1; ORF1ab polyprotein, partial [Severe acute respiratory syndrome coronavirus 2] QKF93248.1 | 63787,7 |
| 252 | Q16653-DECOY | (?)QKPITLRSVAR(?) | portal protein [Mycobacterium phage Ellie] YP_009952151.1; adsorption-associate tail protein [Staphylococcus phage Stab23] VEV88538.1 | 18082,9 |
| 253 | H3BQ24-DECOY | (?)DDRHNRLGCK(?) | hypothetical protein [Corynephage phi16] CAA73076.1; carbamoyltransferase [Synechococcus phage ACG-2014d] AIX37230.1; LysA like endolysin [Siphoviridae sp.] DAE95790.1 | 13293,8 |
| 254 | Q8N9W6-3-DECOY | (?)YPGRSMERMK(?) | minor tail protein [Bacteriophage sp.] DAO41465.1; envelope glycoprotein [Human immunodeficiency virus 1] ACE76326.1 | 8165,93 |
| 255 | Q92837-DECOY | (?)SDDGDDSNSSYK(?)<br><br>Query 2 DDGDDSN--------SSY 11<br>DD DS SSY<br>Sbjct 3 DDVVDSSKSFVMENFSSY 20 | Chain A, DIPHTHERIA TOXIN [Corynephage beta] 1DDT_A; Diphtheria toxin homolog CRM228 P00589.2; envelope glycoprotein [Human immunodeficiency virus 1] AYX71285.1; envelope glycoprotein [Human immunodeficiency virus 1] AKN90061.1; gp160 protein [Human immunodeficiency virus 1]; envelope glycoprotein [Human immunodeficiency virus 1] AYX71214.1 | 4952,7 |
| 256 | O75445-DECOY | (?)LWAHLSSDLVNLESQSIRRTAQSMPWLGALITYTNSTK(?)<br>Query 16 SIRRT 20<br>SI RT<br>Sbjct 404 SIIRT 408<br>Query 10 VNLESQSIR 18<br>+NL+ IR<br>Sbjct 202 INLDWDVIR 210<br>Query 5 LSSDLVNLE 13<br>LS V LE<br>Sbjct 159 LS---VELE 164<br>Query 2 WAHLSSDLVN----LESQSIRRTA 21<br>WA VN +S+ TA<br>Sbjct 281 WA------VNVAQVIDSE----TA 294 | Chain A, DIPHTHERIA TOXIN [Corynephage beta] 1DDT_A; immunity-specific protein Beta286 [Corynephage beta] AAA32186.1; Diphtheria toxin homolog CRM228 P00589.2; hypothetical protein EMB2_00020 [Bacteroides phage EMB2] USR81539.1 | 19887,7 |
| 257 | Q9P225-2-DECOY | (?)EGGNFLPGDFALQVSPANQIETVGLMLQEGSAGCESAERVFMPSGQYKLWELNSQAQSKQMSR(?)<br>Query 11 ALQVSPANQIETV----GLMLQE 29<br>AL V A ET+ GL L E<br>Sbjct 93 ALKVDNA---ETIKKELGLSLTE 112 | Chain A, DIPHTHERIA TOXIN [Corynephage beta] 1DDT_A; Diphtheria toxin homolog CRM228 P00589.2; hypothetical protein Atoyac15_33 [Aeromonas phage Atoyac15] QOC54397.1 | 42,631 |
| 258 | Q92837-DECOY | (?)LLGSSQTASRKR(?) | C-5 cytosine-specific DNA methylase [Bacteriophage sp.] UWI15938.1; E1 [Gammapapillomavirus sp.] YP_009552550.1 | 9596,63 |
| 259 | G8JLB2-DECOY | (?)VMGSQCMPLLSR(?) | hypothetical protein D861_gp39 [Escherichia phage vB_EcoS_ACG-M12] YP_006987858.1; envelope glycoprotein, partial [Human immunodeficiency virus 1] ABA08263.1 | 3233740 |
| 260 | O15354-DECOY | (?)MLFFPTIPDTR(?)<br>Query 5 PTIP 8<br>PTIP<br>Sbjct 435 PTIP 438 | Chain A, DIPHTHERIA TOXIN [Corynephage beta] 1DDT_A; Diphtheria toxin homolog CRM228 P00589.2; putative helix-destabilizing protein [Prokaryotic dsDNA virus sp.] QDP57287.1 ; putative 7 protein [Infectious bronchitis virus] UOF83414.1 | 4767800 |
| 261 | Q03001-8-DECOY | (?)EPVQDKETRKGPPK(?) | dsDNA helicase [Bacteriophage sp.] DAF23414.1; baseplate protein [Bacteriophage sp.] UVN08078.1; reverse transcriptase, partial [Human immunodeficiency virus 1] ACB38325.1 | 412650 |
| 262 | E5RFT4-DECOY | (?)FESLSSTLIVKK(?) | protein of unknown function DUF859 [Siphoviridae sp.] DAZ26141.1; minor tail protein [Bacteriophage sp.] UWF91564.1 | 133225 |
| 263 | K4DIA0-DECOY | (?)RDASFPETAMGR(?) | Baseplate J like protein [Bacteriophage sp.] DAR40887.1; N acetylmuramoyl L alanine amidase endolysin [Myoviridae sp.] DAS42540.1; pol protein, partial [Human immunodeficiency virus 1] UVF23492.1 | 1483240 |
| 264 | Q9NXG6-3-DECOY | (?)APTKMRQLSDGSDRPESCTGELMMARPEQMDGHDTTAQDSHALTTCGVIGSR(?) | immunity-specific protein Beta241 [Corynephage beta] AAA32184.1; hypothetical protein [Bacteriophage sp.] DAG33849.1; DnaA protein [Bacteriophage sp.] UWG88161.1 | 69,002 |
| 265 | Q8WZ42-DECOY | (?)KPIAKPEGALGGPK(?) | peptidase [Burkholderia phage BcepSauron] YP_009904610.1; portal protein [Bacteriophage sp.] UVY05965.1 | 230818 |
| 266 | E9PQB1-DECOY | (?)LLPHTTGALVTNK(?)<br>Query 5 TTGAL 9<br>TT AL<br>Sbjct 300 TTAAL 304<br><br>Query 1 LLPHTTGALVTNK 13<br>LLP G L NK<br>Sbjct 433 LLPTIPGKLDVNK 445 | Chain A, DIPHTHERIA TOXIN [Corynephage beta] 1DDT_A; Diphtheria toxin homolog CRM228 P00589.2; major head protein [Synechococcus phage S-MbCM100] YP_009008057.1; envelope glycoprotein [Human immunodeficiency virus 1] ACO94092.1 | 94260,1 |
| 267 | E9PI10-DECOY | (?)ANNQLRKLIWK(?) | Terminase small subunit [Bacteriophage sp.] UWI17863.1; | 69502,3 |
| 268 | E9PPV8-DECOY | (?)LLPDKLGETELR(?) | tRNA amidotransferase [Vibrio phage nt-1] YP_008125198.1; replisome organizer [Bacteriophage sp.] DAO93909.1; envelope glycoprotein [Human immunodeficiency virus 1] AGC81519.1 | 3154800 |
| 269 | Q01433-3-DECOY | (?)AMKLKGMKGNFK(?) | DNA adenine methylase [Myoviridae sp.] DAU43735.1; hypothetical protein [Bacteriophage sp.] DAP43264.1; surface glycoprotein [Severe acute respiratory syndrome coronavirus 2] QNO98003.1 | **2,58E12** |
| 270 | Q13402-5-DECOY | (?)MSTNAQSMNGIVK (?) | tail tape measure protein [Siphoviridae sp.] DAT40075.1; integrase/recombinase [Streptococcus satellite phage Javan624] QBX11993.1; portal protein [Streptococcus phage Javan399] QBX18159.1 | 12035,1 |
| 271 | Q8TER0-5-DECOY | (?)ANSPQALPINRPK(?) | gp164 [uncultured Mediterranean phage uvMED] BAR32967.1 ; VP1, partial [Norovirus GI] QHD93093.1; polymerase/replicase [Infectious bronchitis virus] AFQ31008.1; RNA-dependent RNA polymerase, partial [Infectious pancreatic necrosis virus] AEX31266.1 | 8024,65 |
| 272 | Q6PJT7-DECOY | (?)LLLGGVPLRRGSAR(?) | head decoration [Corynebacterium phage C3PO] YP_009620174.1; beta-lactamase CTX-M-15 [Enterobacter phage LAU1] QHR63338.1; Replication initiation factor [Inoviridae sp.] DAR73470.1; pectate lyase superfamily protein [Klebsiella phage GADU21] UOX39351.1 | 1031130 |
| 273 | P51825-3-DECOY | (?)IQRAPGPGSAAHGVR(?) | immunity-specific protein Beta241 [Corynephage beta] AAA32184.1; Queuosine biosynthesis protein QueC [Bacteriophage sp.] QGZ17813.1; nef protein [Human immunodeficiency virus 1] AAN03108.1; tegument protein UL37 [Human alphaherpesvirus 2] AMB66001.1 | 82513,2 |
| 274 | Q5SQH8-DECOY | (?)VAIEPTVLAPTVLR(?)<br>Query 7 VLAPTV 12<br>VL PT+<br>Sbjct 432 VLLPTI 437 | Chain A, DIPHTHERIA TOXIN [Corynephage beta] 1DDT_A; Diphtheria toxin homolog CRM228 P00589.2; hypothetical protein JK55_00045 [Shigella phage JK55] QEG06707.1; virion structural protein [Alphaproteobacteria phage PhiJL001] YP_224012.1; envelope glycoprotein [Human immunodeficiency virus 1] ASU63178.1 | 1067,39 |
| 275 | Q9NQ66-2-DECOY | (?)YFDSMAPAACEPREDNLPKSPANDECVPVMNEPCQSEENRLVR(?) | inner spore coat protein [Bacteriophage sp.] DAW11518.1; hypothetical protein [Klebsiella phage 05F01] BBK09298.1; tail length tape measure protein [Pseudomonas phage crassa] YP_009914476.1 | 15,426 |
| 276 | E7ERB7-DECOY | (?)LPEIGITKRVLPR(?)<br>Query 1 LPEIG 5<br>LP IG<br>Sbjct 307 LPGIG 311 | Chain A, DIPHTHERIA TOXIN [Corynephage beta]1DDT_A; immunity-specific protein Beta371 [Corynephage beta] AAA32187.1; Diphtheria toxin homolog CRM228 P00589.2; endolysin [Proteus phage PM 75] YP_009150313.1; tail sheath [Pseudomonas phage PhiPA3] YP_009217243.1; polymerase P3 [Influenza D virus] QTP31438.1 | 71628,2 |
| 277 | Q5SZK8-DECOY | (?)DIPAPPPCTVEASFK(?) | immunity-specific protein Beta371 [Corynephage beta] AAA32187.1; Gp2.5-like ssDNA binding protein and ssDNA annealing protein [Vibrio phage vB_VchM-138] YP_007006392.1; hypothetical protein UFOVP36_5 [uncultured Caudovirales phage] CAB4122271.1 polyprotein, partial [Hepatitis C virus subtype 3a] AOF41208.1 | 22583 |
| 278 | Q9NUM4-DECOY | (?)THSVGNSSLGWSLAR(?) | immunity-specific protein Beta371 [Corynephage beta] AAA32187.1; hypothetical protein UFOVP249_45 [uncultured Caudovirales phage] CAB4132826.1 ; surface glycoprotein [Severe acute respiratory syndrome coronavirus 2] UHI87525.1; surface glycoprotein [Severe acute respiratory syndrome coronavirus 2] UCP54115.1; large surface protein [Hepatitis B virus] ADA82547.1; nef protein [Human immunodeficiency virus 1] QEE94170.1 | 12016,2 |
| 279 | Q9ULG6-5-DECOY | (?)GTFFDDDSKIVVTK(?) | long tail fiber protein distal subunit [Escherichia phage F2] YP_010068895.1; structural protein [Synechococcus phage S-CAM3] AOV58829.1 | 8623,84 |
| 280 | O43768-3-DECOY | (?)RVPVVKESVWKPR(?) | immunity-specific protein Beta371 [Corynephage beta] AAA32187.1;DNA primase [Salmonella phage vB_SenS_ER3] YP_009999602.1; phosphoadenosine-phosphosulfate reductase [Siphoviridae sp.] DAR25076.1 | 1791730 |
| 281 | H3BMZ9-DECOY | (?)VAGSGIWRVLRLPR(?)<br>Query 9 VLRLP 13<br>VL LP<br>Sbjct 135 VLSLP 139 | Chain A, DIPHTHERIA TOXIN [Corynephage beta] 1DDT_A; hypothetical protein H1O17_gp358 [Burkholderia phage BcepSauron] YP_009904736.1; hypothetical protein 9F5_46 [uncultured Caudovirales phage] ASN68733.1 | 5564380 |
| 282 | H0YNG3-DECOY | (?)NLGRLGLVIAIFACK(?) | holin [Myoviridae sp. ctThM1] QGH72592.1 ; hypothetical protein StauST398-4_0059 [Staphylococcus phage StauST398-4] YP_009002815.1; hypothetical protein P9_gp49 [Streptococcus phage P9] YP_001469229.1 | 69614,7 |
| 283 | E7EUM2-DECOY | (?)YLLRRIPLNTLPK(?)<br>Query 2 LLRRIP 7<br>LL IP<br>Sbjct 433 LLPTIP 438 | Chain A, DIPHTHERIA TOXIN [Corynephage beta] 1DDT_A; phosphatidylglycerophosphatase C [Escherichia phage vB_EcoM-606R2] URC08673.1; helix-turn-helix of insertion element transposase [Bacteriophage sp.] UVY12919.1; polymerase basic protein 1, partial [Influenza A virus (A/chicken/Jilin/hn/2003(H5N1))] ABJ80591.1 | 140483 |
| 284 | Q0VDD8-DECOY | (?)AHVFSIHHKVMFR(?) | hypothetical protein 015DV004_49 [Bacillus phage 015DV004] QQO41265.1; structural polyprotein [Picornavirales sp.] QYV43049.1 | 1471070 |
| 285 | E7EUR8-DECOY | (?)HSFPELLLIAHMGK(?) | immunity-specific protein Beta371 [Corynephage beta] AAA32187.1; hypothetical protein [Bacteriophage sp.] DAH40967.1; head completion adaptor [Vibrio phage 1.052.A._10N.286.46.C3] AUR84238.1; | 9818,67 |
| 286 | Q8N157-2-DECOY | (?)RSPNSGHTERRIAK(?)<br>Query 7 HTERRIAK 14<br>HT+ IAK<br>Sbjct 199 HTDQDIAK 206 | immunity-specific protein Beta241 [Corynephage beta] AAA32184.1; replication protein A [Myoviridae sp.] DAY64350.1; Plasmid recombination enzyme [Microviridae sp.] DAN77582.1; site-specific integrase [Mycobacterium phage Adzzy] YP_008409329.1 | 14181 |
| 287 | Q9NUU7-DECOY | (?)RTGGTDCSPMYYDK(?) | putative virion structural protein [Salmonella phage pSal-SNUABM-04] QOC54547.1; phosphoadenosine-phosphosulfate reductase [Siphoviridae sp.] DAK76872.1 | 36216,1 |
| 288 | Q8NGK4-DECOY | (?)SLYLKIKRPCLKR(?)<br>Query 1 SLYLKIK 7<br>SL+ IK<br>Sbjct 528 SLFFEIK 534 | Chain A, DIPHTHERIA TOXIN [Corynephage beta] 1DDT_A; Diphtheria toxin homolog CRM228 P00589.2; flagellar hook-associated protein FlgK [Siphoviridae sp.] DAQ31509.1; hypothetical protein [Bacteriophage sp.] DAL35033.1 | 272679 |
| 289 | Q8N3P4-2-DECOY | (?)TPQSISQDGNAYNLK(?) | tail assembly chaperone protein [Bacteriophage sp.] UWD71945.1; Prophage endopeptidase tail [Bacteriophage sp.] UWG84281.1; Carbapenem-associated resistance protein [Caudovirales sp. ctyaR3] DAF58099.1 | 101737 |
| 290 | Q7Z404-DECOY | (?)EVAECLPLLLPGIMK(?)<br>Query 10 LPGI 13<br>LPGI<br>Sbjct 307 LPGI 310<br>Query 7 PL-----LLPGI 13<br>PL LLP I<br>Sbjct 426 PLPIAGVLLPTI 437<br>Query 2 VAECLPL 8<br>VA+ PL<br>Sbjct 340 VAQAIPL 346 | Chain A, DIPHTHERIA TOXIN [Corynephage beta] 1DDT_A; Diphtheria toxin homolog CRM228 P00589.2; phosphate ABC transporter permease [Acinetobacter phage MD-2021a] CAH1080154.1; terminase large subunit [Aeromonas phage phiAS5] YP_003969300.1 | 824323 |
| 291 | Q13595-DECOY | (?)ILLVSWGTLRQILR(?) | arylsulfotransferase [Bacteriophage sp.] UWF98159.1; helix-turn-helix DNA binding domain protein [Mycobacterium phage phiGD57-1] QPO17290.1; envelope glycoprotein, partial [Human immunodeficiency virus | 6005960 |

| | | | | |
|---|---|---|---|---|
| | | | 1] ADN80780.1; hemagglutinin [Influenza A virus] QOJ98649.1; envelope glycoprotein, partial [Human immunodeficiency virus 1] ABR00968.1 | |
| 292 | P16070-11-DECOY | (?)ARTARRLQISVGVVR(?) | immunity-specific protein Beta241 [Corynephage beta] AAA32184.1; immunity-specific protein Beta371 [Corynephage beta] AAA32187.1; DNA methylase [Mycobacterium phage Madiba] UJE15618.1; Thiamine biosynthesis protein (ThiI) [Siphoviridae sp.] DAH80124.1; DNA-binding domain protein, partial [Bacteriophage sp.] UWI39733.1; tail fiber protein [Staphylococcus phage 6ec] YP_009042548.1; ORF1ab polyprotein, partial [Severe acute respiratory syndrome coronavirus 2] UPU75913.1 | 20770,6 |
| 293 | F5H6V4-DECOY | (?)NIVLVKAISTAVSLKR(?) | Chain A, DIPHTHERIA TOXIN [Corynephage beta] 1DDT_A; immunity-specific protein Beta371 [Corynephage beta] AAA32187.1; holin [Siphoviridae sp.] DAL45079.1; tail tape measure [Siphoviridae sp.] DAO32156.1; nucleocapsid protein, partial [Human respirovirus 3] UTM74195.1 | 3679870 |
| 294 | P78527-2-DECOY | (?)QSIKIVDSGTSVDSMR(?) | immunity-specific protein Beta201 [Corynephage beta] AAA32185.1 ; neck passage structure [Lactococcus phage i0139] YP_009875758.1; DNA polymerase III subunit alpha [Siphoviridae sp.] DAJ88735.1 | 58682,8 |
| 295 | R4GN15-DECOY | (?)MHHSVIVAPSEDMGGCIIIDPPSKIFQYDGWQMI CYNRSCVSHNGQR(?) | immunity-specific protein Beta371 [Corynephage beta] AAA32187.1; Diphtheria toxin homolog CRM228 P00589.2; immunity-specific protein Beta201 [Corynephage beta] AAA32185.1 ; dUTPase [Bacteriophage sp.] UWI15887.1; ead closure knob [Siphoviridae sp.] ; DAZ41614.1 non structural protein, partial [Norovirus Hu/GI/66-1/Tokyo/JPN] BDS00049.1; nonstructural protein [Hepatitis E virus type 3] AOG18220.1; putative chloride channel [Abalone herpesvirus Victoria/AUS/2009] YP_006908742.1; serine protease inhibitor-like protein [Cowpox virus] ADZ30814.1 | 26,405 |
| 296 | Q9UII4-DECOY | (?)SGLKSDFGPLAMNKKK(?) | immunity-specific protein Beta201 [Corynephage beta] AAA32185.1 ; D-tyrosyl-tRNA(Tyr) deacylase [Acinetobacter phage MD-2021a] CAH1082716.1; RNA polymerase sigma factor [Siphoviridae sp. ctXZx16] DAD82881.1; Clp protease [Vibrio phage Marilyn] QKN84422.1 | 3053730 |
| 297 | O75112-2-DECOY | (?)QILLLGEIKLTHLHK(?) | immunity-specific protein Beta201 [Corynephage beta] AAA32185.1 ; GIY-YIG nuclease superfamily protein [Siphoviridae sp.] DAW80734.1; holin [Siphoviridae sp. ctQ0C17] DAD92258.1; DNA topoisomerase [Bacillus phage PBC2] AKQ08441.1; DNA gyrase subunit A [Bacillus phage vB_BanS-Thrax1] UUV46018.1; E7 protein [Human papillomavirus 100] CAW42236.1 | 1126350 |
| 298 | Q9NXH9-2-DECOY | (?)DGDLSRNLIPEQFKK(?) | General substrate transporter:Major facilitator superfamily [Acinetobacter phage MD-2021a] CAH1069775.1; dihydrofolate reductase [Aeromonas phage 25] YP_656433.1; portal protein [Siphoviridae sp.] DAI79382.1; Mn-containing catalase [Siphoviridae sp.] DAE95739.1 | 2555980 |
| 299 | C9JBV8-DECOY | (?)QFISVDSPKDQILNR(?)<br>Query 3 ISVDS 7<br>IS DS<br>Sbjct 504 ISSDS 508 | Chain A, DIPHTHERIA TOXIN [Corynephage beta] 1DDT_A; Diphtheria toxin homolog CRM228 P00589.2; ATP dependent DNA helicase [Siphoviridae sp.] DAU22806.1; lysozyme-like protein [uncultured Mediterranean phage uvMED] BAR13838.1; assembly (protein IV of M13) [Escherichia phage fd] YP_009111306.1; Type II secretion system protein [Inoviridae sp.] DAH58658.1; universal stress protein [Siphoviridae sp.] DAZ31422.1 | 17273,8 |
| 300 | P78563-5-DECOY,Q96JP9-DECOY | (?)MVVQHQILLQVGYLNNGDLHIGTGPTSELVEKA PTWYRECGVCLPGCNR(?)<br>Query 13 YLNNG---DLHI 21<br>Y NG +LH+<br>Sbjct 478 YVGNGVHANLHV 489 | Chain A, DIPHTHERIA TOXIN [Corynephage beta] 1DDT_A; Diphtheria toxin homolog CRM228 P00589.2; immunity-specific protein Beta286 [Corynephage beta] AAA32186.1; putative DNA-binding motif protein [Siphoviridae sp.] DAK99044.1; distal tail protein [Lactobacillus phage ATCCB] QBJ03769.1; holin [Mycobacterium phage Luchador] YP_009203069.1; DNA polymerase III [Siphoviridae sp.] DAH84098.1; polyprotein [Dengue virus type 4] AMW17747.1 | 26,789 |
| 301 | Q9Y4X0-4-DECOY | (?)VVRAPDLLGCAVAKIVMK(?)<br>Query 14 KIVM 17<br>KI M<br>Sbjct 456 KIRM 459 | Chain A, DIPHTHERIA TOXIN [Corynephage beta] 1DDT_A; immunity-specific protein Beta241 [Corynephage beta] AAA32184.1; immunity-specific protein Beta286 [Corynephage beta] AAA32186.1; Diphtheria toxin homolog CRM228 P00589.2; immunity-specific protein Beta201 [Corynephage beta] AAA32185.1 ; endolysin [Lactobacillus phage f-0303] AAQ06663.1; Cpl 1 lysin [Myoviridae sp.] DAQ70471.1; tax, partial [Human T-cell leukemia virus type I] QKE53338.1; large terminase [Siphoviridae sp.] DAG15952.1; | 10596,9 |
| 302 | F8VXU8-DECOY | (?)EIKVRPAAKCLKCALLR(?) | immunity-specific protein Beta201 [Corynephage beta] AAA32185.1 ; Fe-S oxidoreductase [Siphoviridae sp.] DAE95140.1, Rad50 zinc hook motif [Siphoviridae sp.] DAH49148.1, anaerobic ribonucleoside triphosphate reductase [Bacteriophage sp.] DAV36557.1, gag-pol polyprotein, partial [Human immunodeficiency virus 1] BAC02546.1 | 8604,94 |
| 303 | F6UWW1-DECOY | (?)KRAVGLRLAMTILMLAR(?) | immunity-specific protein Beta371 [Corynephage beta] AAA32187.1; PilA, Type IV-A pilus assembly, pilus, ring, membrane channel [Siphoviridae sp.] DAP67763.1, envelope glycoprotein, partial [Human immunodeficiency virus 1] AAQ83732.1, replication protein E1 [Human papillomavirus type 78] BAO74124.1, | 10723,8 |
| 304 | K7EKF8-DECOY | (?)CLMPGCGKVSMLCCTSAK(?) | ORF015 [Staphylococcus phage 55] YP_240470.1, Rep, partial [Human betaherpesvirus 6B] AAC40339.1, orf 58 [Ateline gammaherpesvirus 3] NP_048029.1, REP [Human betaherpesvirus 6A] AVI09293.1 | 136395 |
| 305 | Q70CQ2-DECOY | (?)RKKTLAETSPMLFLCGK(?) | immunity-specific protein Beta241 [Corynephage beta] AAA32184.1 | 702910 |
| 306 | P20929-DECOY | (?)QKMSMDHDTEQVGRCK(?)<br>Query 4 SMDHD 8<br>SM+HD<br>Sbjct 81 SMEHD 85<br>Query 10 EQVG 13<br>EQVG<br>Sbjct 116 EQVG 119 | Chain A, DIPHTHERIA TOXIN [Corynephage beta] 1DDT_A; Diphtheria toxin homolog CRM228 P00589.2; | 149318 |
| 307 | Q07889-DECOY | (?)NLGLSKSAFGLRINKLIR(?)<br>Query 2 LGLS 5<br>LGLS<br>Sbjct 106 LGLS 109 | Chain A, DIPHTHERIA TOXIN [Corynephage beta] 1DDT_A; Diphtheria toxin homolog CRM228 P00589.2 | 7905,17 |
| 308 | Q06210-2-DECOY | (?)LADLPLQPDNLTRISIVK(?) | | 6645,39 |
| 309 | Q8NDW8-5-DECOY | (?)VLLALSLRVESSSELLER(?)<br>Query 11 SSSE 14<br>SSSE<br>Sbjct 494 SSSE 497<br>Query 5 LSLR-VE--SSSE 14<br>LSL E SS E<br>Sbjct 136 LSLPFAEGSSSVE 148 | Chain A, DIPHTHERIA TOXIN [Corynephage beta] 1DDT_A; Diphtheria toxin homolog CRM228 P00589.2 | 31764,3 |
| 310 | G3V1B7-DECOY | (?)SPALSSQATVDRGEKQSLR(?)<br>Query 3 ALSS----QA 8<br>ALSS QA<br>Sbjct 334 ALSSLMVAQA 343<br>Query 2 PALSSQATV 10<br>P LS TV<br>Sbjct 258 PELSELKTV 266 | Chain A, DIPHTHERIA TOXIN [Corynephage beta] 1DDT_A; Diphtheria toxin homolog CRM228 P00589.2; immunity-specific protein Beta286 [Corynephage beta] AAA32186.1; | 158023 |
| 311 | Q5U5Z8-3-DECOY | (?)RNLGDSCNSVGALINMIPR(?) | immunity-specific protein Beta201 [Corynephage beta] AAA32185.1 ; | 251608 |
| 312 | E7EWN1-DECOY | (?)DISQQEWSSMSMDMNLK(?) | immunity-specific protein Beta286 [Corynephage beta] AAA32186.1; | 822877 |
| 313 | Q15375-4-DECOY | (?)DSKEMMQVCCKDENEIK(?) | | 184045 |
| 314 | Q9BQG2-DECOY | (?)TDRMEVVMQDPMDFSCK(?) | immunity-specific protein Beta371 [Corynephage beta] AAA32187.1; | 2816,49 |
| 315 | Q13472-2-DECOY | (?)LLSKQEEKIIHLGAPNAVK(?) | immunity-specific protein Beta241 [Corynephage beta] AAA32184.1; Diphtheria toxin homolog CRM228 P00589.2; | 8373,78 |
| 316 | Q5H909-DECOY | (?)LISQPFHLNKRSIKWPK(?) | immunity-specific protein Beta286 [Corynephage beta] AAA32186.1; | 8469,08 |
| 317 | Q96A54-DECOY | (?)LDIEGVEALPNILIGGRSLK(?) | Diphtheria toxin homolog CRM228 P00589.2; immunity-specific protein Beta286 [Corynephage beta] AAA32186.1 | 118336 |
| 318 | H7BY53-DECOY | (?)ICQSTTLLLMSCTMKKALK(?) | | 550335 |
| 319 | H7BY53-DECOY | (?)EKVGLKSTSAILMAPRVLYR(?) | immunity-specific protein Beta371 [Corynephage beta] AAA32187.1; immunity-specific protein Beta201 [Corynephage beta] AAA32185.1 | 550335 |
| 320 | E9PDH4-DECOY | (?)GDQKCIQRGRPDTIGSEEVK(?) | Chain A, DIPHTHERIA TOXIN [Corynephage beta] 1DDT_A; Diphtheria toxin homolog CRM228 P00589.2; ribonucleoside-triphosphate reductase [Escherichia phage grams] YP_009901833.1 | 7424,82 |
| 321 | Q96T58-DECOY | LLKKNVQAPQFLGSQTGRLVR(?) | immunity-specific protein Beta241 [Corynephage beta] AAA32184.1 | 2822,7 |
| 322 | B7Z7Y1-DECOY | (?)TSIAQFVPHHEFELTKGPRPKCAPLPMAALASVLI LFLAVGSESRFASIPCK(?)<br>Query 24 PLPMA 28<br>PLP A<br>Sbjct 426 PLPIA 430<br>Query 17 VGS 19<br>VGS<br>Sbjct 195 VGS 197<br>Query 8 AS--VLIL-FLAVGSES 21<br>AS VL L F A GS S<br>Sbjct 131 ASRVVLSLPF-AEGSSS 146 | Chain A, DIPHTHERIA TOXIN [Corynephage beta] 1DDT_A; Diphtheria toxin homolog CRM228 P00589.2; putative membrane protein [Staphylococcus phage VB-SauS-SA2] AWY02954.1 | 27782,3 |
| 323 | H0YHL5-DECOY | (?)WWNNKSAMMIGFESCYACAR(?) | WbbJ Acetyltransferase (isoleucine patch superfamily) [uncultured Caudovirales phage] CAB4133703.1 | 5163,2 |
| 324 | P49756-DECOY | (?)IIDSSELNPKSRGLWIVMTDLR(?) | immunity-specific protein Beta241 [Corynephage beta] AAA32184.1 | 326,699 |
| 325 | P49756-DECOY | (?)GKLPLLQNPSPAGAGGVEVVLDVTQK(?) | immunity-specific protein Beta286 [Corynephage beta] AAA32186.1; immunity-specific protein Beta371 [Corynephage beta] AAA32187.1; Diphtheria toxin homolog CRM228 P00589.2 | 8292,01 |
| 326 | F8WAK8-DECOY | (?)AYVGAPPDPAEMMYAGECDLDEHK(?) | immunity-specific protein Beta201 [Corynephage beta] AAA32185.1; Diphtheria toxin homolog CRM228 P00589.2 ; | 6616,48 |
| 327 | F5GWR7-DECOY | (?)ESREAESTRQEVGTGGDGNLQTDLR(?)<br>Query 12 VGTGGDGNLQ 21<br>VG G NL+<br>Sbjct 479 VGNGVHANLH 488<br>Query 21 QTDL 24<br>QT L<br>Sbjct 252 QTAL 255 | DIPHTHERIA TOXIN [Corynephage beta] 1DDT_A; Diphtheria toxin homolog CRM228 P00589.2 ; tail protein [Acinetobacter phage AP22] YP_006383804.1 | 4782,83 |
| 328 | Q9UBB6-DECOY | (?)NRDSKGEWLSQMLEPSNVQDEIN(?) | immunity-specific protein Beta241 [Corynephage beta] AAA32184.1; Diphtheria toxin homolog CRM228 P00589.2 ; site specific tyrosine recombinase [Bacteriophage sp.] DAI17020.1 | 84456,6 |
| 329 | Q96G46-DECOY | (?)MRVHGNDWHIYESAIKVQSAPLR(?) | Diphtheria toxin homolog CRM228 P00589.2; immunity-specific protein Beta286 [Corynephage beta] AAA32186.1 | 61260,5 |
| 330 | F8VVB4-DECOY | (?)IPAAAKQNVPQWPPKKFSQVPPNAK(?)<br>Query 8 NVPQ 11<br>NV Q<br>Sbjct 284 NVAQ 287 | DIPHTHERIA TOXIN [Corynephage beta] 1DDT_A; Diphtheria toxin homolog CRM228 P00589.2; immunity-specific protein Beta286 [Corynephage beta] AAA32186.1 | 975603 |
| 331 | J3QSB2-DECOY | (?)QGLLIQLARGTGLEVIQARGKWHVK(?) | immunity-specific protein Beta286 [Corynephage beta] AAA32186.1; tail protein [Bacteriophage sp.] UWG76518.1 | 35394,2 |
| 332 | A6PVK7-DECOY | (?)MRIGLDAHGFLGVTEASVNIGTGDSSLLARYHL ARDIKPPAIIVAFCMSR(?) | capsid and scaffold protein [Streptococcus phage Javan288] QBX26045.1; terminase [Streptococcus phage IPP24] APD22428.1 | 7923,09 |
| 333 | Q14938-3-DECOY | (?)QTLVLKMQLLAFHSALLVKQGQQLR(?) | immunity-specific protein Beta241 [Corynephage beta] AAA32184.1; Diphtheria toxin homolog CRM228 P00589.2; | 45,382 |

| | | | | |
|---|---|---|---|---|
| 334 | E5RIW1-DECOY | (?)VPVHASPGTDLKSSTLVGPNRAYSQRR(?)<br>Query 12 KSSTLVG 18<br>KS VG<br>Sbjct 474 KSPVYVG 480 | immunity-specific protein Beta286 [Corynephage beta] AAA32186.1; immunity-specific protein Beta201 [Corynephage beta] AAA32185.1; DIPHTHERIA TOXIN [Corynephage beta] 1DDT_A; Diphtheria toxin homolog CRM228 P00589.2 | 15278,9 |
| 335 | Q4AC94-4-DECOY | (?)VFEGESSFQECVCHVSRIQALDYCPVKPMLKAPA AEDPTCSLSNARMAEEAATTR(?)<br>Query 3 EGESS 7<br>EG SS<br>Sbjct 142 EGSSS 146<br>Query 16 AR--MAEEA 22<br>AR M E A<br>Sbjct 220 ARVEMVEAA 228 | DIPHTHERIA TOXIN [Corynephage beta] 1DDT_A; immunity-specific protein Beta241 [Corynephage beta] AAA32184.1; immunity-specific protein Beta201 [Corynephage beta] AAA32185.1 | 31,029 |
| 336 | P53804-3-DECOY | (?)KMMDIGVIEMGDALHADLKDDSPMLDHEPAGN PLAHTLHCAQETAEIEKESGNSNEGHQCTNSK(?)<br>Query 2 MMDIGVI 8<br>M +IG I<br>Sbjct 181 MNEIGAI 187<br>Query 21 EKESGNSNE 29<br>EK SNE<br>Sbjct 497 EKI--HSNE 503 | immunity-specific protein Beta371 [Corynephage beta] AAA32187.1; DIPHTHERIA TOXIN [Corynephage beta] 1DDT_A; Diphtheria toxin homolog CRM228 P00589.2 | 700,643 |
| 337 | Q8WZ42-13-DECOY | (?)CTQTLPNTGPIGEDTKEDESLCKVMSQDEWGDC FNCIPRGVSDSDISMYYTQSYISQGVGSTENEANLS | DIPHTHERIA TOXIN [Corynephage beta] 1DDT_A; Receptor Binding Protein [Siphoviridae sp.] DAM26724.1; lysozyme [Enterocloster phage CB457P1] WAX11085.1 | 18389,2 |
| 338 | Q8N1G0-2-DECOY | (?)QSPTVAEDLCEAHEEVYEFASAVKGMVLGTVQ MLNINPYEQGVDVIFSMAYLGSDRQSFTKSLRPPIR<br>Query 8 INPYEQ 13<br>IN EQ<br>Sbjct 150 INNWEQ 155<br>Query 6 LNINPYE----QGVDVIFSM 21<br>L IN +E G D M<br>Sbjct 163 LEIN-FETRGKRGQD---AM 178 | DIPHTHERIA TOXIN [Corynephage beta] 1DDT_A 232-250; terminase large subunit [Vibrio phage 275E43-1] CAH9015435.1; DNA polymerase [Staphylococcus phage VB-SauS-SA2] AWY02983.1; holin/anti-holin [Escherichia phage Lambda] YP_001551775.1 | 462,07 |
| 339 | Q86UR5-5-DECOY | (?)LLLLLRVLCSEAGPENSESFFYEGAEPLSATGKM GLTILLRLILEPGEALTSVAR(?) | Putative ATP dependent Clp protease [Siphoviridae sp.] DAN90103.1; RNA polymerase beta subunit [Pseudomonas phage Psa21] YP_010347754.1 PLP-dependent enzyme [Siphoviridae sp.] DAL56212.1 | 1236,64 |
| 340 | E7EQT2-DECOY | (?)LHSLCTMMSKSDSIPSTTVMARARPNQCIPMPSG AAKFCGTMPTGSNAEVIGCGTVPR(?) | PRTRC system ThiF family protein [Bacteriophage sp.] DAD60148.1; phage tail tape measure protein-like protein [Virus Rctr71] APU89196.1; Putative Head Tail Connector Protein [Bacteriophage sp.] DAK94203.1 | 52,541 |
| 341 | Q96T58-DECOY | (?)SPVELVTVLNEVSSITPEEAGVMLWCSGIQLSGR VPESSSSKPCREMTCCTNTFNSVNVFELEANMYCF VFRLK(?)<br>Query 10 NEVSS 14<br>NE+SS<br>Sbjct 502 NEISS 506<br>Query 14 SITPEEAGVMLWCSGI 29<br>SI P VM GI<br>Sbjct 305 SILPGIGSVM----GI 316<br>Query 4 ELVTV 8<br>EL TV<br>Sbjct 262 ELKTV 266<br>Query 8 ESSSS 12<br>E SSS<br>Sbjct 142 EGSSS 146 | Chain A, DIPHTHERIA TOXIN [Corynephage beta] 1DDT_A; lysin [Streptococcus phage Javan272] QBX25795.1; exonuclease [Salmonella phage 9NA] YP_009101237.1; cell division protein [Siphoviridae sp.] DAS99818.1; Clp protease-like protein [Streptococcus phage Javan139] QBX14465.1; hypothetical protein [Bacteriophage sp.] DAH86615.1 | 2822,7 |
| 342 | P02751-3-DECOY | (?)ILVAFENK(?)<br>Query 1 ILVAFEN 7<br>ILVAFEN<br>Sbjct 260 ILVAFEN 266 | tail length tape measure protein [Pseudomonas phage PS-1]<br>Sequence ID: YP_009222841.1 | **1,37E12** |
| 343 | Q9UPP1-2-DECOY | (?)VLGLKNMR(?)<br>Query 1 VLGLK 5<br>VL LK<br>Sbjct 91 VLALK 95 | Chain A, DIPHTHERIA TOXIN [Corynephage beta] 1DDT_A | 201956 |
| 344 | Q9P1Z9-4-DECOY | (?)SVLNDYMPQCALALALVKPGQKICLPGDTESRD MSELVEGAICMMLLTIIRHGHK(?)<br>Query 6 GDTESR 11<br>GD SR<br>Sbjct 128 GDGASR 133 | Chain A, DIPHTHERIA TOXIN [Corynephage beta] 1DDT_A | 20123,5 |
| 345 | Q8WXH0-2-DECOY | (?)IKSARENTK(?)<br>Query 1 IK-SARENT 8<br>IK A ENT<br>Sbjct 418 IKITA-ENT 425 | Chain A, DIPHTHERIA TOXIN [Corynephage beta] 1DDT_A | 146444 |
| 346 | O15321-DECOY | (?)DLGPAEIKQK(?)<br>**Beta241**<br>Query 5 AEIKQ 9<br>AEIKQ<br>Sbjct 5 AEIKQ 9 | immunity-specific protein Beta241 [Corynephage beta] AAA32184.1 | 137368 |
| 347 | Q9UGN4-DECOY | (?)TAAMLEKILR(?)<br>Query 1 TAAMLEK 7<br>TA LEK<br>Sbjct 293 TADNLEK 299 | Chain A, DIPHTHERIA TOXIN [Corynephage beta] 1DDT_A | 477348 |
| 348 | I3L136-DECOY | (?)QDAKTREIPGK(?)<br>Query 8 IPGK 11<br>IPGK<br>Sbjct 437 IPGK 440 | Chain A, DIPHTHERIA TOXIN [Corynephage beta] 1DDT_A | 357102 |
| 349 | Q01484-DECOY | (?)LSLTKLQEGAAK(?)<br>Query 1 LSLT 4<br>LSLT<br>Sbjct 108 LSLT 111 | Chain A, DIPHTHERIA TOXIN [Corynephage beta] 1DDT_A | 78260,6 |
| 350 | Q6ZRS2-3-DECOY | (?)HEKCINLELVKVR(?)<br>Query 4 CINLE 8<br>CINL+<br>Sbjct 226 CINLD 230 | Chain A, DIPHTHERIA TOXIN [Corynephage beta] 1DDT_A | 173714 |
| 351 | Q9P0M6-DECOY | (?)MAEAGRAEGLSVAITIR(?)<br>Query 1 MAEA--G----RAEGLSV 12<br>MA+A G R SV<br>Sbjct 182 MAQACAGNRVRR----SV 195 | Chain A, DIPHTHERIA TOXIN [Corynephage beta] 1DDT_A | 1699950 |
| 352 | P0C0L4-2-DECOY | (?)TDSRLALKGDMMTVPER(?)<br>Query 5 LALKGD 10<br>LALK D<br>Sbjct 92 LALKVD 97 | Chain A, DIPHTHERIA TOXIN [Corynephage beta] 1DDT_A | 2019460 |
| 354 | G3V1B7-DECOY | (?)SPALSSQATVDRGEKQSLR(?)<br>Query 3 ALSS----QA 8<br>ALSS QA<br>Sbjct 334 ALSSLMVAQA 343<br><br>Query 2 PALSSQATV 10<br>P LS TV<br>Sbjct 258 PELSELKTV 266 | Chain A, DIPHTHERIA TOXIN [Corynephage beta] 1DDT_A | **1,27E12** |
| 355 | F8WA11-DECOY | (?)LPVSPAKSAPSEELVRVAAGER(?)<br>**Beta241**<br>Query 18 AAGE 21<br>AAGE<br>Sbjct 123 AAGE 126 | immunity-specific protein Beta241 [Corynephage beta] AAA32184.1 | 427609 |
| 356 | Q8TCU4-DECOY | (?)LVPQDRQLCVITASCGFQALRMLSEIQMGPAKL QECDGTK(?)<br>Query 23 LS---EI 26<br>LS EI<br>Sbjct 159 LSVELEI 165 | Chain A, DIPHTHERIA TOXIN [Corynephage beta] 1DDT_A | 217605 |
| 357 | Q86YD3-5-DECOY | (?)CPIGESLVVITLRAAEIEMSLALQGSMGAWMSGF QVLYDLPILELLDFYLFFKVMK(?)<br>Query 4 GESLVVITLRAAE 16<br>G S VV L AE<br>Sbjct 130 GASRVVLSLPFAE 142<br>Query 17 IEMSL 21<br>IE SL<br>Sbjct 217 IE-SL 220 | Chain A, DIPHTHERIA TOXIN [Corynephage beta] 1DDT_A | 277953 |
| 358 | Q9Y6K9-2-DECOY | (?)CGLSSPIGLCTEVEGQILLISSQSVLRQIFELSVMQ CGDPIGPEFKPSRCIERYGIR(?)<br>**Beta201**<br>Query 27 RQIF 30<br>RQIF<br>Sbjct 82 RQIF 85<br>Query 12 EVEGQI 17<br>EV GQ+<br>Sbjct 44 EVPGQV 49 | immunity-specific protein Beta201 [Corynephage beta] AAA32185.1 | 61232,3 |
| 359 | Q96DY2-3-DECOY | (?)TKNSTQTALQGKLGATIVTHCSFGKLFENAPVYT FGLTIIADQTMSLTYGITGEVMPVCVIYSFGR(?)<br>Query 6 QTALQ 10<br>QTAL+<br>Sbjct 252 QTALE 256 | Chain A, DIPHTHERIA TOXIN [Corynephage beta] 1DDT_A | 68060,1 |
| 360 | J3KS19-DECOY | (?)DQELQKIMHEEPEAGEPSLRYIINVPVDFSDVFQ GELLEFLGANAVPETNLILHVIMTVTKGR(?)<br>Query 7 FLGAN----AVPETNLILHVI 23<br>F GAN AV N + +VI<br>Sbjct 273 FAGANYAAWAV---N-VAQVI 289<br>**Beta241**<br>Query 4 LQKIMHE 10<br>LQ H+<br>Sbjct 79 LQSMEHD 85 | Chain A, DIPHTHERIA TOXIN [Corynephage beta] 1DDT_A;<br>immunity-specific protein Beta241 [Corynephage beta] AAA32184.1;<br>immunity-specific protein Beta371 [Corynephage beta] AAA32187.1;<br>immunity-specific protein Beta201 [Corynephage beta] AAA32185.1 | 232101 |
| 361 | Q8N531-DECOY | (?)SSSVANLEHIPMPRPQTLSSSLIAQKERSIIQKDSN DLFFYLLASVLSVHTADPMVYDNVVFR(?)<br>Query 18 LSSSLIAQ 25<br>LSS ++AQ<br>Sbjct 335 LSSLMVAQ 342<br>Query 1 SSSVANLEHI 10<br>SSSV E I<br>Sbjct 144 SSSV---EYI 150<br>Query 10 IPM 12<br>IP+<br>Sbjct 344 IPL 346<br>Query 21 SLIAQKE 27<br>SL KE<br>Sbjct 219 SL---KE 222<br>**Beta371**<br>Query 7 ASVLSV 12<br>ASVL V<br>Sbjct 306 ASVLGV 311<br>**Beta201**<br>Query 6 LASVLSV---HTA 15<br>L S+ SV TA<br>Sbjct 108 LDSIISVPSQDTA 120 | Chain A, DIPHTHERIA TOXIN [Corynephage beta] 1DDT_A;<br>immunity-specific protein Beta371 [Corynephage beta] AAA32187.1;<br>immunity-specific protein Beta201 [Corynephage beta] AAA32185.1 | 152688 |
| 362 | Q9BTC0-DECOY | (?)LQATNFYGEVFPCLELTVFAIGKIPVVILEHQHVR ASYSSLSKMSSLPNGLLMGGQPVSPAGSCNGR(?) | Chain A, DIPHTHERIA TOXIN [Corynephage beta] 1DDT_A;<br>immunity-specific protein Beta201 [Corynephage beta] AAA32185.1 | 29668,8 |

| | | | | |
|---|---|---|---|---|
|  |  | Query 10 KMSSLPN 16<br>KMS PN<br>Sbjct 229 KMSESPN 235<br>Query 6 SSLS 9<br>SSLS<br>Sbjct 197 SSLS 200<br>**Beta201**<br>Query 2 NG----LLMGG 8<br>NG LL GG<br>Sbjct 17 NGSPAFLLKGG 27 |  |  |
| 363 | Q6ZRS4-2-DECOY | (?)LFPANWSSKMRPHTLILSKYLSCAPSGMPACLEFVWLVMGPVDFGWFLELSNPKIEYTTLIR(?)<br>Query 13 HTLILSKYLS 22<br>HT + SK LS<br>Sbjct 520 HTKVNSK-LS 528<br>Query 15 VDFG 18<br>VD G<br>Sbjct 351 VDIG 354 | Chain A, DIPHTHERIA TOXIN [Corynephage beta] 1DDT_A | 28339,3 |
| 364 | F5GWS0-DECOY | (?)IDTQAFIGDQIRQSNAKAVLYLADEWCVESFLLPPTLYVFELVTTIIAVIMLIPDPLLPASYGLK(?)<br>Query 14 SNAKAVLYL 22<br>S KA YL<br>Sbjct 239 SEEKAKQYL 247<br>Query 25 EW 26<br>+W<br>Sbjct 136 QW 137<br>Query 23 IAVIML--IP 30<br>IA ++L IP<br>Sbjct 429 IAGVLLPTIP 438 | Chain A, DIPHTHERIA TOXIN [Corynephage beta] 1DDT_A | 36557,3 |
| 365 | Q13574-5-DECOY | QAHEEGLLCIDGPMGIGLGGVKPVQPAKALGTMCDDDAPNQCPALKCELSVVIVYNELVALYLHKIVPK<br>Query 1 AKAL 4<br>AKAL<br>Sbjct 156 AKAL 159<br>Query 18 ALKCEL 23<br>AL EL<br>Sbjct 158 ALSVEL 163<br>**Beta286**<br>Query 21 VKPV 24<br>VKPV<br>Sbjct 220 VKPV 223 | Chain A, DIPHTHERIA TOXIN [Corynephage beta] 1DDT_A;<br>immunity-specific protein Beta286 [Corynephage beta] AAA32186.1 | 96480 |
| 366 | E7ERL6-DECOY | (?)SKDMNGIVKISVGPEIEVCEPVAFIMMVQMLPLLSTHVVFLQMSEVDENDMNSPNEGIPALKGLLSR(?)<br>Query 22 VAF 24<br>VAF<br>Sbjct 489 VAF 491<br>Query 11 SVGPEI 16<br>SV EI<br>Sbjct 160 SVELEI 165<br>Query 26 MM 27<br>+M<br>Sbjct 338 LM 339<br>**Beta201**<br>Query 11 GIPALKG 17<br>GI LKG<br>Sbjct 27 GITDLKG 33 | Chain A, DIPHTHERIA TOXIN [Corynephage beta] 1DDT_A;<br>immunity-specific protein Beta201 [Corynephage beta] AAA32185.1 | 162079 |
| 367 | K7EIR0-DECOY | (?)NVILIERDQKEHACLFQGLLEHSILALHDQECLQESLITLANAAVSSTQLFPIAHAVLMMLHPDSAK(?)<br>Query 20 LEHSILAL 27<br>LE AL<br>Sbjct 297 LEKTTAAL 304<br>Query 13 IAHAVLM 19<br>IA LM<br>Sbjct 333 IALSSLM 339<br>**Beta241**<br>Query 8 LH-DQE---CLQESLITLANA 24<br>LH DQ+ L+E L NA<br>Sbjct 198 LHTDQDIAKALEETL----NA 214 | Chain A, DIPHTHERIA TOXIN [Corynephage beta] 1DDT_A;<br>immunity-specific protein Beta241 [Corynephage beta] AAA32184.1 | 150810 |
| 368 | B4DWL0-DECOY | (?)KGTAHVSNPDNLWKAVAGANFLAIDHLACPLFMEIQASALATPGFADLKVNQLWAHMVSPAWAGNTGAAGALPTHR(?)<br>Query 17 AGANF 21<br>AGAN+<br>Sbjct 274 AGANY 278<br>Query 15 LFMEI 19<br>LF EI<br>Sbjct 529 LFFEI 533<br>Query 20 QASAL 24<br>QA AL<br>Sbjct 155 QAKAL 159<br>Query 19 IQ 20<br>IQ<br>Sbjct 31 IQ 32<br>Query 27 PG 28<br>PG<br>Sbjct 382 PG 383<br>Query 13 SPAWAGN 19<br>SP GN<br>Sbjct 475 SPVYVGN 481<br>**Beta371**<br>Query 14 PAWAG 18<br>P WAG<br>Sbjct 277 PVWAG 281<br>**Diphtheria toxin, partial**<br>Query 3 AWAGN----------------TGAAGALP 15<br>AWA N TGA LP<br>Sbjct 308 AWAVNVGPVIESETADNLEKTTGALSILP 336 | Chain A, DIPHTHERIA TOXIN [Corynephage beta] 1DDT_A;<br>immunity-specific protein Beta371 [Corynephage beta] AAA32187.1;<br>diphtheria toxin, partial [Corynephage beta] Sequence ID: ABU25232.1 | 4371,08 |
| 369 | F5H0F9-DECOY | NSLPMKPSPK<br>Query 3 LPMKPSP 9<br>LPMKP P<br>Sbjct 95 LPMKPNP 101 | holin protein [Caudoviricetes sp.]DAE42861.1 | **1,82E12** |
| 370 | Q8IZP9-4-DECOY | AEDDKEAFHSLGKVSALPTVLVSLLEKLWGTVVCPLSTLQYPTVMLTTR<br>Query 11 LGKVSAL 17<br>L KV AL<br>Sbjct 88 LTKVLAL 94<br>**Beta201**<br>Query 13 YPT-------VMLTTR 21<br>YPT M TR<br>Sbjct 67 YPTKEMPMEKAMRKTR 82 | Chain A, DIPHTHERIA TOXIN [Corynephage beta] 1DDT_A; immunity-specific protein Beta201 [Corynephage beta] AAA32185.1 | **1,12E12** |

**Supplementary Table S1: Putatively *Corynephage*-derived "DECOY"-Peptides identified within the immunoreactive 16kDa *C. simulans*-WB-Band**

The 16 kDa-WB-band of a freshly generated heat-treated *C. simulans* extract was cut out and analyzed by proteomic analyses of in-blot-trypsin-digested proteins by LC-MS/MS. Analyzes were performed using Scaffold Software™ (version 5.2.1), showing data from a Peptide Report. Filters excluded human proteins and other known contaminating proteins. Except peptide 231, only peptides declared as "DECOY"-peptides, which revealed no hits in the Universal Protein Resource (UniProt) databank, are shown. Since it was hypothesized that these peptides originate from phages, alignments for the peptides were performed after BLASTP (protein-protein BLAST®)-search towards viruses (taxid:10239), *Corynephage beta* (taxid:10703), and also *Corynebacterium diphtheriae* (taxid:1717, highlighted in red). Note the high abundance of peptides with similarity to diphtheria toxin, its variants, and a homolog (highlighted in yellow). An alignment of selected immunopeptides with Chain A, DIPHTHERIA TOXIN[*Corynephage beta*]1DDT_A , variants, and diphtheria toxin-homologs is also shown. Immunopeptide no. 231 is identical with Testis associated actin remodelling kinase 2, TESK2 (highlighted in turquoise), and is almost identical with a diphtheria toxin variant.

Immunopeptide no. 136 reveals similarity to an iron ABC transporter and a putative siderophore transport system in *Corynebacterium diphtheria*, and other *Corynebacterium taxa*.

Immunopeptides with extraordinary abundance ($10^7$ – $>10^{12}$ TICs, peptides numbers 8, 25, 26, 72, 73, 78, 79, 85, 135-138, 210, 230, 231, and 238-240, 342, 354, 369-370) are shown in bold.
