## Supplementary material for "Discovery of Natural Bispecific Antibodies: Is Psoriasis Induced by a Toxigenic *Corynebacterium simulans* and Maintained by CIDAMPs as Autoantigens?": Suppl Table S2

### Supplementary Table S2

| Peptide number | Protein accession number | Sequence; BLAST® search for Corynephage beta proteins (taxid:10703) | BLASTP (protein-protein BLAST®)-search *Corynephage beta* (taxid:10703) | Abundance (TIC) |
|---|---|---|---|---|
| 1 | tr\|A4QGJ2\|A4QGJ2_CORGB-DECOY | TLEAIR | | 50.131,4 |
| 2 | tr\|I7H9P5\|I7H9P5_CORUL-DECOY | HEGTIR | | 417515 |
| 3 | tr\|B1VGJ2\|B1VGJ2_CORU7-DECOY | AVEFLR | | 21.074,3 |
| 4 | tr\|C0DZG1\|C0DZG1_9CORY-DECOY | PEILER | | 8.492,4 |
| 5 | tr\|E9SZK1\|E9SZK1_COREQ-DECOY | LEIAGTR | | 90.732,7 |
| 6 | tr\|E2MYN9\|E2MYN9_CORAY-DECOY | DWELAK<br>Query 1 DWELAK 6<br>+WE AK<br>Sbjct 152 NWEQAK 157 | Chain A, DIPHTHERIA TOXIN [Corynephage beta] 1DDT_A; Diphtheria toxin homolog CRM228 P00589.2 | 13.929,4 |
| 7 | tr\|S4X9Q6\|S4X9Q6_9CORY-DECOY | AMGIRAK | | 3.137,6 |
| 8 | tr\|C0VT52\|C0VT52_9CORY-DECOY | LIVGLQK | | 10.295,5 |
| 9 | tr\|D9QDV2\|D9QDV2_CORP2-DECOY | WAEAAAR | | 88.530,6 |
| 10 | tr\|M1NNX3\|M1NNX3_9CORY-DECOY | ADILAGAR<br>Query 1 ADILAGA 7<br>A IL GA<br>Sbjct 7 ASILIGA 13 | immunity-specific protein Beta371 [Corynephage beta] AAA32187.1; Diphtheria toxin homolog CRM228 P00589.2 | 9.289,1 |
| 11 | tr\|Q8FR48\|Q8FR48_COREF-DECOY | ANMDVAR | | 9.945,7 |
| 12 | tr\|H2GPS6\|H2GPS6_CORDB-DECOY | GPPEIQR | | 30.157,8 |
| 13 | tr\|S4XH24\|S4XH24_9CORY-DECOY | AAMIVRK | | 83.433,5 |
| 14 | tr\|F8E2W4\|F8E2W4_CORRG-DECOY | AIADDFR | | 127988 |
| 15 | tr\|C3PIL7\|C3PIL7_CORA7-DECOY | VGIDNFR | | 94.633,4 |
| 16 | tr\|H2GFN4\|H2GFN4_CORDN-DECOY | ALSYITR | | 1022520 |
| 17 | tr\|C8RVT8\|C8RVT8_CORJE-DECOY | ADDKRVK<br>Query 3 DKRV 6<br>DKRV<br>Sbjct 44 DKRV 47 | immunity-specific protein Beta371 [Corynephage beta] AAA32187.1 | 336111 |
| 18 | tr\|G0CVL9\|G0CVL9_CORUB-DECOY | GVPGGFGSR<br>Query 1 GVPGG----FG 7<br>GV GG FG<br>Sbjct 40 GVMGGIASFFG 50 | immunity-specific protein Beta241 [Corynephage beta] AAA32184.1 | 195936 |
| 19 | tr\|C0E4C1\|C0E4C1_9CORY-DECOY | HEVIALR | | 39.970,8 |
| 20 | tr\|E9SXW6\|E9SXW6_COREQ-DECOY | HIAAVLAR<br>Query 2 IAAVL 6<br>IA VL<br>Sbjct 429 IAGVL 433 | Chain A, DIPHTHERIA TOXIN [Corynephage beta] 1DDT_A; immunity-specific protein Beta286 [Corynephage beta] AAA32186.1 | 8.128,8 |
| 21 | tr\|C2CRE6\|C2CRE6_CORST-DECOY | SLQSLDLK<br>Query 5 LDLK 8<br>LDLK<br>Sbjct 271 LDLK 274 | immunity-specific protein Beta286 [Corynephage beta] AAA32186.1; immunity-specific protein Beta371 [Corynephage beta] AAA32187.1 | 30.395,6 |
| 22 | sp\|Q6NJ04\|DNAE2_CORDI-DECOY | QMLEIGAK<br>Query 2 MLEIGA 7<br>M EIGA<br>Sbjct 181 MNEIGA 186 | immunity-specific protein Beta371 [Corynephage beta] AAA32187.1 | 11.746,5 |
| 23 | tr\|H2I1T6\|H2I1T6_CORDW-DECOY | MNKQELK | | 5.986,56 |
| 24 | tr\|C0XUY5\|C0XUY5_9CORY-DECOY | YIELLTR<br>Query 1 YIEL 4<br>Y+EL<br>Sbjct 261 YVEL 264 | immunity-specific protein Beta371 [Corynephage beta] AAA32187.1 | 11.774,8 |
| 25 | tr\|G2EK45\|G2EK45_CORGT-DECOY | AMNDITTR | | 57.064,4 |
| 26 | tr\|D5UTB6\|D5UTB6_TSUPD-DECOY | FESDKLASKEMSR | | 8.130,5 |
| 27 | tr\|V6V4E2\|V6V4E2_CORUL-DECOY | ILIPRWR | | 39.711,2 |
| 28 | tr\|U7K5H8\|U7K5H8_9CORY-DECOY | GGVYAESFE | | 41.613,1 |
| 29 | tr\|Q6NI43\|Q6NI43_CORDI-DECOY | LQTGPPAER | | 793166 |
| 30 | tr\|S5TGD4\|S5TGD4_9CORY-DECOY | SPIEAGLGAR<br>Query 6 GLGAR 10<br>GL AR<br>Sbjct 217 GLEAR 221 | immunity-specific protein Beta241 [Corynephage beta] AAA32184.1 | 35.715,2 |
| 31 | tr\|S2Z3K6\|S2Z3K6_9CORY-DECOY | RSDVAEAPK | | 99.538,7 |
| 32 | tr\|B1VFD2\|B1VFD2_CORU7-DECOY | IEIDTINR | | 43.214,4 |
| 33 | tr\|X5DUC4\|X5DUC4_9CORY-DECOY | AAGPFLEIR | | 13.258,5 |
| 34 | tr\|E4WHL8\|E4WHL8_RHOE1-DECOY | NEMGIVGMK | | 87.271,4 |
| 35 | tr\|G0CP56\|G0CP56_CORUL-DECOY | AGYTQQALK | | 35.177,1 |
| 36 | tr\|U7KLF8\|U7KLF8_9CORY-DECOY | MDLTKLIR | | 46.341,2 |
| 37 | tr\|D5UP96\|D5UP96_TSUPD-DECOY | TIIFEITR | | 26.667,0 |
| 38 | tr\|C8NS66\|C8NS66_COREF-DECOY | LVLLSSPER | | 333117 |
| 39 | tr\|M1UJB9\|M1UJB9_9CORY-DECOY | LDILLDTAR<br>Query 5 LDTA 8<br>LDTA<br>Sbjct 87 LDTA 90 | immunity-specific protein Beta371 [Corynephage beta] AAA32187.1 | 33.560,7 |
| 40 | tr\|C0XSN0\|C0XSN0_9CORY-DECOY | EGSARSKTAR | | 33.027,0 |
| 41 | tr\|G7HUI9\|G7HUI9_9CORY-DECOY | LAAVDDVLQK<br>Query 1 LAAVDD 6<br>LA VD+<br>Sbjct 79 LAKVDE 84 | immunity-specific protein Beta286 [Corynephage beta] AAA32186.1 | 229679 |
| 42 | tr\|C2GK55\|C2GK55_9CORY-DECOY | TGFADVLEAR<br>Query 4 ADVLE 8<br>AD LE<br>Sbjct 294 ADNLE 298 | Chain A, DIPHTHERIA TOXIN [Corynephage beta] 1DDT_A; immunity-specific protein Beta241 [Corynephage beta] AAA32184.1; Diphtheria toxin homolog CRM228 P00589.2 | 23.104,2 |
| 43 | tr\|E2MTN2\|E2MTN2_CORAY-DECOY | LAATSSLFLR<br>Query 2 AATSSLFL 9<br>AA FL<br>Sbjct 91 AAQDAVFL 98 | immunity-specific protein Beta286 [Corynephage beta] AAA32186.1 | 47.265,5 |
| 44 | tr\|G2EIC2\|G2EIC2_CORGT-DECOY | ETESACLER | | 75.179,5 |
| 45 | tr\|H2H4N5\|H2H4N5_CORDD-DECOY | DVKEMGEGSR | | 30.082,9 |
| 46 | tr\|C2CMH5\|C2CMH5_CORST-DECOY | TIQELQPLGK | | 33.472,3 |
| 47 | tr\|D5NVS7\|D5NVS7_CORAM-DECOY | LVPDSWEKR<br>Query 4 DSWE 7<br>DSW+<br>Sbjct 273 DSWD 276 | immunity-specific protein Beta371 [Corynephage beta] AAA32187.1 | 151955 |
| 48 | tr\|M1UES1\|M1UES1_9CORY-DECOY | GGGDGQMLWAK | | 33.660,2 |
| 49 | tr\|G4QPV7\|G4QPV7_CORPS-DECOY | AISAFDGESLK<br>Query 8 ESLK 11<br>ESLK<br>Sbjct 218 ESLK 221<br><br>Query 3 SAFD 6<br>SAFD<br>Sbjct 86 SAFD 89 | Chain A, DIPHTHERIA TOXIN [Corynephage beta] 1DDT_A;<br><br>immunity-specific protein Beta201 [Corynephage beta] AAA32185.1 | 40.881,2 |
| 50 | tr\|S2YAM4\|S2YAM4_9CORY-DECOY | GSAIGLGAPELR | | 18.946,8 |
| 51 | tr\|I0LHC2\|I0LHC2_CORGK-DECOY | LMDMRLALR | | 14.963,5 |
| 52 | tr\|H2GCX3\|H2GCX3_CORD2-DECOY | VFDQMNEVR | | 98.448,6 |
| 53 | tr\|H2H4F2\|H2H4F2_CORDD-DECOY | DADGLGQQPVR | | 7.855,9 |
| 54 | tr\|E4W9S8\|E4W9S8_RHOE1-DECOY | AFSGSEQVAIR | | 72.030,0 |
| 55 | tr\|R0JIG6\|R0JIG6_CORCT-DECOY | TGLSNMSAILR<br>Query 3 LSNMSAI 9<br>LS AI<br>Sbjct 297 LSPSNAI 303 | immunity-specific protein Beta371 [Corynephage beta] AAA32187.1 | 5.230,7 |
| 56 | tr\|I7HCG8\|I7HCG8_CORUL-DECOY | ACETTVALLLR<br>Query 4 TTVAL 8<br>TT AL<br>Sbjct 300 TTAAL 304 | Chain A, DIPHTHERIA TOXIN [Corynephage beta] 1DDT_A; immunity-specific protein Beta286 [Corynephage beta] AAA32186.1; Diphtheria toxin homolog CRM228 P00589.2 | 13.036,5 |
| 57 | tr\|C0E262\|C0E262_9CORY-DECOY | QDTIVISMLR | | 12.363,7 |
| 58 | tr\|U7KBF1\|U7KBF1_9CORY-DECOY | ATSDMANLILK | | 73.536,8 |
| 59 | tr\|D5UND1\|D5UND1_TSUPD-DECOY | KNTLEFSTLR | | 53.116,5 |
| 60 | tr\|G0CR63\|G0CR63_CORUL-DECOY | NHCNVALLGLR<br>Query 6 ALLG 9<br>ALLG<br>Sbjct 13 ALLG 16 | Diphtheria toxin homolog CRM228 P00589.2 | 94.061,2 |
| 61 | tr\|E4WIU1\|E4WIU1_RHOE1-DECOY | GALEDIQSMIK | | 36.934,4 |
| 62 | tr\|C6R6B6\|C6R6B6_9CORY-DECOY | EDLTLNLADPK | | 34.372,6 |
| 63 | tr\|U3GWZ4\|U3GWZ4_9CORY-DECOY | LEDAQDITALR<br>Query 3 DAQDITA 9<br>DA TA<br>Sbjct 202 DAKAETA 208 | immunity-specific protein Beta371 [Corynephage beta] AAA32187.1 | 76.631,3 |
| 64 | tr\|H2GXZ8\|H2GXZ8_CORD7-DECOY | KAGEFTEPLIR<br>Query 6 TEPL 9<br>TEPL<br>Sbjct 111 TEPL 114 | Chain A, DIPHTHERIA TOXIN [Corynephage beta] 1DDT_A; Diphtheria toxin homolog CRM228 P00589.2 | 28.805,3 |
| 65 | tr\|U7LS26\|U7LS26_9CORY-DECOY | ILQMSLAEISR | | 22.820,8 |
| 66 | tr\|E4WJ61\|E4WJ61_RHOE1-DECOY | LIATLDLADYR<br>Query 6 DLAD 9<br>DLAD<br>Sbjct 170 DLAD 173 | immunity-specific protein Beta241 [Corynephage beta] AAA32184.1; | 340890 |
| 67 | tr\|U7L1M5\|U7L1M5_9CORY-DECOY | SFHGAGHLVALR | | 39.605,3 |
| 68 | sp\|Q8NP53\|FTSK_CORGL-DECOY | LMGGGDPMILNR<br>Query 3 GGGDPMI 9<br>GG MI<br>Sbjct 50 GGNGAMI 56 | immunity-specific protein Beta241 [Corynephage beta] AAA32184.1 | 59.265,0 |
| 69 | tr\|U7KXL6\|U7KXL6_9CORY-DECOY | FGIERNAARGGK<br>Query 4 ERNAA 8<br>ER+AA<br><br>Sbjct 114 ERDAA 118 | immunity-specific protein Beta241 [Corynephage beta] AAA32184.1 | 7.616,4 |
| 70 | tr\|B1VHK5\|B1VHK5_CORU7-DECOY | QQTVLSPTATLK<br>Query 4 VLSPT 8<br>VL PT<br>Sbjct 432 VLLPT 436 | Chain A, DIPHTHERIA TOXIN [Corynephage beta] 1DDT_A; Diphtheria toxin homolog CRM228 P00589.2 | 12348 |
| | tr\|E4WHE4\|E4WHE4_RHOE1-DECOY | SPYDYVVNPDGALDAK | | 13.480,6 |
| 71 | tr\|B0RFP2\|B0RFP2_CLAMS-DECOY | KLIQGQEQEIK<br>Query 4 QGQEQE 9<br>QG QE<br>Sbjct 9 QGRMQE 14 | immunity-specific protein Beta241 [Corynephage beta] AAA32184.1 | 59.267,1 |
| 72 | tr\|U2GKZ7\|U2GKZ7_9CORY-DECOY | ERIGCLVESPVK | | 10.668,0 |
| 73 | tr\|L1MMA3\|L1MMA3_9CORY-DECOY | ADLSHGPLNTTDK | | 886,114 |
| 74 | tr\|C6R7Z6\|C6R7Z6_9CORY-DECOY | AGMETIMSLNPAR | | 54.924,1 |
| 75 | tr\|U7LSI9\|U7LSI9_9CORY-DECOY | WGELEAIIIQAR<br>Query 3 ELEA 6<br>ELEA<br>Sbjct 250 ELEA 253 | immunity-specific protein Beta371 [Corynephage beta] AAA32187.1 | 21.770,6 |
| 76 | tr\|E4W947\|E4W947_RHOE1-DECOY | RLNLMQVEGGIR<br>Query 3 NLMQV 7<br>NL QV<br>Sbjct 366 NLFQV 370 | Chain A, DIPHTHERIA TOXIN [Corynephage beta] 1DDT_A; Diphtheria toxin homolog CRM228 P00589.2 | 780098 |
| 77 | tr\|M4KGV3\|M4KGV3_9CORY-DECOY | TCAPYVGVGLYIR | | 233685 |
| 78 | tr\|Q6NJ70\|Q6NJ70_CORDI-DECOY | LGLNLMQRFGVR | | 47.846,8 |
| 79 | tr\|G7I0V2\|G7I0V2_9CORY-DECOY | QDTVEWLSKYR<br>Query 6 WLSKYR 11<br>W SK R<br>Sbjct 166 W-SKTR 170 | immunity-specific protein Beta286 [Corynephage beta] AAA32186.1 | 238037 |
| 80 | tr\|E4WFZ0\|E4WFZ0_RHOE1-DECOY | LQAIAWSGPDTLR<br>Query 1 LQAI 4<br>LQAI<br>Sbjct 202 LQAI 205 | immunity-specific protein Beta286 [Corynephage beta] AAA32186.1; immunity-specific protein Beta371 [Corynephage beta] AAA32187.1 | 122889 |
| 81 | tr\|S2YAU2\|S2YAU2_9CORY-DECOY | EFEWGYNGAEIK<br>Query 7 NG-AEIK 12<br>NG AEIK<br>Sbjct 2 NGIAEIK 8 | immunity-specific protein Beta241 [Corynephage beta] AAA32184.1; | 14.837,0 |
| 82 | tr\|Q8FNW8\|Q8FNW8_COREF-DECOY | QIECGWVIIADR | | 20.149,3 |
| 83 | tr\|I4JV79\|I4JV79_CORDP-DECOY | MEGSLAQLEVGTIK | | 87.215,9 |
| 84 | tr\|U7KBY5\|U7KBY5_9CORY-DECOY | EDDAAWLKGIDVR<br>Query 8 KGIDV 12<br>KG DV<br>Sbjct 36 KG-DV 39 | immunity-specific protein Beta371 [Corynephage beta] AAA32187.1 | 50.696,1 |
| 85 | tr\|G0CPT6\|G0CPT6_CORUL-DECOY | ITVAGEGKMAIGEGR | | 67.967,9 |
| 86 | tr\|H2G5T8\|H2G5T8_CORD3-DECOY | VSAVLMVVQSHAPR<br>Query 6 MVVQSHAP 13<br>MV AP<br>Sbjct 224 MVEAARAP 231 | immunity-specific protein Beta241 [Corynephage beta] AAA32184.1 | 53.531,1 |
| 87 | tr\|B0RBS7\|B0RBS7_CLAMS-DECOY | SSVADLRTNDFHR<br>Query 1 SSVA 4<br>SSVA<br>Sbjct 188 SSVA 191 | immunity-specific protein Beta371 [Corynephage beta] AAA32187.1 | 60.503,7 |
| 88 | tr\|S4XKG1\|S4XKG1_9CORY-DECOY | ADASIQERNLDINK<br>Query 10 LDINK 14<br>LD+NK<br>Sbjct 441 LDVNK 445 | Chain A, DIPHTHERIA TOXIN [Corynephage beta] 1DDT_A | 32.411,3 |
| 89 | tr\|G0CXV1\|G0CXV1_CORUB-DECOY | QDGKSTAGPEAADVGAR | diphtheria toxin, partial [Corynephage beta] ABU25232.1 | 35.802,8 |
| 90 | tr\|Q6M3F5\|Q6M3F5_CORGL-DECOY | ALNEVELVGGHLNNR | | 104022 |
| 91 | tr\|H2HYN6\|H2HYN6_CORDW-DECOY | FRDIMTEVASLKSR | immunity-specific protein Beta241 [Corynephage beta] AAA32184.1 | 35.633,2 |
| 92 | tr\|X5DJP3\|X5DJP3_9CORY-DECOY | IMNRGPEELGLALER<br>Query 8 ELGLAL 13<br>ELGL L<br>Sbjct 105 ELGLSL 110 | Chain A, DIPHTHERIA TOXIN [Corynephage beta] 1DDT_A | 76.245,0 |
| 93 | tr\|C0VWY2\|C0VWY2_9CORY-DECOY | DGADIVVDTPDVAEYK<br>Query 8 GP----EAAD-----VGA 16<br>GP E AD GA<br>Sbjct 314 GPVIESETADNLEKTTGA 331 | Chain A, DIPHTHERIA TOXIN [Corynephage beta] 1DDT_A; Diphtheria toxin homolog CRM228 P00589.2 | 35.106,0 |
| 94 | tr\|H2HDN7\|H2HDN7_CORDJ-DECOY | MLTGATVDGCMLMTAK | | 39.646,5 |
| 95 | tr\|C0E5H1\|C0E5H1_9CORY-DECOY | AWGLSMKDLGSNAAPKR | | 53.003,1 |
| 96 | tr\|E0DHG9\|E0DHG9_9CORY-DECOY | GQEKGMRNVRVEIFR<br>Query 3 EKGMRNVRVEIF 14<br>EK MR R +IF<br>Sbjct 75 EKAMRKTR-QIF 85<br><br>Query 3 EKGMRNVRVEIF 14<br>EK MR R +IF<br>Sbjct 75 EKAMRKTR-QIF 85 | immunity-specific protein Beta201 [Corynephage beta] AAA32185.1<br>immunity-specific protein Beta286 [Corynephage beta] AAA32186.1 | 117398 |
| 97 | tr\|U5E253\|U5E253_COREQ-DECOY | MWGVPEDDAKWSNLHK | | 23.636,7 |

| | | | | |
|---|---|---|---|---|
| 98 | tr\|W5Y2I9\|W5Y2I9_9CORY-DECOY | DGICAGRLAQVEEYGGRR | | 10.776,1 |
| 99 | tr\|U7LQA2\|U7LQA2_9CORY-DECOY | LEGAAMEESIPSSMVEVGR<br>Query 12 SSMVE 16<br>SS VE<br>Sbjct 144 SSSVE 148 | Chain A, DIPHTHERIA TOXIN [Corynephage beta] 1DDT_A; immunity-specific protein Beta241 [Corynephage beta] AAA32184.1; Diphtheria toxin homolog CRM228 P00589.2 | 20.134,2 |
| 100 | tr\|C0VRQ3\|C0VRQ3_9CORY-DECOY | QWTTFPEDVEEELLTIR<br>Query 8 DVEEE 12<br>DV+EE<br>Sbjct 38 DVQEE 42 | immunity-specific protein Beta371 [Corynephage beta] AAA32187.1 | 34.199,2 |
| 101 | tr\|G0HBN0\|G0HBN0_CORVD-DECOY | AVDDSEYPHLLIMMVGLAR<br>Query 14 MVGLAR 19<br>MV AR<br>Sbjct 224 MVEAAR 229 | immunity-specific protein Beta241 [Corynephage beta] AAA32184.1 | 29.976,4 |
| 102 | tr\|Q6NHI4\|Q6NHI4_CORDI-DECOY | AELHPDEQIIGAIIACVDER<br>Query 6 DEQIIGAIIA 15<br>DEQ G IA<br>Sbjct 83 DEQ--GQLIA 90 | immunity-specific protein Beta286 [Corynephage beta] AAA32186.1;<br>immunity-specific protein Beta371 [Corynephage beta]AAA32187.1;<br>immunity-specific protein Beta241 [Corynephage beta]AAA32184.1 | 20.764,9 |
| 103 | tr\|E4WGF5\|E4WGF5_RHOE1-DECOY | VMTPCKDDSNKEGTIMIDK | | 40.311,2 |
| 104 | tr\|I4JU32\|I4JU32_CORDP-DECOY | GETVMFPTQPNKDTCPSSSEAVR<br>Query 16 PSSSEAV 22<br>PSS AV<br>Sbjct 52 PSSCKAV 58 | immunity-specific protein Beta286 [Corynephage beta] AAA32186.1; immunity-specific protein Beta371 [Corynephage beta]AAA32187.1; Chain A, DIPHTHERIA TOXIN [Corynephage beta] 1DDT_A | 14.265,2 |
| 105 | C4LG57DECOY | LIEVVK | | 15.284,8 |
| 106 | C4LHZ0DECOY | AAEVQLK<br>Query 2 AEVQLK 7<br>AE +LK<br>Sbjct 21 AEEELK 26 | immunity-specific protein Beta241 [Corynephage beta] AAA32184.1 | 21161,7 |
| 107 | C4LIC6-DECOY | GTGKEALK | | 29088,8 |
| 108 | C4LKL5DECOY | LSSIEKR<br>Query 2 SSIEK 6<br>SS EK<br>Sbjct 494 SSSEK 498 | Chain A, DIPHTHERIA TOXIN [Corynephage beta]1DDT_A | 118232 |
| 109 | C4LKU8DECOY | VEIIVGGAASGR<br>Query 2 EI----IVGGAAS 10<br>EI + GG AS<br>Sbjct 35 EIHSAGVMGGIAS 47<br><br>Query 4 IVGGAA 9<br>IV AA<br>Sbjct 354 IVNFAA 359 | immunity-specific protein Beta241 [Corynephage beta]AAA32184.1;<br>immunity-specific protein Beta371 [Corynephage beta]AAA32187.1 | 9819,4 |
| 110 | C4LL20DECOY | SWLMDRPSK | | |
| 111 | C4LKI5-DECOY | EGTLEADSLIR<br>Query 3 TLEADSLIR 11<br>T E DS IR<br>Sbjct 400 TVE-DSIIR 407 | Chain A, DIPHTHERIA TOXIN [Corynephage beta]1DDT_A | 22098,3 |
| 112 | C4LJF6-DECOY | GWIYGKDLVCK<br>Query 3 IYGKDLV 9<br>I GK+LV<br>Sbjct 128 IWGKELV 134 | immunity-specific protein Beta371 [Corynephage beta]AAA32187.1 | 14149,2 |
| 113 | C4LK42DECOY | ASGSKQVTSLSDSAR<br>Query 12 DSAR 15<br>DSAR<br>Sbjct 142 DSAR 145 | immunity-specific protein Beta241 [Corynephage beta]AAA32184.1 | 9781,68 |
| 114 | C4LGM5-DECOY | DEIVYSITSTSTDLNTR<br>Query 1 DEIV 4<br>DEIV<br>Sbjct 162 DEIV 165 | immunity-specific protein Beta241 [Corynephage beta]AAA32184.1 | 17907,6 |
| 115 | C4LLA2-DECOY | DFGSTLLAMAATNLGYHDHCR<br>DMMGVGK<br>Query 14 LGYH---DH 19<br>LGY+ DH<br>Sbjct 512 LGYQKTVDH 520 | Chain A, DIPHTHERIA TOXIN [Corynephage beta]1DDT_A | 4646,4 |
| 116 | C4LHT1-DECOY | RLIIRKPIHFVCGAWQSGWGLYAGSTSSMNLLDLALIGAAM<br>TK DLSEL<br>Query 2 LIIRKPI-HFVCGAW 15<br>L+IR P F+ W<br>Sbjct 232 LVIR-PTDRFI---W 242<br><br>Query 22 DLSEL 26<br>+LSEL<br>Sbjct 259 ELSEL 263<br><br>Query 2 GST-SSMNLLD 11<br>GS S NL D<br>Sbjct 196 GSSLSCINL-D 205 | immunity-specific protein Beta286 [Corynephage beta]AAA32186.1;<br>Chain A, DIPHTHERIA TOXIN [Corynephage beta]1DDT_A | 3325,9 |
| 117 | C4LJ65-DECOY | VSDADRVSITK | | 15683,4 |
| 118 | C4LHJ8-DECOY | SSRGAELK | | 18353,8 |
| 119 | C4LIF2-DECOY | TDIVETR | | 22487,6 |
| 120 | tr\|E4WIT2\|E4WIT2_RHOE1-DECOY | GGLLDKPLAAG<br>Query 8 LAAG 11<br>LAAG<br>Sbjct 250 LAAG 253 | immunity-specific protein Beta286 [Corynephage beta]AAA32186.1 | 101298 |
| 121 | tr\|A4QE67\|A4QE67_CORGB-DECOY | ELIAADALGLEK<br>Query 1 ELIAA--DA--LGLE 11<br>+LIAA DA L E<br>Sbjct 87 QLIAAAQDAVFLAIE 101 | immunity-specific protein Beta286 [Corynephage beta]AAA32186.1 | 136195 |
| 122 | tr\|C0VVB6\|C0VVB6_9CORY-DECOY | TDAADLKAR<br>Query 4 ADLKA 8<br>AD KA<br>Sbjct 201 ADAKA 205 | immunity-specific protein Beta371 [Corynephage beta]AAA32187.1 | 67628,4 |
| 123 | tr\|C0E6A3\|C0E6A3_9CORY-DECOY | SVDGEGRLK<br>Query 6 GRLK 9<br>GRLK<br>Sbjct 166 GRLK 169 | immunity-specific protein Beta241 [Corynephage beta]AAA32184.1 | 73668,8 |
| 124 | tr\|C6R757\|C6R757_9CORY-DECOY | KAETVVAGSR<br>Query 1 KAETVV 6<br>KA VV<br>Sbjct 76 KAGGVV 81 | Chain A, DIPHTHERIA TOXIN [Corynephage beta]1DDT_A | 53384,4 |
| 125 | tr\|U7KZM3\|U7KZM3_9CORY-DECOY | GDAVSAVAQVK<br>Query 3 AVSAVAQV 10<br>AV VAQV<br>Sbjct 282 AVN-VAQV 288<br><br>Query 3 AVSAVAQV 10<br>AV VAQV<br>Sbjct 314 AVN-VAQV 320 | Chain A, DIPHTHERIA TOXIN [Corynephage beta]1DDT_A; Diphtheria toxin; Short=DT;<br>AltName: Full=NAD(+)--diphthamide ADP-ribosyltransferase P00588.2 | 354692 |
| 126 | tr\|G7TYF7\|G7TYF7_CORPS-DECOY | VAGLDLFYR<br>Query 1 VAGLDL 6<br>V GLDL<br>Sbjct 268 VPGLDL 273 | immunity-specific protein Beta286 [Corynephage beta]AAA32186.1 | 229395 |
| 127 | tr\|C6RBL0\|C6RBL0_9CORY-DECOY | VDLNETITR<br>Query 1 VDLNETI 7<br>VD ETI<br>Sbjct 96 VDNAETI 102 | Chain A, DIPHTHERIA TOXIN [Corynephage beta]1DDT_A<br>immunity-specific protein Beta286 [Corynephage beta]AAA32186.1;<br>Diphtheria toxin homolog CRM228 P00589.2 | 450580 |
| 128 | tr\|C3PJP0\|C3PJP0_CORA7-DECOY | VTEQGALPTR<br>Query 5 GALP 8<br>GALP<br>Sbjct 140 GALP 143 | immunity-specific protein Beta286 [Corynephage beta]AAA32186.1 | 328669 |
| 129 | tr\|H2G5F7\|H2G5F7_CORD3-DECOY | LLGVLILADR<br>Query 7 LADR 10<br>LADR<br>Sbjct 171 LADR 174 | immunity-specific protein Beta241 [Corynephage beta]AAA32184.1;<br>immunity-specific protein Beta371 [Corynephage beta]AAA32187.1 | 64726,3 |
| 130 | tr\|U2GKG6\|U2GKG6_9CORY-DECOY | ILGPYIGSR<br>Query 5 YIGSR 9<br>Y+ SR<br>Sbjct 6 YVVSR 10 | Diphtheria toxin; Short=DT; AltName: Full=NAD(+)--diphthamide ADP-ribosyltransferaseP00588.2 | 53978,6 |
| 131 | tr\|E0DG18\|E0DG18_9CORY-DECOY | SIATTTSGTR<br>Query 6 TSGTR 10<br>TS TR<br>Sbjct 138 TSITR 142 | immunity-specific protein Beta371 [Corynephage beta]AAA32187.1 | 172670 |
| 132 | tr\|Q9EZ58\|Q9EZ58_CORUL-DECOY | IKGYSTVAR<br>Query 6 TVA 8<br>TVA<br>Sbjct 115 TVA 117 | immunity-specific protein Beta286 [Corynephage beta]AAA32186.1 | 124725 |
| 133 | tr\|F1TJK8\|F1TJK8_COREQ-DECOY | LQTITGQLK<br>Query 4 ITGQ 7<br>ITGQ<br>Sbjct 94 ITGQ 97 | immunity-specific protein Beta286 [Corynephage beta]AAA32186.1 | 175965 |
| 134 | tr\|V6VCH5\|V6VCH5_CORUL-DECOY | IGADDENLR<br>Query 2 GADD 5<br>GADD<br>Sbjct 1 GADD 4 | Chain A, DIPHTHERIA TOXIN [Corynephage beta]1DDT_A; Diphtheria toxin; Short=DT; AltName: Full=NAD(+)--diphthamide ADP-ribosyltransferaseP00588.2 | 270185 |
| 135 | tr\|S4XBN0\|S4XBN0_9CORY-DECOY | AVEGALDLVK<br>Query 1 AVEGALDLVK 13<br>A E L+<br>Sbjct 207 ALEETLN 215 | immunity-specific protein Beta241 [Corynephage beta]AAA32184.1 | 138859 |
| 135 | tr\|E4WD30\|E4WD30_RHOE1-DECOY | KDSPVAAAGAAK<br>Query 3 SPVAAAG--AAK 12<br>S VA A AAK<br>Sbjct 188 SSVASAASVAAK 199 | immunity-specific protein Beta371 [Corynephage beta]AAA32187.1 | 318078 |
| 136 | tr\|Q4JXD2\|Q4JXD2_CORJK-DECOY | DKRGQDAFR<br>Query 2 KRGQDA 7<br>KRGQDA<br>Sbjct 172 KRGQDA 177 | **Sequence motif conserved in:** Diphtheria toxin; Short=DT; AltName: Full=NAD(+)--diphthamide ADP-ribosyltransferaseP00588.2; diphtheria toxin, partial [Corynephage beta]BAB03348.1;<br>Diphtheria toxin homolog CRM228 P00589.2; diphtheria toxin (gtg start codon) [Corynephage beta]AAA32182.1; diphtheria toxin, partial [Corynephage beta]ABU25232.1 | 136415 |
| 137 | tr\|M1NPI7\|M1NPI7_9CORY-DECOY | LSPELRNGWR<br>Query 1 LSPEL 5<br>LS EL<br>Sbjct 159 LSVEL 163 | Chain A, DIPHTHERIA TOXIN [Corynephage beta]1DDT_A | 294450 |
| 138 | tr\|D7WC36\|D7WC36_9CORY-DECOY | IPDFLGALLR<br>Query 6 GALL 9<br>GALL<br>Sbjct 15 GALL 18 | diphtheria toxin, partial [Corynephage beta]ABU25232.1 | 52327,3 |
| 139 | tr\|E4W9X2\|E4W9X2_RHOE1-DECOY | RDTAAGASRGR<br>Query 6 GASR 9<br>GASR<br>Sbjct 130 GASR 133 | Chain A, DIPHTHERIA TOXIN [Corynephage beta]1DDT_A<br>immunity-specific protein Beta371 [Corynephage beta]AAA32187.1 | 93057,4 |
| 140 | tr\|G0HEX8\|G0HEX8_CORVD-DECOY | LDTAGADVVIR<br>Query 4 AGAD-VV 9<br>AGAD VV<br>Sbjct 28 AGADDVV 34 | Diphtheria toxin homolog CRM228 P00589.2 | 100781 |
| 141 | tr\|B1VEE5\|B1VEE5_CORU7-DECOY | DELIIDREK<br>Query 2 ELIIDR 7<br>+LI DR<br>Sbjct 154 QLINDR 159 | immunity-specific protein Beta371 [Corynephage beta]AAA32187.1 | 242425 |
| 142 | tr\|U3GYX9\|U3GYX9_9CORY-DECOY | AGMGIAQEDVR<br>Query 3 MGIA 6<br>MGIA<br>Sbjct 314 MGIA 317 | Chain A, DIPHTHERIA TOXIN [Corynephage beta]1DDT_A | 203518 |
| 143 | tr\|E2MY77\|E2MY77_CORAY-DECOY | APEAFPPGKLVR<br>Query 7 PGKL 10<br>PGKL<br>Sbjct 438 PGKL 441 | Chain A, DIPHTHERIA TOXIN [Corynephage beta]1DDT_A | 65854,9 |
| 144 | tr\|E0DC96\|E0DC96_9CORY-DECOY | EDSRMASAAVR<br>Query 3 SRMASAA 9<br>S ASAA<br>Sbjct 188 SSVASAA 194 | immunity-specific protein Beta371 [Corynephage beta]AAA32187.1 | 55650,6 |
| 145 | tr\|U7KP17\|U7KP17_9CORY-DECOY | SALEGPDVLIAR<br>Query 1 SALE 4<br>SALE<br>Sbjct 232 SALE 235 | immunity-specific protein Beta241 [Corynephage beta]AAA32184.1 | 95544 |
| 146 | tr\|S2Z0N6\|S2Z0N6_9CORY-DECOY | SASVLAIQYFR<br>Query 2 ASVL 5<br>ASVL<br>Sbjct 306 ASVL 309 | immunity-specific protein Beta371 [Corynephage beta]AAA32187.1 | 60004,8 |
| 147 | tr\|Q6NJE5\|Q6NJE5_CORDI-DECOY | VDGGMGMTATIGR<br>Query 5 MGMTATIGR 13<br>M MT I R<br>Sbjct 14 MSMT--IPR 20 | immunity-specific protein Beta371 [Corynephage beta]AAA32187.1 | 320418 |
| 148 | tr\|D5UP67\|D5UP67_TSUPD-DECOY | ASGDKLAGEAQAR<br>Query 2 SGDKLAGE 9<br>SG KL GE<br>Sbjct 9 SGQKLFGE 16 | immunity-specific protein Beta201 [Corynephage beta] AAA32185.1; immunity-specific protein Beta241 [Corynephage beta]AAA32184.1; immunity-specific protein Beta371 [Corynephage beta]AAA32187.1 | 116720 |
| 149 | tr\|H2I288\|H2I288_CORDW-DECOY | VTANDDTAMTGAK<br>Query 10 TGA 12<br>TGA<br>Sbjct 329 TGA 331 | diphtheria toxin, partial [Corynephage beta]ABU25232.1 | 27794,6 |
| 150 | tr\|E2MYV3\|E2MYV3_CORAY-DECOY | MALDHCFVTGR<br>Query 2 ALDH 5<br>AL+H<br>Sbjct 254 ALEH 257 | Chain A, DIPHTHERIA TOXIN [Corynephage beta]1DDT_A | 45995,8 |
| 151 | tr\|U5DUN6\|U5DUN6_COREQ-DECOY | MDDVKVGGSSEGR<br>Query 6 VGGS 9<br>VGGS<br>Sbjct 220 VGGS 223 | Diphtheria toxin homolog CRM228 P00589.2 | 37819,4 |
| 152 | tr\|U7KAZ9\|U7KAZ9_9CORY-DECOY | DGDGLSFTAAIGLR<br>Query 8 TAA------IG 12<br>TAA IG<br>Sbjct 301 TAALSILPGIG 311 | Chain A, DIPHTHERIA TOXIN [Corynephage beta]1DDT_A<br>immunity-specific protein [Corynephage beta]AAA32187.1, Diphtheria toxin homolog CRM228 P00589.2 | 53394,9 |
| 153 | tr\|G7HV77\|G7HV77_9CORY-DECOY | MVEEVFAIEVAR<br>Query 5 VF-AIEVA 11<br>VF AIEVA<br>Sbjct 96 VFLAIEVA 103 | immunity-specific protein Beta286 [Corynephage beta]AAA32186.1 | 115837 |
| 154 | tr\|G2EKZ0\|G2EKZ0_CORGT-DECOY | MPGAMVAFIGEAATGRK<br>Query 5 MVA----FIGE------AA 13<br>MVA +GE AA<br>Sbjct 339 MVAQAIPLVGELVDIGFAA 357 | Chain A, DIPHTHERIA TOXIN [Corynephage beta]1DDT_A<br>immunity-specific protein Beta241 [Corynephage beta]AAA32184.1 | 45186,1 |
| 155 | tr\|T9X0R9\|T9X0R9_CORDP-DECOY | FRVDDVFDNVTI<br>Query 6 VFDNVTI 12<br>VFD VTI<br>Sbjct 49 VFDGVTI 55 | immunity-specific protein Beta201 [Corynephage beta] AAA32185.1 | 52705 |
| 156 | tr\|I7HBP0\|I7HBP0_CORUL-DECOY | WNGGDSKTVVIGEK<br>Query 8 TVVIG-EK 14<br>TVV+ EK<br>Sbjct 187 TVVVKLEK 194 | immunity-specific protein Beta241 [Corynephage beta]AAA32184.1 | 123687 |
| 157 | tr\|L1MFI1\|L1MFI1_9CORY-DECOY | LAQDAQAPIDPLPR<br>Query 1 LAQD 4<br>LAQD<br>Sbjct 139 LAQD 142 | immunity-specific protein Beta241 [Corynephage beta]AAA32184.1 | 35988,8 |
| 158 | tr\|S5SX28\|S5SX28_9CORY-DECOY | INETPVMGALELVK<br>Query 9 AL--EL 12 | Chain A, DIPHTHERIA TOXIN [Corynephage beta]1DDT_A | 26072,1 |

| | | | | |
|---|---|---|---|---|
| | | AL EL<br>Sbjct 158 ALSVEL 163 | | |
| 159 | tr\|E4WJ64\|E4WJ64_RHOE1-DECOY | ESAEKRVVDSEWK<br>Query 7 VVDS 10<br>VVDS<br>Sbjct 5 VVDS 8 | Chain A, DIPHTHERIA TOXIN [Corynephage beta]1DDT_A | 88607,2 |
| 160 | tr\|C6R8P5\|C6R8P5_9CORY-DECOY | MLYEERLDEMSR<br>Query 3 YEERLD 8<br>YE LD<br>Sbjct 83 YELALD 88 | immunity-specific protein Beta371 [Corynephage beta]AAA32187.1 | 89989,8 |
| 161 | tr\|I7GRI0\|I7GRI0_CORUL-DECOY | LLAEQVSQKSPELSK<br>Query 11 PELS 14<br>PELS<br>Sbjct 258 PELS 261 | Chain A, DIPHTHERIA TOXIN [Corynephage beta]1DDT_A | 134167 |
| 162 | tr\|Q6NEZ6\|Q6NEZ6_CORDI-DECOY | MEHAALSEIKALNLR<br>Query 1 MEHAALSEIK 10<br>+EH LSE K<br>Sbjct 255 LEHPELSELK 264<br><br>Query 4 AALS 7<br>AALS<br>Sbjct 302 AALS 305<br><br>Query 7 SEIKA 11<br>SE KA<br>Sbjct 239 SEEKA 243 | Chain A, DIPHTHERIA TOXIN [Corynephage beta]1DDT_A | 127916 |
| 163 | tr\|I4AST9\|I4AST9_CORPS-DECOY | GFNELVRPLTALEVSENA<br>Query 4 ELVRPLTA 11<br>EL RP TA<br>Sbjct 263 ELSRPGTA 270<br><br>Query 10 TALE 13<br>TALE<br>Sbjct 253 TALE 256 | immunity-specific protein Beta371 [Corynephage beta]AAA32187.1; Chain A, DIPHTHERIA TOXIN [Corynephage beta]1DDT_A | 38823,9 |
| 164 | tr\|D5UY40\|D5UY40_TSUPD-DECOY | VGVHMTDNVSALVRLVER<br>Query 9 VSAL 12<br>VSAL<br>Sbjct 314 VSAL 317 | immunity-specific protein Beta371 [Corynephage beta]AAA32187.1 | 50023,4 |
| 164 | tr\|D5UY40\|D5UY40_TSUPD-DECOY | VGVHMTDNVSALVRLVER<br>Query 9 VSAL 12<br>VSAL<br>Sbjct 314 VSAL 317 | immunity-specific protein Beta371 [Corynephage beta]AAA32187.1 | 50023,4 |
| 165 | tr\|D8KKN3\|D8KKN3_CORPF-DECOY | LLAVMELAPICVDPFAVELAR<br>Query 17 VELAR 21<br>VE AR<br>Sbjct 225 VEAAR 229 | immunity-specific protein Beta241 [Corynephage beta]AAA32184.1 | 48237,1 |
| 166 | tr\|G4QRX3\|G4QRX3_CORPS-DECOY | LPLASVSLIK<br>Query 1 LPLASV 6<br>LP ASV<br>Sbjct 452 LPIASV 457 | Diphtheria toxin homolog CRM228 P00589.2 | 537282 |
| 167 | tr\|H2GSR9\|H2GSR9_CORDB-DECOY | EAFFLTAAPAQAK<br>Query 7 AAPAQAK 13<br>AA AK<br>Sbjct 193 AASVAAK 199 | immunity-specific protein Beta371 [Corynephage beta]AAA32187.1 | 1,48E+12 |
| 168 | tr\|I7HCK6\|I7HCK6_CORUL-DECOY | SIVMKNTGETIR<br>Query 1 SIVMKNTG 8<br>SI TG<br>Sbjct 139 SITRETTG 146 | immunity-specific protein Beta371 [Corynephage beta]AAA32187.1 | 201847 |
| 169 | tr\|G4QPT9\|G4QPT9_CORPS-DECOY | GGCALLKVPGLEPNAQV<br>Query 7 KV--PGL 11<br>KV PGL<br>Sbjct 82 KVTYPGL 88 | Chain A, DIPHTHERIA TOXIN [Corynephage beta]1DDT_A | 98756,3 |
| 170 | tr\|W5XZ77\|W5XZ77_9CORY-DECOY | LASSRRADEEGLAEVR<br>Query 1 LA--SSRRA-DEEGLAE 14<br>LA S RRA D LAE<br>Sbjct 139 LAQDSARRARD---LAE 152 | immunity-specific protein Beta241 [Corynephage beta]AAA32184.1 | 159087 |
| 171 | tr\|G2EL51\|G2EL51_CORGT-DECOY | DLWGGDLILEFGRLR<br>Query 4 GGDLI 8<br>GG LI<br>Sbjct 152 GGQLI 156 | immunity-specific protein Beta371 [Corynephage beta]AAA32187.1 | 2873890 |
| 172 | tr\|S5T5Y4\|S5T5Y4_9CORY-DECOY | MSIPRSADFPIGALIR<br>Query 11 IGAL 14<br>IGAL<br>Sbjct 18 IGAL 2 | Diphtheria toxin; Short=DT; AltName: Full=NAD(+)--diphthamide ADP-ribosyltransferaseP00588.2 | 344878 |
| 173 | tr\|E2MU26\|E2MU26_CORAY-DECOY | TARAIEEVSELDPQLLR<br>Query 4 AIE--EVSEL 11<br>A E E SEL<br>Sbjct 254 ALEHPELSEL 263<br><br>Query 2 ARAIEEVSEL 11<br>A AI V EL<br>Sbjct 341 AQAIPLVGEL 350 | Chain A, DIPHTHERIA TOXIN [Corynephage beta]1DDT_A | 16577,9 |
| 174 | tr\|U7L2Q8\|U7L2Q8_9CORY-DECOY | AGDAVDDAITKPVNRGILR<br>Query 3 DA----VDD 7<br>DA VD+<br>Sbjct 61 DAAGYSVDN 69<br><br>Query 4 AVDDAIT----K-PV 13<br>A+D +T K PV<br>Sbjct 463 AIDGDVTFCRPKSPV 477 | Chain A, DIPHTHERIA TOXIN [Corynephage beta]1DDT_A | 669745 |
| 175 | tr\|W5XUR4\|W5XUR4_9CORY-DECOY | AKDLFMQLATLLESMGVK<br>Query 10 TLLE 13<br>TLLE<br>Sbjct 125 TLLE 128 | immunity-specific protein Beta286 [Corynephage beta]AAA32186.1 | 22657 |
| 176 | tr\|E9T6Z1\|E9T6Z1_COREQ-DECOY | APTAGGTSTAEHVDVPFGMTADK<br>Query 6 GTSTAE 11<br>GT TA+<br>Sbjct 268 GTATAD 273 | immunity-specific protein Beta371 [Corynephage beta]AAA32187.1 | 19411,5 |
| 177 | tr\|G4QS18\|G4QS18_CORPS-DECOY | QVEAMLPGSALGIHGGVEAGTPQIR<br>Query 6 LPG--SALGI 13<br>LPG S +GI<br>Sbjct 307 LPGIGSVMGI 316 | Chain A, DIPHTHERIA TOXIN [Corynephage beta]1DDT_A | 20035,1 |
| 178 | tr\|C3PJU1\|C3PJU1_CORA7-DECOY | AMTVQTFPNDAQWALLTQVTTCRK<br>Query 19 VTTCR 23<br>VT CR<br>Sbjct 468 VTFCR 472 | Chain A, DIPHTHERIA TOXIN [Corynephage beta]1DDT_A | 29457 |
| 179 | tr\|H2I973\|H2I973_CORDV-DECOY | ILQGVTINGLIPAKHRTAVKGEGALSVNPK<br>Query 24 ALSV 27<br>ALSV<br>Sbjct 158 ALSV 161<br><br>Query 22 EGALSV 27<br>EG SV<br>Sbjct 142 EGSSSV 147<br><br>Query 26 SV---NP 29<br>SV NP<br>Sbjct 66 SVDNENP 72<br><br>Query 16 RTAVKGE 22<br>RT GE<br>Sbjct 407 RTGFQGE 413<br><br>Query 5 VTINGL 10<br>VT GL<br>Sbjct 83 VTYPGL 88 | Chain A, DIPHTHERIA TOXIN [Corynephage beta]1DDT_A | 14725,4 |
| 180 | tr\|U7LKI2\|U7LKI2_9CORY-DECOY | GPDLLGFDDLVFAMVFPDDQTSATARSIDGDCA REFTMF<br>Query 2 PDLLGFD 8<br>P LL F+<br>Sbjct 124 PTLLEFN 130 | immunity-specific protein Beta286 [Corynephage beta]AAA32186.1 | 4302,9311 |
| 181 | tr\|H2GIV1\|H2GIV1_CORDN-DECOY | NAALWSSSDSNVKALGNEMQNAAGTVGPGLAM HNGAEGIIKTFSPIEQYHEFPSSTR<br>Query 7 SSDS 10<br>SSDS<br>Sbjct 505 SSDS 508 | Chain A, DIPHTHERIA TOXIN [Corynephage beta]1DDT_A | 329,91 |
| 182 | tr\|E9SXB3\|E9SXB3_COREQ-DECOY | AYYALTFDMALLVTADLLAEMYAILVMATYAAI GWTSLTHYGATTHSAHAHDGER<br>Query 14 TADLLAEMYAIL-----VM 27<br>TA L IL VM<br>Sbjct 301 TAAL-----SILPGIGSVM 314<br><br>Query 22 SAHA 25<br>SAHA<br>Sbjct 29 SAHA 32<br><br>Query 15 HYGA 18<br>H GA<br>Sbjct 31 HAGA 34 | Chain A, DIPHTHERIA TOXIN [Corynephage beta]1DDT_A; Diphtheria toxin; Short=DT; AltName: Full=NAD(+)--diphthamide ADP-ribosyltransferaseP00588.2 | 103,83 |

**Supplementary Table S2: Putatively *Corynephage*-derived "DECOY"-Peptides identified within corresponding 16kDa section of the *C. simulans*-SDS-PAGE area**

The 16 kDa-SDS-PAGE section of a freshly generated heat-treated *C. simulans* extract was cut out and analyzed by proteomic analyses of in-blot-trypsin-digested proteins by LC-MS/MS. Analyzes were performed using Scaffold Software$^{TM}$ (version 5.2.1), showing data of a Peptide Report. Filters excluded human proteins and other known contaminating proteins. Only peptides declared as "DECOY"-peptides, which reveal no hits in the Universal Protein Resource (UniProt) databank, are shown. Since it was hypothesized that these peptides originate from a corynephage, alignments for the peptides were performed after BLASTP (protein-protein BLAST®)-search towards ***Corynephage beta*** (taxid:10703). Note the high abundance of peptides with similarity to diphtheria toxin, its variants and a homolog.
