## Supplementary material for "Discovery of Natural Bispecific Antibodies: Is Psoriasis Induced by a Toxigenic *Corynebacterium simulans* and Maintained by CIDAMPs as Autoantigens?": Suppl Table S3

### Supplementary Table S3

| Peptide Number | Protein Name | Database Source | Peptide Sequence | Comments | Abundance (Total TIC) | Peptide Start/Stop |
|---|---|---|---|---|---|---|
| 1 | Prostaglandin E2 receptor EP4 subtype OS=Homo sapiens GN=PTGER4 PE=1 SV=1 | P35408 | SGQHCSDSQRTSSAMSGHSRSFISR | Receptor for prostaglandin E2 (PGE2). The activity of this receptor is mediated by G(s) proteins that stimulate adenylate cyclase. Has a relaxing effect on smooth muscle. Knockout studies in mice suggest that this receptor may be involved in the neonatal adaptation of circulatory system, osteoporosis, as well as initiation of skin immune responses. Cytoplasmic staining in most cancer cells, strongest staining in gliomas. | 8865,22 | 359-383 |
| 2 | Vitamin D3 receptor (Fragment) OS=Homo sapiens GN=VDR PE=3 SV=1 | F8VPF8,F8VRJ4,F8VVY8,F8VXQ9,P11473, P11473-2 | ICGVCGDR | The vitamin D3 receptor, which is a member of the nuclear hormone receptor superfamily of ligand-inducible transcription factors. Downstream targets of vitamin D3 receptor are principally involved in mineral metabolism, though this receptor regulates a variety of other metabolic pathways, such as those involved in immune response and cancer. | 454435 | 23-30 |
| 3 | Interleukin-10 receptor subunit beta (Fragment) OS=Homo sapiens GN=IL10RB PE=4 SV=1 | H7C0Z5,Q08334 | NKAGEWSEPVCEQTTHDETVPSWMVAVILMASVFMVCLALLGCFALLWCVYKK | Important for antiviral defense. Shared cell surface receptor required for the activation of five class 2 cytokines: IL10, IL22, IL26, IL28, and IFNL1. The ligand/receptor complex stimulate the activation of the JAK/STAT signaling pathway leading to the expression of IFN-stimulated genes (ISG), which contribute to the antiviral state. | 4333,41 | 136-188 |
| 4 | Dermokine OS=Homo sapiens GN=DMKN PE=1 SV=3 | DMKN_HUMAN,E7EUS0,Q6E0U4-16,Q6E0U4-2,Q6E0U4-3,Q6E0U4-4,Q6E0U4-5,Q6E0U4-6,Q6E0U4-7 | QAEDVIR<br>QAEDVIRHGADAVR<br>VGEAAHALGNTGHEIGR | May act as a soluble regulator of keratinocyte differentiation. Is upregulated in inflammatory diseases, and it was first observed as expressed in the differentiated layers of skin. The most interesting aspect of this gene is the differential use of promoters and terminators to generate isoforms with unique cellular distributions and domain components. Alternatively spliced transcript variants encoding different isoforms have been identified for this gene. Rare cases of squamous cell carcinomas along with a subsets of cells in few cases of urothelial cancers displayed strong staining. | 92318,1<br>26015<br>178913 | 116-122<br>116-129<br>99-115 |
| 5 | HLA class I histocompatibility antigen, Cw-14 alpha chain OS=Homo sapiens GN=HLA-C PE=4 SV=1 | A2BF26,P30505, Q9TNN7 | YTCHVQHEGLPEPLTLRWGPSSQPTIPIVGIVAGLAVLAVLAVLGAVMAVVMCR | HLA-C14:02 proteins, which are known to bind conventional long peptides, also have the potential to bind N-myristoylated short lipopeptides. | 3617,3 | 281-334 |
| 6 | Cytochrome P450 4F22 OS=Homo sapiens GN=CYP4F22 PE=2 SV=1 | Q6NT55 | LSVDRTRK | Involved in epidermal ceramide biosynthesis. Hydroxylates the terminal carbon (omega-hydroxylation) of ultra-long-chain fatty acyls (C28-C36) prior to ceramide synthesis [1]. Contributes to the synthesis of three classes of omega-hydroxy-ultra-long chain fatty acylceramides having sphingosine, 6-hydroxysphingosine and phytosphingosine bases, all major lipid components that underlie the permeability barrier of the stratum corneum. | 31499,9 | 498-505 |
| 7 | Interleukin-12 subunit beta OS=Homo sapiens GN=IL12B PE=1 SV=1 | P29460 | GSSDPQGVTCGAATLSAERVRGDNKEYEYSVECQEDSACPAAEESLPIEVMVDAVHK | Associates with IL23A to form the IL-23 interleukin. IL-23 binds to a heterodimeric receptor complex composed of IL12RB1 and IL23R, activates the Jak-Stat signaling cascade, stimulates memory rather than naive T-cells and promotes production of pro-inflammatory cytokines. IL-23 induces autoimmune inflammation and thus may be responsible for autoimmune inflammatory diseases and may be important for tumorigenesis. | 31,321 | 161-217 |
| 8 | Interleukin-12 subunit alpha OS=Homo sapiens GN=IL12A PE=4 SV=1 | E7ENE1,E9PGR3 | LSMCPARSLLLVATLVLLDHLSLARNLPVATPDPGMFPCLHHSQNLLR | Heterodimerizes with IL12B to form the IL-12 cytokine or with EBI3/IL27B to form the IL-35 cytokine. This cytokine is required for the T-cell-independent induction of interferon (IFN)-gamma, and is important for the differentiation of both Th1 and Th2 cells. | 703962 | 33-80 |
| 9 | Lactoperoxidase OS=Homo sapiens GN=LPO PE=1 SV=2 | F5H386,P220792, PERL_HUMAN | ISNVFTFAFR | Responsible for the inactivation of a wide range of micro-organisms and hence, important component of defense mechanism against infection. May contribute to maintaining an appropriate H2O2 cellular level, therefore protecting cells from H2O2-caused injuries and inflammation. The gene is present in a gene cluster on chromosome 17. | 205772 | 456-465 |
| 10 | SRSF protein kinase 1 OS=Homo sapiens GN=SRPK1 PE=1 SV=1 | Q5R363 | LKVKIADLGNACWHK | Plays a central role in the regulatory network for splicing, controlling the intranuclear distribution of splicing factors in interphase cells and the reorganization of nuclear speckles during mitosis. Can influence additional steps of mRNA maturation, as well as other cellular activities, such as chromatin reorganization in somatic and sperm cells and cell cycle progression. It is essential for CD8 T cell function and host antigen-specific viral immunity. Hodgkins lymphomas and most cases of colorectal, testis and urothelial cancers displayed strong cytoplasmic positivity. | 134435 | 384-398 |
| 11 | Serine palmitoyltransferase 3 OS=Homo sapiens GN=SPTLC3 PE=4 SV=2 | C9JC36,Q9NUV7 | LLRDAVIYGQPRTRR | This in the stratum granulosum of skin enriched protein is a subunit of the serine palmitoyltransferase complex which catalyzes the rate-limiting step in sphingolipid biosynthesis. This subunit metabolizes lauroyl- and myristoyl-CoA and generates C14 and C16-sphingoid bases. Breast, prostate, ovarian, urothelial and pancreatic cancers as well as a few cholangiocarcinomas showed moderate to strong cytoplasmic positivity. | 755683 | 283-297 |
| 12 | Interleukin-8 OS=Homo sapiens GN=CXCL8 PE=3 SV=1 | C9J4T6,P10145,P10145-2 | LAVALLAAFLISAALCEGAVLPR | Chemokine that plays an important role in neutrophil activation and recruitment. It is highly present in lesional psoriasis scales. | 7575,02 | 5-27 |
| 13 | WD repeat-containing protein 76 OS=Homo sapiens GN=WDR76 PE=1 SV=2 | Q9H967 | VVTTCADCNLR | Specifically binds 5-hydroxymethylcytosine (5hmC), suggesting that it acts as a specific reader of 5hmC. Cell type enriched (Skin - Mitotic cells (Skin), Testis - Spermatocytes). Has a role in T-regs - Cell cycle regulation. | 42453,2 | 511-521 |
| 14 | Carnitine O-palmitoyltransferase 1, muscle isoform (Fragment) OS=Homo sapiens GN=CPT1B PE=4 SV=1 | H7C0S1,K7N7A7,Q92523,Q92523-2, Q92523-3,Q92523-4 | NTDVQAAR | CPT1B is the rate-controlling enzyme of the long-chain fatty acid beta-oxidation pathway in muscle mitochondria. This enzyme is required for the net transport of long-chain fatty acyl-CoAs from the cytoplasm into the mitochondria. | 609287 | 17-24 |
| 15 | Keratin, type II cytoskeletal 2 epidermal OS=Homo sapiens GN=KRT2 PE=1 SV=2 | K22E_HUMAN | GFSSGSAVVSGGSR | Probably contributes to terminal cornification. Associated with keratinocyte activation, proliferation and keratinization. Required for maintenance of corneocytes and keratin filaments in suprabasal keratinocytes in the epidermis of the ear. | 40050,8 | 21-34 |
| 16 | Fatty acid-binding protein, heart OS=Homo sapiens GN=FABP3 PE=1 SV=4 | FABPH_HUMAN,S4R371,S4R3A2 | LGVEFDETTADDR<br>NTEISFK<br>SLGVGFATR | Is thought to play a role in the intracellular transport of long-chain fatty acids and their acyl-CoA esters. Is a candidate tumor suppressor gene for human breast cancer. Alternative splicing results in multiple transcript variants. | 45308,8<br>61277<br>143995 | 67-79<br>60-66<br>23-31 |
| 17 | Clusterin OS=Homo sapiens GN=CLU PE=1 SV=1 | CLUS_HUMAN, P10909-2, P10909-4,P10909-5 | ASSIIDELFQDR<br>KYNELLK<br>LFDSDPITVTVPVEVSR<br>TLLSNLEEAK<br>VTTVASHTSDSDVPSGV | Functions as extracellular chaperone that prevents aggregation of non native proteins. Protects cells against apoptosis and against cytolysis by complement. Modulates NF-kappa-B transcriptional activity. Plays a role in the clearance of immune complexes that arise during cell injury. A mitochondrial form suppresses BAX-dependent release of cytochrome c into the cytoplasm and inhibit apoptosis. Cancer enhanced. | 246081<br>32128,1<br>423554<br>170263<br>241740 | 183-194<br>340-346<br>409-425<br>69-78<br>386-408 |
| 18 | Pancreatic secretory granule membrane major glycoprotein GP2 OS=Homo sapiens GN=GP2 PE=2 SV=3 | GP2_HUMAN, P55259-2, P55259-3,P55259-4 | VLDLGPITR | Functions as an intestinal M-cell transcytotic receptor specific for type-I-piliated bacteria that participates in the mucosal immune response toward these bacteria. At the apical membrane of M-cells it binds fimH, a protein of the bacteria type I pilus tip. Internalizes bound bacteria, like E.coli and S.typhimurium, from the lumen of the intestine and delivers them, through M-cells, to the underlying organized lymphoid follicles where they are captured by antigen-presenting dendritic cells to elicit a mucosal immune response. | 301831 | 494-502 |
| 19 | Lactotransferrin OS=Homo sapiens GN=LTF PE=1 SV=6 | E7EQB2,E7ER44, P02788-2, TRFL_HUMAN | YYGYTGAFR | Transferrins are iron binding transport proteins. The protein demonstrates a broad spectrum of properties, including regulation of iron homeostasis, host defense against a broad range of microbial infections, anti-inflammatory activity, regulation of cellular growth and differentiation and protection against cancer development and metastasis. Antimicrobial, antiviral, antifungal and antiparasitic activity has been found for this protein and its peptides. | 100966 | 544-552 |
| 20 | Arginase-1 OS=Homo sapiens GN=ARG1 PE=1 SV=2 | ARGI1_HUMAN, P05089-2,P05089-3 | GGVEEGPTVLR | Key element of the urea cycle converting L-arginine to urea and L-ornithine. Arginine metabolism is a critical regulator of innate and adaptive immune responses. Involved in an antimicrobial effector pathway in polymorphonuclear granulocytes (PMN). | 121154 | 22-32 |
| 21 | Desmoglein-1 OS=Homo sapiens GN=DSG1 PE=1 SV=2 | B7Z845,DSG1_HUMAN | ESSNVVVTER | Component of intercellular desmosome junctions. Involved in the interaction of plaque proteins and intermediate filaments mediating cell-cell adhesion. has been identified as a target of auto-antibodies in the autoimmune skin blistering disease pemphigus foliaceus. Disruption of this gene has also been associated with the skin diseases palmoplantar keratoderma and erythroderma. | 35669,2 | 916-925 |
| 22 | Dermcidin OS=Homo sapiens GN=DCD PE=1 SV=2 | DCD_HUMAN | SSLLEKGLDGAK | Has an antimicrobial activity during early bacterial colonization. A glycosylated form of the N-terminal peptide may be associated with cachexia (muscle wasting) in cancer patients. Cancer enriched (breast cancer). | 819530 | 63-75 |
| 23 | Lipopolysaccharide-binding protein OS=Homo sapiens GN=LBP PE=1 SV=3 | LBP_HUMAN | ITLPDFTGDLR | Acts as an affinity enhancer for CD14, facilitating its association with LPS. Promotes the release of cytokines in response to bacterial lipopolysaccharide. Liver cancer enriched. | 146181 | 57-67 |
| 24 | Cholesteryl ester transfer protein OS=Homo sapiens GN=CETP PE=1 SV=2 | CETP_HUMAN,H3BRJ9,P11597-2 | GVSLFDIINPEIITR | Involved in the transfer of neutral lipids. Regulates the reverse cholesterol transport, by which excess cholesterol is removed from peripheral tissues and returned to the liver for elimination. Candidate cardiovascular disease genes. | 67090,4 | 454-468 |
| 25 | Tyrosine-protein phosphatase non-receptor type 13 OS=Homo sapiens GN=PTPN13 PE=1 SV=2 | PTN13_HUMAN, Q12923-3,Q12923-4 | LNNSKSVASLNRSPER | Tyrosine phosphatase which regulates negatively FAS-induced apoptosis and NGFR-mediated pro-apoptotic signaling | 169269 | 1021-1036 |
| 26 | Fatty acid-binding protein, epidermal OS=Homo sapiens GN=FABP5 PE=1 SV=3 | FABP5_HUMAN | LVDSKGFDEYMKELGVGIALR | Intracellular carrier for long-chain fatty acids and related active lipids, such as endocannabinoids, that regulate the metabolism and actions of the ligands they bind. Delivers retinoic acid to the nuclear receptor peroxisome proliferator-activated receptor delta; which promotes proliferation and survival. Cancer enriched (cervical cancer, head and neck cancer, urothelial cancer). | 479937 | 13-33 |
| 27 | Alpha-enolase OS=Homo sapiens | ENOA_HUMAN, | EIFDSRGNPTVEVDLFTSK | Glycolytic enzyme that catalyzes the conversion of 2-phosphoglycerate to | 78535,6 | 10-28 |

| | | | | | | |
|---|---|---|---|---|---|---|
| | GN=ENO1 PE=1 SV=2 | K7EM90 | | phosphoenolpyruvate. Stimulates immunoglobulin production. Alpha-enolase has also been identified as an autoantigen in Hashimoto encephalopathy. Is a cancer-related gene. | | |
| 28 | Leucine-rich repeat-containing protein 10B OS=Homo sapiens GN=LRRC10B PE=4 SV=2 | A6NIK2 | ILALDFNK | Unknown function. Is upregulated after viral infection. Several cases of prostate cancer showed moderate to strong cytoplasmic staining. A few pancreatic cancers displayed moderate cytoplasmic and membranous staining. | 138665 | 71-78 |
| 29 | Zinc finger protein 717 OS=Homo sapiens GN=ZNF717 PE=4 SV=1 | C9JSV9,C9JVC3, Q9BY31 | SVLTVHHR | Encodes a Kruppel-associated box (KRAB) zinc-finger protein, which belongs to a large group of transcriptional regulators in mammals. These proteins bind nucleic acids and play important roles in various cellular functions, including cell proliferation, differentiation and apoptosis, and in regulating viral replication and transcription. | 44474,1 | 743-750 |
| 30 | Dipeptidyl peptidase 7 OS=Homo sapiens GN=DPP7 PE=1 SV=3 | Q9UHL4,R4GMR2 | REQQPALR | This protein is a post-proline cleaving aminopeptidase expressed in quiescent lymphocytes. The resting lymphocytes are maintained through suppression of apoptosis, a state which is disrupted by inhibition of this novel serine protease. Most cancer cells showed weak to moderate cytoplasmic positivity. Melanomas, carcinoids, prostate and endometrial cancers displayed moderate to strong cytoplasmic staining. | 16870,2 | 478-458 |
| 31 | Beta-1,4-galactosyltransferase 6 OS=Homo sapiens GN=B4GALT6 PE=4 SV=1 | G3XA83,Q6NT09, Q9UBX8 | GEVQFLGRYK | Catalyzes the synthesis of lactosylceramide (LacCer) via the transfer of galactose from UDP-galactose to glucosylceramide (GlcCer). LacCer is the starting point in the biosynthesis of all gangliosides (membrane-bound glycosphingolipids) which play pivotal roles in the CNS including neuronal maturation and axonal and myelin formation. | 53583,9 | 288-297 |
| 32 | Splicing factor, proline- and glutamine-rich (Fragment) OS=Homo sapiens GN=SFPQ PE=1 SV=1 | H0Y9K7,H0Y9U2, P23246,P23246-2 | MGYMDPRERDMRMGGGGAMN MGDPYGSGGQK | DNA- and RNA binding protein, involved in several nuclear processes. Plays a role in the regulation of DNA virus-mediated innate immune response by assembling into the HDP-RNP complex, a complex that serves as a platform for IRF3 phosphorylation and subsequent innate immune response activation through the cGAS-STING pathway. Regulates platinum response in ovarian cancer-modulating SRSF2 activity | 426,211 | 154-184 |
| 33 | Voltage-dependent L-type calcium channel subunit alpha-1D OS=Homo sapiens GN=CACNA1D PE=1 SV=2 | Q01668, Q01668-2, Q01668-3,Q01668-4 | AAQTMSTSAPPPVGSLSQRK R | Voltage-sensitive calcium channels (VSCC) mediate the entry of calcium ions into excitable cells and are also involved in a variety of calcium-dependent processes. Cancer enhanced (prostate cancer). | 229905 | 65-85 |
| 34 | Disintegrin and metalloproteinase domain-containing protein 28 OS=Homo sapiens GN=ADAM28 PE=4 SV=1 | E7EMU1,H0YBG8, H0YBQ8,Q9UKQ2 | CGDNKVCINAECVDIEK | May play a role in the adhesive and proteolytic events that occur during lymphocyte emigration or may function in ectodomain shedding of lymphocyte surface target proteins, such as FASL and CD40L. | 1313,01 | 353-369 |
| 35 | Tetratricopeptide repeat protein 39B OS=Homo sapiens GN=TTC39B PE=2 SV=4 | Q5VTQ0,Q5VTQ0-2,Q5VTQ0-3,Q5VTQ0-4,Q5VTQ0-5,Q5VTQ0-6,Q5VTQ0-7 | LLKYDHYLVPFTLFELASLY K | Regulates high density lipoprotein (HDL) cholesterol metabolism by promoting the ubiquitination and degradation of the oxysterols receptors LXR (NR1H2 and NR1H3). | 66593,3 | 617-637 |
| 36 | Histamine H1 receptor OS=Homo sapiens GN=HRH1 PE=1 SV=1 | P35367 | WSLGRPLCLFWLSMDYVAST ASIFSVFILCIDRYRSVQQP LRYLK | In peripheral tissues, the H1 subclass of histamine receptors mediates the contraction of smooth muscles, increase in capillary permeability due to contraction of terminal venules, and catecholamine release from adrenal medulla, as well as mediating neurotransmission in the central nervous system. Expressed in Tregs (cell cycle regulation), and memory CD4 T-cells.<br>Hepatocellular carcinomas showed strong cytoplasmic positivity. Moderate positivity was observed in several colorectal and ovarian cancers. | 2163510 | 93-137 |
| 37 | Palmitoyltransferase ZDHHC21 OS=Homo sapiens GN=ZDHHC21 PE=2 SV=1 | Q8IVQ6 | RMDHHCPWINNCVGEDNHWL FLQLCFYTELLTCYALMFSF CHYYYFLPLKK | Palmitoylates FYN, regulates its localization in hair follicles and plays a key role in epidermal homeostasis and hair follicle differentiation. Through the palmitoylation of PLCB1 and the regulation of PLCB1 downstream signaling may indirectly regulate the function of the endothelial barrier and the adhesion of leukocytes to the endothelium. Moderate to strong cytoplasmic staining was displayed in all cancers. | 61,286 | 115-165 |
| 38 | Mineralocorticoid receptor OS=Homo sapiens GN=NR3C2 PE=3 SV=1 | B0ZBF6,B0ZBF8,P08235 , P08235-2, P08235-3,P08235-4 | SSVSSPANINNSRCSVSSPS NTNNRSTLSSPAASTVGSIC SPVNNAFSYTASGTSAGSST LRDVVPSPDTQEK | Receptor for both mineralocorticoids (MC) such as aldosterone and glucocorticoids (GC) such as corticosterone or cortisol. Binds to mineralocorticoid response elements (MRE) and transactivates target genes. The effect of MC is to increase ion and water transport and thus raise extracellular fluid volume and blood pressure and lower potassium levels. Cancer tissues displayed moderate to strong cytoplasmic positivity with a granular pattern. | 24,576 | 295-367 |
| 39 | Nuclear receptor coactivator 2 OS=Homo sapiens GN=NCOA2 PE=1 SV=2 | Q15596 | LIAMKTEK | The protein functions as a transcriptional coactivator for nuclear hormone receptors, including steroid, thyroid, retinoid, and vitamin D receptors. It controls immune tolerance by promoting induced $T_{reg}$ differentiation. | 256657 | 781-788 |
| 40 | Nucleotide-binding oligomerization domain-containing protein 2 OS=Homo sapiens GN=NOD2 PE=4 SV=3 | E9PLF7 | ASVLLGHSPGKR | Apart from sensing bacterial peptidoglycans it plays a role in sensing single-stranded RNA (ssRNA) from viruses. Interacts with mitochondrial antiviral signaling/MAVS, leading to activation of interferon regulatory factor-3/IRF3 and expression of type I interferon. | 54272,1 | 16-27 |
| 41 | Integrin beta-4 OS=Homo sapiens GN=ITGB4 PE=1 SV=5 | P16144, P16144-2, P16144-3,P16144-4, P16144-5 | CSMGQCVCEPGWTGPSCDCP LSNATCIDSNGGICNGRGHC ECGRCHCHQQSLYTDTICEI NYSAIHPGLCEDLR | Plays a critical structural role in the hemidesmosome of epithelial cells. Is required for the regulation of keratinocyte polarity and motility. Is likely to play a pivotal role in the biology of invasive carcinoma. A few cases of pancreatic, urothelial, skin and coloreactal cancers showed moderate to strong membranous positivity. | 19387,8 | 557-630 |
| 42 | Basic leucine zipper transcriptional factor ATF-like 2 OS=Homo sapiens GN=BATF2 PE=2 SV=1 | B4DV37,Q8N1L9, Q8N1L9-2 | LGSSPDNPSSALGLARLQSR EHKPALSAATWQGLVVDPSP HPLLAFPLLSSAQVHF | AP-1 family transcription factor that controls the differentiation of lineage-specific cells in the immune system. Following infection, participates in the differentiation of CD8(+) thymic conventional dendritic cells in the immune system. Selectively suppresses CCN1 transcription and hence blocks the downstream cell proliferation signals produced by CCN1 and inhibits CCN1-induced anchorage-independent growth and invasion in several cancer types, such as breast cancer, malignant glioma and metastatic melanoma. Possibly acts by interfering with AP-1 binding to CCN1 promoter. Cases of colorectal and testis cancers along with rare cases of urothelial, lung cancers and melanomas showed strong immunoreactivity. | 2377930 | 195-250 |
| 43 | Keratin-associated protein 5-1 OS=Homo sapiens GN=KRTAP5-1 PE=2 SV=1 | Q6L8H4 | GGCGSCGGSKGGCGSCGGSK GGCGSGCGGCGSSCCVPVCC CKPMCCCVPACSCSSCGK | In the hair cortex, hair keratin intermediate filaments are embedded in an interfilamentous matrix, consisting of hair keratin-associated protein (KRTAP), which are essential for the formation of a rigid and resistant hair shaft through their extensive disulfide bond cross-linking with abundant cysteine residues of hair keratins. | 2115980 | 97-154 |
| 44 | Complement component C9 OS=Homo sapiens GN=C9 PE=1 SV=2 | P02748 | QCVPTEPCEDAEDDCGNDFQ CSTGRCIKMRLR | Constituent of the membrane attack complex (MAC) that plays a key role in the innate and adaptive immune response by forming pores in the plasma membrane of target cells. | 4017,86 | 87-118 |
| 45 | Small proline-rich protein 4 OS=Homo sapiens GN=SPRR4 PE=2 SV=1 | Q96PI1 | KQCPPK | Cross-linked envelope protein of keratinocytes. Fraction of tumor cells in squamous cell carcinomas and urothelial cancers exhibited strong cytoplasmic and nuclear positivity. | 18409,1 | 52-57 |
| 46 | Palmitoyltransferase ZDHHC17 OS=Homo sapiens GN=ZDHHC17 PE=1 SV=2 | Q8IUH5 | DGFWTYITQIATCSPWMFWM FLNSVFHFMWVAVLLMCQMY QISCLGITTNERMNAR | Palmitoyltransferase that catalyzes the addition of palmitate onto various protein substrates and is involved in a variety of cellular processes. Has no stringent fatty acid selectivity and in addition to palmitate can also transfer onto target proteins myristate from tetradecanoyl-CoA and stearate from octadecanoyl-CoA. May be involved in the sorting or targeting of critical proteins involved in the initiating events of endocytosis at the plasma membrane. Highly expressed in plasmacytoid DCs. Most malignant tissues showed moderate to strong cytoplasmic positivity. | 38,754 | 515-570 |
| 47 | Probable ATP-dependent RNA helicase DDX41 (Fragment) OS=Homo sapiens GN=DDX41 PE=1 SV=1 | H0Y8L8 | IIPPPIKSFK | Is required during post-transcriptional gene expression. May be involved in pre-mRNA splicing.DDX41 is required for cGAS-STING activation against DNA virus infection. DDX41 germline variants cause donor cell leukemia. | 3596980 | 1-10 |
| 48 | SNW domain-containing protein 1 OS=Homo sapiens GN=SNW1 PE=1 SV=1 | G3V3A4, SNW1_HUMAN | VAAAMPVR | Involved in pre-mRNA splicing as component of the spliceosome. Encodes a coactivator that enhances transcription from some Pol II promoters. It can bind to the ligand-binding domain of the vitamin D receptor and to retinoid receptors to enhance vitamin D-, retinoic acid-, estrogen-, and glucocorticoid-mediated gene expression. May be involved in oncogenesis. | 6353000 | 159-166 |
| 49 | Cancer/testis antigen 47A OS=Homo sapiens GN=CT47A1 PE=1 SV=1 | Q5JQC4 | LSEEATEEPDAEEPATEEPT AQEATAPEEVTKSQPEK | Only cytoplasmic expression in seminiferous ducts in testis. This protein represents a member of the cancer/testis gene family 47. This family, also known as CT47, is comprised of 13 nearly identical loci clustered at Xq24. This locus is the most telomeric of the cluster. | **8014,83** | 220-256 |
| 50 | Serologically defined colon cancer antigen 3 (Fragment) OS=Homo sapiens GN=SDCCAG3 PE=4 SV=1 | H7C331,Q96C92, Q96C92-2, Q96C92-3,Q96C92-4 | RAVKAENHVVKLK | Endosome-associated protein that plays a role in membrane receptor sorting, cytokinesis and ciliogenesis. Involved in the presentation of the tumor necrosis factor (TNF) receptor TNFRSF1A on the cell surface, and hence in the modulation of the TNF-induced apoptosis. Weak to moderate cytoplasmic positivity was observed in most malignancies. | **768084** | 69-81 |
| 51 | Breast cancer type 2 susceptibility protein OS=Homo sapiens GN=BRCA2 PE=1 SV=2 | P51587 | NLVSIETVVPPK | Involved in double-strand break repair and/or homologous recombination. Acts by targeting RAD51 to ssDNA over double-stranded DNA, enabling RAD51 to displace replication protein-A (RPA) from ssDNA and stabilizing RAD51-ssDNA filaments by blocking ATP hydrolysis. Silencing of BRCA2 promotes R-loop accumulation at actively transcribed genes in replicating and non-replicating cells, suggesting that BRCA2 mediates the control of R-loop associated genomic instability, independently of its known role in homologous recombination | 19698,1 | 1603-1614 |
| 52 | DIS3-like exonuclease 2 OS=Homo sapiens GN=DIS3L2 PE=1 SV=4 | Q8IYB7,Q8IYB7-2,Q8IYB7-3 | KYALFSPSDHR | Essential for correct mitosis, and negatively regulates cell proliferation. It degrades aberrant RNAs and has an oncogenic role, in which it promotes liver cancer progression. | 1683820 | 264-274 |
| 53 | Melanoma-associated antigen 6 (Fragment) OS=Homo sapiens | E7ETG4,E7EUF2, P43360 | AEMLGSVVGNWQYFFPVIFS KASDSLQLVFGIELMEVDPI | Activator of ubiquitin ligase activity of RING-type zinc finger-containing E3 ubiquitin-protein ligases that acts as a as repressor of autophagy. May play a role in tumor | **1454190** | 133-198 |

| | | | | | | |
|---|---|---|---|---|---|---|
| | GN=MAGEA6 PE=4 SV=1 | | GHVYIFATCLGLSYDGLLGD NQIMPK | transformation or aspects of tumor progression. In vitro promotes cell viability in melanoma cell lines. | | |
| 54 | Replication factor C subunit 1 (Fragment) OS=Homo sapiens GN=RFC1 PE=1 SV=1 | D6RAD2,E0CX09, P35251,P35251-2 | LPVSSKPGKISR | Could play a role in DNA transcription regulation as well as DNA replication and/or repair.<br>It is biologically active in various malignant tumors, may play an important role in the proliferation, progression, invasion, and metastasis of cancer cells. It could act as an oncogene or tumor suppressor gene based on the cellular and histological characteristics of the tumor. | 36557,4 | 61-72 |
| 55 | Ras-specific guanine nucleotide-releasing factor 2 OS=Homo sapiens GN=RASGRF2 PE=1 SV=1 | D6RAS9,O14827 | LKRSIQKAVLESAPADR | Functions as a calcium-regulated nucleotide exchange factor activating both Ras and RAC1 through the exchange of bound GDP for GTP. Preferentially activates HRAS in vivo. In lung adenocarcinoma is associated with tumor invasion and poor prognosis | 20364 | 817-833 |
| 56 | Armadillo repeat-containing protein 8 OS=Homo sapiens GN=ARMC8 PE=1 SV=3 | A8MTG8,B7Z637, C9J2I1,Q8IUR7, Q8IUR7-2, Q8IUR7-3, Q8IUR7-7,Q8IUR7-8 | IVTGLSESSVKVRLAAVR | Component of the CTLH E3 ubiquitin-protein ligase complex that selectively accepts ubiquitin from UBE2H and mediates ubiquitination and subsequent proteasomal degradation of the transcription factor HBP1. Downregulation of ARMC8 promotes tumorigenesis through activating Wnt/β-catenin pathway and EMT in cutaneous squamous cell carcinomas. | 8219,28 | 379-396 |
| 57 | Proteasome-associated protein ECM29 homolog OS=Homo sapiens GN=KIAA0368 PE=1 SV=1 | J3KN16,Q5VYK3, R4GMY1 | EDPKLLSMAYSAVGK | Adapter/scaffolding protein that binds to the 26S proteasome, motor proteins and other compartment specific proteins. Several hepatocellular carcinomas and renal cancers along with a few cases of prostate, lung, urothelial and pancreatic cancers showed moderate to strong cytoplasmic and occasional membranous positivity. | **191475** | 583-597 |
| 58 | Remodeling and spacing factor 1 (Fragment) OS=Homo sapiens GN=RSF1 PE=1 SV=1 | H0YCN2,Q96T23, Q96T23-2,Q96T23-3 | KPDSPPK | It is an important interphase centromere protein and is overexpressed in many types of cancers and correlated with poor overall survival. | **183314** | 546-552 |
| 59 | TATA element modulatory factor OS=Homo sapiens GN=TMF1 PE=1 SV=2 | F8WF45, P820942, TMF1_HUMAN | LNSMVERQEK | Potential coactivator of the androgen receptor. Mediates STAT3 degradation. newly-defined breast cancer antigen. A majority of cancer tissues displayed weak to moderate granular cytoplasmic positivity. | 230648 | 628-637 |
| 60 | Inhibitor of growth family member 5 OS=Homo sapiens GN=ING5 PE=1 SV=1 | E9PEN0,Q8WYH8, Q8WYH8-2 | CKEYSDDKVQLAMQTYEMVD K | A tumor suppressor protein that inhibits cell growth and induces apoptosis. This protein contains a PHD-type zinc finger. It interacts with tumor suppressor p53 and p300, a component of the histone acetyl transferase complex, suggesting a role in transcriptional regulation. Inhibits cell growth, induces a delay in S-phase progression and enhances Fas-induced apoptosis in an INCA1-dependent manner. | 383754 | 75-95 |
| 61 | Centrosomal protein of 78 kDa OS=Homo sapiens GN=CEP78 PE=1 SV=2 | A8MST6,Q5JTW2, Q5JTW2-2, Q5JTW2-3,Q5JTW2-5 | ALKGCLSISSVLKNLELNGL ILRER | May be required for efficient PLK4 centrosomal localization and PLK4-induced overduplication of centrioles. May play a role in cilium biogenesis. Low expression of CEP78 is associated with poor prognosis of colorectal cancer patients. | 24763,9 | 111-135 |
| 62 | Transformation/transcription domain-associated protein OS=Homo sapiens GN=TRRAP PE=1 SV=3 | F2Z2U4,H0Y4W2, Q9Y4A5-2 | ILEAKTK | Adapter protein, which is found in various multiprotein chromatin complexes with histone acetyltransferase activity (HAT), which gives a specific tag for epigenetic transcription activation.<br>Cancer related gene. | 183397 | 3447-3453 |
| 63 | Zinc finger protein 880 OS=Homo sapiens GN=ZNF880 PE=2 SV=2 | F5H026,M0R0M5, Q6PDB4-2 | NLQSEVK | regulation of DNA-templated transcription. Detected in all cancer. Prognostic marker in renal cancer. | 430843 | 67-73 |
| 64 | Retinitis pigmentosa 1-like 1 protein OS=Homo sapiens GN=RP1L1 PE=1 SV=4 | A6NKC6, RP1L1_HUMAN | KVSPMSPK | Involved in Occult macular dystrophy (OCMD) | 1692150 | 1746-1753 |
| 65 | Ankyrin repeat domain-containing protein 26 OS=Homo sapiens GN=ANKRD26 PE=1 SV=3 | ANR26_HUMAN, E7ESJ3,Q9UPS8-3 | DDLTPLLLAVSGK | Acts as a regulator of adipogenesis. Involved in the regulation of the feeding behavior. ANKRD26- related thrombocytopenia is a rare, autosomal dominant condition caused by ANKRD26 gene mutation | 53670,5 | 178-190 |
| 66 | Lebercilin OS=Homo sapiens GN=LCA5 PE=1 SV=2 | LCA5_HUMAN,S4R3K6 | SSLVSSSPASVR | This protein is thought to be involved in centrosomal or ciliary functions. Mutations in this gene cause Leber congenital amaurosis type V. | 332552 | 36-47 |
| 67 | Receptor tyrosine-protein kinase erbB-2 OS=Homo sapiens GN=ERBB2 PE=4 SV=1 | F5H1T4 | FPDEEGACQPCPINCTHS | Encodes a member of the epidermal growth factor (EGF) receptor family of receptor tyrosine kinases. It is upregulated in Prostate cancer, Breast cancer and Endometrial cancer | **44500,1** | 586-603 |
| 68 | Isoform 2 of Regulator of G-protein signaling 8 OS=Homo sapiens GN=RGS8 | P57771-2 | SLSDHPVGK | Regulates G protein-coupled receptor signaling cascades. Modulates the activity of potassium channels that are activated in response to DRD2 and CHRM2 signaling. Cancer enriched (glioma, testis cancer, thyroid cancer). | **10197,5** | 8-16 |
| 69 | Isoform 4 of Xin actin-binding repeat-containing protein 2 OS=Homo sapiens GN=XIRP2 | A4UGR9-4, A4UGR9-5, A4UGR9-6,A4UGR9-7 | EMKMPEGRK<br>MARYQAAVSRGDCR | Protects actin filaments from depolymerization. A few tumor cells in pancreatic, breast, endometrial and ovarian cancers displayed membranous positivity of varying intensity. | 2255360<br>5801,89 | 620-628<br>223-236 |
| 70 | Ribonuclease 4 OS=Homo sapiens GN=RNASE4 PE=1 SV=3 | RNAS4_HUMAN | FNTFIHEDIWNIR | This RNase has marked specificity towards the 3' side of uridine nucleotides. Ribonuclease 4 is associated with aggressiveness and progression of prostate cancer. | 44847,2 | 70-82 |
| 71 | E3 ubiquitin-protein ligase HECTD1 (Fragment) OS=Homo sapiens GN=HECTD1 PE=1 SV=1 | H0YJP0,Q9ULT8 | LGPDSSVR | E3 ubiquitin-protein ligase which accepts ubiquitin from an E2 ubiquitin-conjugating enzyme in the form of a thioester and then directly transfers the ubiquitin to targeted substrates. Catalyzes ubiquitination and degradation of ZNF622, an assembly factor for the ribosomal 60S subunit, in hematopoietic cells, thereby promoting hematopoietic stem cell renewal. A majority of malignant tissues displayed weak to moderate cytoplasmic immunoreactivity. | 2380820 | 891-898 |
| 72 | Aldo-keto reductase family 1 member C4 OS=Homo sapiens GN=AKR1C4 PE=1 SV=3 | P17516 | SIGVSNFNCR | Metabolizes a broad spectrum of natural and synthetic therapeutic steroid and plays an important role in metabolism of androgens, estrogens, progestereone and conjugated steroids. Cancer enriched (liver cancer). | 73243 | 162-171 |
| 73 | Isoform 3 of Relaxin receptor 1 OS=Homo sapiens GN=RXFP1 | Q9HBX9-3 | GLNSLTKLSRAVK | Receptor for relaxins. Cytoplasmic expression in squamous epithelia, basal layers of respiratory epithelia and several glandular epithelia, including prostate. Most squamous cell carcinomas and cases of urothelial cancers were moderately stained. | 20901,1 | 172-184 |
| 74 | Zinc finger protein ZFAT OS=Homo sapiens GN=ZFAT PE=4 SV=1 | E5RH37,E9PBN4,F8W7 M8,H0YC51,Q9P243, Q9P243-2, Q9P243-3,Q9P243-4 | NCLKRHVIQKHSNILLK | May be involved in transcriptional regulation. Overexpression causes down-regulation of a number of genes involved in the immune response. Some genes are also up-regulated. | 5121,26 | 783-799 |
| 75 | HERV-K_12q14.1 provirus ancestral Gag polyprotein OS=Homo sapiens PE=1 SV=2 | P62683,P62684,P62685, P63126,P63128,P63130, P63145,P87889,Q7LDI9, Q9YNA8 | NGQPLSGNEQR<br>"GAG-polyprotein viral zinc-finger; pfam14787"<br><br>**Chain A, DIPHTHERIA TOXIN [Corynephage beta]**<br><br>Query 1 NGQPLSG 7<br>N PLSG<br>Sbjct 69 NENPLSG 75<br><br>**AAA family ATPase [Corynebacterium simulans]**<br><br>Query 4 PLSGNEQR 11<br>PLSG+E R<br>Sbjct 722 PLSGDETR 729 | The products of the Gag polyproteins of infectious retroviruses perform highly complex orchestrated tasks during the assembly, budding, maturation, and infection stages of the viral replication cycle. During viral assembly, the proteins form membrane associations and self-associations that ultimately result in budding of an immature virion from the infected cell. HERV-K Gag and HIV-1 Gag coassemble and that this appears to correlate with the effect of HERV-K Gag expression on HIV-1 particle release and its infectivity. | 12647,6 | 602-612 |
| 76 | Ret finger protein-like 4A OS=Homo sapiens GN=RFPL4A PE=1 SV=3 | A6NLU0,F8VTS6 | CGYACCLQCLNSLQK | Unknown function. Cell type enhanced (Schwann cells, Sertoli cells, Hofbauer cells, Spermatogonia, Langerhans cells, Early spermatids, Exocrine glandular cells). Malignant tissues displayed moderate to strong cytoplasmic immunoreactivity. | **4758,36** | 26-40 |
| 77 | Nicotinamide phosphoribosyltransferase OS=Homo sapiens GN=NAMPT PE=1 SV=1 | P43490,Q5SYT8 | AVPEGFVIPR | The secreted form behaves both as a cytokine with immunomodulating properties and an adipokine with anti-diabetic properties. Plays a role in the modulation of circadian clock function. In many tumors cytoplasmic positivity was observed in lymphoid cells. | 625814 | 118-127 |
| 78 | MAX gene-associated protein (Fragment) OS=Homo sapiens GN=MGA PE=1 SV=1 | H3BTF4,Q8IWI9, Q8IWI9-3,Q8IWI9-4 | VVLVKGGSKTKHFQR | This protein is a cancer related gene. It functions as a dual-specificity transcription factor, regulating the expression of both MAX-network and T-box family target genes. Functions as a repressor or an activator. Most cancer tissues displayed weak to moderate nucleolar and occasional cytoplasmic immunoreactivity. | **149431** | 1087-1101 |
| 79 | Isoform 3 of Receptor tyrosine-protein kinase erbB-3 OS=Homo sapiens GN=ERBB3 | P21860-3 | NGLKMCEPCGGLCPKAF | This cancer related gene encodes a member of the epidermal growth factor receptor (EGFR) family of receptor tyrosine kinases. This membrane-bound protein has a neuregulin binding domain but not an active kinase domain. It plays an essential role as cell surface receptor for neuregulins. Involved in the regulation of myeloid cell differentiation.<br>Cancer tissues displayed moderate to strong cytoplasmic staining. | **748086** | 315-331 |
| 80 | Immunoglobulin-like and fibronectin type III domain-containing protein 1 OS=Homo sapiens GN=IGFN1 PE=4 SV=1 | B7WP51, Q86VF2-2,Q86VF2-3 | MGWQPMGENWGCLEEMLNED QSR | Unknown function. Cancer enriched (thyroid cancer). Prognostic marker in renal cancer. | 327014 | 1-23 |
| 81 | Nerve growth factor receptor OS=Homo sapiens GN=NGFR PE=2 SV=1 | B4E096 | MSAPCVEADDAVCRCAYGYY QDETTGRCEACR | Cancer-related gene.Low affinity receptor which can bind to NGF, BDNF, NTF3, and NTF4. Plays an important role in differentiation and survival of specific neuronal populations during development. Can mediate cell survival as well as cell death of neural cells. Malignant gliomas displayed strong membranous immunoreactivity in several cases. | **27932** | 1-32 |
| 82 | Rapamycin-insensitive companion of mTOR OS=Homo sapiens GN=RICTOR PE=1 SV=1 | Q6R327,Q6R327-3 | FLASKMGIIATFRSWAGIIN LCKPGNSGIQSLIGVLCIPN MEIR | Subunit of mTORC2, which regulates cell growth and survival in response to hormonal signals. Plays an essential role in embryonic growth and development. A majority of cancer tissues displayed moderate cytoplasmic staining with a granular pattern. Several cases of endometrial, renal, pancreatic and liver cancers were strongly stained. | **42659,4** | 281-324 |
| 83 | Nucleosome-remodeling factor subunit BPTF OS=Homo sapiens GN=BPTF PE=1 SV=3 | E7ETD6,F5GXF5, Q12830, Q12830-2,Q12830-4 | NETENDSKDAEKNREEFEDQ SLEK | The nucleosome remodeling factor (NURF) alters chromatin accessibility through interactions with its largest subunit,the bromodomain PHD finger transcription factor BPTF. BPTF is overexpressed in several cancers | **148551** | 442-465 |

| | | | | | | |
|---|---|---|---|---|---|---|
| 84 | U6 snRNA-associated Sm-like protein LSm4 OS=Homo sapiens GN=LSM4 PE=1 SV=1 | E9PKF6-DECOY, M0QXB0, Q9NYW1-DECOY, Q9Y4Z0,V9GZ56 | LPLSLLK | Plays role in pre-mRNA splicing as component of the U4/U6-U5 tri-snRNP complex that is involved in spliceosome assembly, and as component of the precatalytic spliceosome. Most cancers displayed weak to moderate granular cytoplasmic immunoreactivity. Hepatocellular carcinomas were strongly stained. A majority of skin, urothelial and renal cancers were negative. | 9768390 | 2-8 |
| 85 | Zinc phosphodiesterase ELAC protein 2 (Fragment) OS=Homo sapiens GN=ELAC2 PE=1 SV=2 | E7ES68,G5E9D5,H7C2I4, Q9BQ52,Q9BQ52-2, Q9BQ52-4,V9GYS7 | RGVRDSSLVVAFICK | Associates with mitochondrial DNA complexes at the nucleoids to initiate RNA processing and ribosome assembly. The protein also interacts with activated Smad family member 2 (Smad2) and its nuclear partner forkhead box H1 (also known as FAST-1), and reduced expression can suppress transforming growth factor-beta induced growth arrest. Mutations in this gene result in an increased risk of prostate cancer. | **422193** | 57-71 |
| 86 | Protein tyrosine kinase 2 OS=Homo sapiens GN=PTK2 PE=1 SV=1 | B4DWJ1,E5RHD8,E7ESA6,E9PEI4,H0YB16, H0YBP1,H0YBZ1, J3QT16,Q05397, Q05397-2,Q05397-5,Q05397-6, Q8IYN9,Q8N9D7 | EIEMAQKLLNSDLGELINKMKLAQQYVMTSLQQEYK | A cancer related cytoplasmic non-receptor protein tyrosine kinase which is found concentrated in the focal adhesions that form between cells growing in the presence of extracellular matrix constituents. It plays an essential role in regulating cell migration, adhesion, spreading, reorganization of the actin cytoskeleton, formation and disassembly of focal adhesions and cell protrusions, cell cycle progression, cell proliferation and apoptosis. A majority of cancer cells displayed weak to moderate cytoplasmic immunoreactivity. | **1548,28** | 292-327 |
| 87 | IQ calmodulin-binding motif-containing protein 1 (Fragment) OS=Homo sapiens GN=IQCB1 PE=1 SV=1 | C9J6Z7,C9JXD7 | MMLQNILQINSGDLLRIGRK | A nephrocystin protein that interacts with calmodulin and the retinitis pigmentosa GTPase regulator protein. The encoded protein has a central coiled-coil region and two calmodulin-binding IQ domains. It is localized to the primary cilia of renal epithelial cells and connecting cilia of photoreceptor cells. The protein is thought to play a role in ciliary function. Moderate cytoplasmic staining was displayed in thyroid cancer. | **333149** | 2-21 |
| 88 | Kelch-like protein 25 OS=Homo sapiens GN=KLHL25 PE=1 SV=1 | Q9H0H3 | YDPGANK | Substrate-specific adapter of a BCR (BTB-CUL3-RBX1) E3 ubiquitin ligase complex required for translational homeostasis. A majority of malignant cells displayed weak to moderate cytoplasmic staining. | **888328** | 417-423 |
| 89 | Juxtaposed with another zinc finger protein 1 (Fragment) OS=Homo sapiens GN=JAZF1 PE=4 SV=1 | H0Y403,Q86VZ6-2 | NCVFITDTDPR | Acts as a transcriptional corepressor of orphan nuclear receptor NR2C2. Plays a role in lipid metabolism by suppressing lipogenesis, increasing lipolysis and decreasing lipid accumulation in adipose tissue. Plays a role in glucose homeostasis by improving glucose metabolism and insulin sensitivity. Several cases of malignant glioma and melanoma, breast, prostate and thyroid cancers showed moderate nuclear and sometimes cytoplasmic staining. | **309891** | 20-30 |
| 90 | Iron-sulfur protein NUBPL (Fragment) OS=Homo sapiens GN=NUBPL PE=4 SV=1 | F8VP02 | MMLSGAGSETLKQR | Required for the assembly of the mitochondrial membrane respiratory chain NADH dehydrogenase (Complex I). Several cases of ovarian, prostate, breast, endometrial and liver cancers showed moderate cytoplasmic immunoreactivity. | **89842,6** | 10-23 |
| 91 | C-Jun-amino-terminal kinase-interacting protein 4 (Fragment) OS=Homo sapiens GN=SPAG9 PE=1 SV=1 | D6RHI8,O60271-9 | MSPGCMLLFVFGFVGGAVVINSAILVSLSVLLLVHFSISTGVPALTQNLPR | This protein is member of the cancer testis antigen gene family. It functions as a scaffold protein that structurally organizes mitogen-activated protein kinases and mediates c-Jun-terminal kinase signaling. This protein also binds to kinesin-1 and may be involved in microtubule-based membrane transport. It may play a role in tumor growth and development. Essentially all cancers showed moderate to strong cytoplasmic positivity. | **6463790** | 1-51 |
| 92 | Colorectal cancer-associated protein 1 OS=Homo sapiens GN=COLCA1 PE=2 SV=1 | Q6ZS62.1 | MESCSVAQAGVLTSPFMWRWTGMAGALSALDNTIEDDADDQLPCGEGRPGWVRGELLGSQGVCKDSK | The gene is expressed by immune cells of the colon that are regulated by colorectal cancer-associated variants. | **46,101** | 1-67 |
| 93 | Selenocysteine insertion sequence-binding protein 2-like (Fragment) OS=Homo sapiens GN=SECISBP2L PE=4 SV=1 | H0YKY4 | RPLVKNVATQKETNAAGALRNPDSGTMNHVESSMCAGGVNWSNVTCQATQK | Binds SECIS (Sec insertion sequence) elements present on selenocysteine (Sec) protein mRNAs, but does not promote Sec incorporation into selenoproteins in vitro. Cancer cells displayed moderate to strong cytoplasmic positivity with a granular pattern. | **61,009** | 188-238 |
| 94 | Beta-defensin 115 OS=Homo sapiens GN=DEFB115 PE=2 SV=1 | Q30KQ5 | MLPDHFSPLSGDIKLSVLALVVLVVLAQTAPDGWIRR | Produced only in Epididymis and is secreted in male reproductive system. Hepatocellular and thyroid cancer exhibited moderate to strong cytoplasmic and nuclear positivity. A few cases of breast and ovarian malignancies showed moderate positivity. | **13031,4** | 1-37 |
| 95 | Lon protease homolog 2, peroxisomal OS=Homo sapiens GN=LONP2 PE=1 SV=1 | B7ZKL7,E7EN44, Q86WA8 | YTREAGVRSLDR | ATP-dependent serine protease that mediates the selective degradation of misfolded and unassembled polypeptides in the peroxisomal matrix. Most malignant cells displayed moderate to strong nuclear immunoreactivity. | **333088** | 511-522 |
| 96 | Rac GTPase-activating protein 1 OS=Homo sapiens GN=RACGAP1 PE=1 SV=1 | Q9H0H5 | VVSHPECRDR | Plays key roles in controlling cell growth and differentiation of hematopoietic cells through mechanisms other than regulating Rac GTPase activity. It promotes oncogenic progression of epithelial ovarian cancer. | **724993** | 321-330 |
| 97 | NACHT, LRR and PYD domains-containing protein 13 OS=Homo sapiens GN=NLRP13 PE=2 SV=2 | Q86W25 | SVTPEWVLQDLIIALQGNSKLTHLNFSSNKLGMTVPLILK | Associations of interactions between NLRP3 SNPs and HLA mismatch with acute and extensive chronic graft-versus-host diseases. | 776355 | 764-803 |
| 98 | Regulating synaptic membrane exocytosis protein 1 OS=Homo sapiens GN=RIMS1 PE=1 SV=2 | E7ENC2,H0YBU6,Q86UR5,Q86UR5-12,Q86UR5-13,Q86UR5-2,Q86UR5-3,Q86UR5-4,Q86UR5-5,Q86UR5-6,Q86UR5-7,Q86UR5-8,Q86UR5-9 | LIGRVILNKRTTMPK | May act as scaffold protein that regulates neurotransmitter release at the active zone. Plays a role in dendrite formation by melanocytes. | 1322130 | 596-610 |
| 99 | E3 ubiquitin-protein ligase TRIM9 OS=Homo sapiens GN=TRIM9 PE=1 SV=1 | Q9C026-4,Q9C026-5,TRIM9_HUMAN | ENDPSGFLQISDALIR | May act as a regulator of synaptic vesicle exocytosis. E3 ubiquitin-protein ligase TRIM9. May play a role in regulation of neuronal functions and may also participate in the formation or breakdown of abnormal inclusions in neurodegenerative disorders. | **89604,5** | 376-391 |
| 100 | Dual-specificity testis-specific protein kinase 2 OS=Homo sapiens GN=TESK2 PE=4 SV=1 | F5GWP9,Q96S53, Q96S53-2,Q96S53-3, | QDLMGGK | This gene is predominantly expressed in testis and prostate. The developmental expression pattern of the rat gene in testis suggests an important role for this gene in meitoic stages and/or early stages of spermiogenesis. | **4151310** | 323-329 |
| | NACHT domain-containing protein [Pseudomonas sp. PDM29] | WP_218543972 | | NACHT domain-containing protein [Pseudomonas sp. PDM29. Predicted NTPases implicated in apoptosis and MHC transcription activation (10782090). | | 1547-1553 |
| 101 | Beta-catenin-like protein 1 OS=Homo sapiens GN=CTNNBL1 PE=1 SV=1 | Q8WYA6, Q8WYA6-2, Q8WYA6-3,Q8WYA6-4 | HDMVRR | Component of the PRP19-CDC5L complex that forms an integral part of the spliceosome and is required for activating pre-mRNA splicing. It is a potent inhibitor of HIV-1 integration via association with viral-encoded integrase and its cofactor, lens epithelium-derived growth factor/p75. | 21196,4 | 464-469 |
| 102 | WD repeat-containing protein 60 (Fragment) OS=Homo sapiens GN=WDR60 PE=1 SV=1 | H7C022,Q8WVS4 | EKYSKEK | Plays a major role in retrograde ciliary protein trafficking in cilia and flagella. Members of the WD-repeat family are involved in a variety of cellular processes including cell cycle progression, signal transduction, apoptosis, and gene regulation. | **11686,3** | 15-21 |
| 103 | Glucosamine-6-phosphate isomerase 2 OS=Homo sapiens GN=GNPDA2 PE=1 SV=1 | Q8TDQ7, Q8TDQ7-2, Q8TDQ7-4,Q8TDQ7-5 | LVDPLFSMK | Has a role in fine tuning the metabolic fluctuations of cytosolic UDP-GlcNAc and their effects on hyaluronan synthesis that occur during tissue remodeling. | 795871 | 265-273 |
| 104 | Nebulin-related-anchoring protein OS=Homo sapiens GN=NRAP PE=4 SV=1 | B1ANW7,Q86VF7, Q86VF7-2, Q86VF7-3,Q86VF7-4 | YEGVGMDRR | May be involved in anchoring the terminal actin filaments in the myofibril to the membrane and in transmitting tension from the myofibrils to the extracellular matrix. | 17219,1 | 149-157 |
| 105 | Dynein heavy chain 3, axonemal OS=Homo sapiens GN=DNAH3 PE=2 SV=1 | Q8TD57 | KTMQIGESLPK | Force generating protein of respiratory cilia. Produces force towards the minus ends of microtubules. It is involved in producing force for ciliary beating by using energy from ATP hydrolysis. | 6585,99 | 4030-4040 |
| 106 | HMG domain-containing protein 3 OS=Homo sapiens GN=HMGXB3 PE=4 SV=1 | E9PEK0,G5E9C5, O76074,O76074-2, Q12766,Q12766-2, Q96GL9 | DSSESSSSAPATQFIMLPLPAYSVVENPTSIKLTTTYTRRGHGTCTSPGCSFTYVTR | This protein contains an HMG-box domain found in DNA binding proteins such as transcription factors and chromosomal proteins. | 9,978 | 271-327 |
| 107 | Protein transport protein Sec61 subunit alpha isoform 2 OS=Homo sapiens GN=SEC61A2 PE=3 SV=1 | F8W776,H7C1S8, Q9H9S3,Q9H9S3-3 | DTSMVHELNR | Found to be tightly associated with membrane-bound ribosomes, either directly or through adaptor proteins. It may also be required for the assembly of membrane and secretory proteins. | 4867,3 | 340-349 |
| 108 | SRSF protein kinase 3 (Fragment) OS=Homo sapiens GN=SRPK3 PE=4 SV=1 | H7C1T4 | RAAAAAAVAAGRAR | Phosphorylates the SR splicing factor SRSF1 and the lamin-B receptor (LBR) in vitro. Required for normal muscle development. | 34606,3 | 1-14 |
| 109 | Sorting nexin-2 OS=Homo sapiens GN=SNX2 PE=1 SV=2 | O60749,O60749-2 | MVNKAADAVNKMTIK | Involved in several stages of intracellular trafficking. Interacts with membranes containing phosphatidylinositol 3-phosphate (PtdIns(3P)) or phosphatidylinositol 3,5-bisphosphate (PtdIns(3,5)P2). Promotes KALRN- and RHOG-dependent but retromer-independent membrane remodeling such as lamellipodium formation | 69424,1 | 284-298 |
| 110 | Regulating synaptic membrane exocytosis protein 1 OS=Homo sapiens GN=RIMS1 PE=1 SV=2 | E7ENC2,H0YBU6,Q86UR5,Q86UR5-12,Q86UR5-13,Q86UR5-2,Q86UR5-3,Q86UR5-4,Q86UR5-5,Q86UR5-6,Q86UR5-7,Q86UR5-8,Q86UR5-9 | LIGRVILNKRTTMPK | Rab effector involved in exocytosis. May act as scaffold protein that regulates neurotransmitter release. Plays a role in dendrite formation by melanocytes. | 1322130 | 596-610 |
| 111 | Tenascin OS=Homo sapiens GN=TNC PE=1 SV=1 | E9PC84,F5H7V9,J3QSU6, P24821,P24821-2, P24821-3,P24821-4, P24821-5,P24821-6 | DCKEQRCPSDCHGQGRCVDGQCICHEGFTGLDCGQHSCPSDCNNLGQCVSGR | Ligand for integrins alpha-8/beta-1, alpha-9/beta-1, alpha-V/beta-3 and alpha-V/beta-6. In tumors, stimulates angiogenesis by elongation, migration and sprouting of endothelial cells | 2792,45 | 557-608 |
| 112 | Neuropeptide FF receptor 2 OS=Homo sapiens GN=NPFFR2 PE=1 SV=2 | Q9Y5X5,Q9Y5X5-2, Q9Y5X5-3 | NKHMHTVTNLFILNLAISDLLVGIFCMPITLLDNIIAGWPFGNTMCK | This receptor mediates its action by association with G proteins that activate a phosphatidylinositol-calcium second messenger system. | **9167,33** | 175-221 |
| 113 | Voltage-dependent T-type calcium channel subunit alpha-1H OS=Homo sapiens GN=CACNA1H | O95180,O95180-2 | GSTSPGPK | T-type channels serve pacemaking functions in both central neurons and cardiac nodal cells and support calcium signaling in secretory cells and vascular smooth muscle (Probable). They may also be involved in the modulation of firing patterns of neurons. | **347417** | 626-633 |

| | | | | | | |
|---|---|---|---|---|---|---|
| | PE=1 SV=4 | | | | | |
| 114 | Proteasome-associated protein ECM29 homolog OS=Homo sapiens GN=KIAA0368 PE=1 SV=1 | J3KN16,Q5VYK3, R4GMY1 | EDPKLLSMAYSAVGK | May couple the proteasome to different compartments including endosome, endoplasmic reticulum and centrosome. May play a role in ERAD and other enhanced proteolysis. Promotes proteasome dissociation under oxidative stress. | **191475** | 583-597 |
| 115 | WD repeat-containing protein 13 OS=Homo sapiens GN=WDR13 PE=1 SV=2 | Q9H1Z4,Q9H1Z4-2 | ALGARGHR | Members of this family are involved in a variety of cellular processes, including cell cycle progression, signal transduction, apoptosis, and gene regulation. A similar protein in mouse is thought to be a negative regulator of the pancreatic beta cell proliferation. Mice lacking this gene exhibit increased pancreatic islet mass and higher serum insulin levels, and are mildly obese. | **2141670** | 102-109 |
| 116 | NAD kinase 2, mitochondrial OS=Homo sapiens GN=NADK2 PE=1 SV=2 | Q4G0N4,Q4G0N4-2 | LGGDGGGRR | Mitochondrial $NAD^+$ kinase that phosphorylates $NAD^+$ to yield $NADP^+$. Mitochondrial NADP(H) generation is essential for proline biosynthesis. | **97387,9** | 37-45 |
| 117 | Striatin-interacting protein 1 OS=Homo sapiens GN=STRIP1 PE=1 SV=1 | Q5VSL9,Q5VSL9-2, Q5VSL9-3, Q9ULQ0,Q9ULQ0-2 | ILLAAAPTSKAK | Plays a role in the regulation of cell morphology and cytoskeletal organization. Required in the cortical actin filament dynamics and cell shape. | **3233750** | 527-538 |
| 118 | Dystonin OS=Homo sapiens GN=DST PE=1 SV=1 | E7ERU0,E7ERU2,E9PEB9,E9PHM6,F8W9J4, H0YC65,J3QTQ0, Q03001,Q03001-8 | IPTPQRKSPASK | Cytoskeletal linker protein. Acts as an integrator of intermediate filaments, actin and microtubule cytoskeleton networks. | **43314,2** | 5357-5368 |
| 119 | Dynein heavy chain 14, axonemal (Fragment) OS=Homo sapiens GN=DNAH14 PE=4 SV=1 | H7BXS7,Q0VDD8, Q0VDD8-4 | DRFHMGLSTILEATTLVTEMQEELLILGPQVEQKTK | Force generating protein of respiratory cilia. Produces force towards the minus ends of microtubules. Expression mainly in ciliated cells in fallopian tube and respiratory epithelia. | **184022** | 708-743 |
| 120 | Ankyrin-2 (Fragment) OS=Homo sapiens GN=ANK2 PE=1 SV=1 | B7Z651,E9PHW9, Q01484,Q01484-2, Q01484-5 | HRLATMPPMVEGEGLASRLIEVGPSGAQFLGKLHLPTAPPPLNEGESLVSR | Plays an essential role in the localization and membrane stabilization of ion transporters and ion channels in several cell types. Plays a role in endocytosis and intracellular protein transport. | **1926550** | 991-1041 |
| 121 | Thyroid adenoma-associated protein OS=Homo sapiens GN=THADA PE=1 SV=2 | C9JER1,C9JJB1, Q6YHU6, Q6YHU6-2, Q6YHU6-4 | FPELYPFLLKQLETVANTVDSDMGEPNRHPSMFLLLLVLERLYASPMDGTSSALSMGPFVPFIMR | The protein is likely involved in the death receptor pathway and apoptosis. | **63550,4** | 999-1063 |
| 122 | Olfactory receptor 2F2 OS=Homo sapiens GN=OR2F2 PE=2 SV=1 | O95006 | IISTILKIQSR | Olfactory receptors interact with odorant molecules in the nose, to initiate a neuronal response that triggers the perception of a smell. | **2251280** | 221-231 |
| 123 | Coiled-coil domain-containing protein 171 (Fragment) OS=Homo sapiens GN=CCDC171 PE=4 SV=2 | H0Y701,Q6TFL3,Q6TFL3-3,Q6TFL3-4 | GVIAVLAANRLKILGQSCASLFTWMESFKEGIGMLVCTGEPQDK | Cytoplasmic expression in several different tissue types. Function of this protein is unknown. | **24,942** | 44-87 |
| 124 | Killer cell lectin-like receptor subfamily F member 1 OS=Homo sapiens GN=KLRF1 PE=1 SV=2 | Q9NZS2,Q9NZS2-2 | ENSCAAIKESK | KLRF1, an activating homodimeric C-type lectin-like receptor (CTLR), is expressed on nearly all natural killer (NK) cells and stimulates their cytotoxicity and cytokine release | 112325 | 206-216 |
| 125 | ER membrane protein complex subunit 4 (Fragment) OS=Homo sapiens GN=EMC4 PE=1 SV=1 | H0YKL2 | HCWDIALGPLKQIPMNLFIMYMAGNTISIFPTMMVCMMAWRPIQALMAISANVRK | Part of the endoplasmic reticulum membrane protein complex (EMC) that enables the energy-independent insertion into endoplasmic reticulum membranes of newly synthesized membrane proteins. | **419011** | 1-55 |
| 126 | Synaptotagmin-6 OS=Homo sapiens GN=SYT6 PE=1 SV=3 | Q5T7P8,V9GYB1 | MSGVWGAGGPRCQEALAVLASLCRARPPPLGLDVETCR | May be involved in Ca(2+)-dependent exocytosis of secretory vesicles through Ca(2+) and phospholipid binding to the C2 domain or may serve as Ca(2+) sensors in the process of vesicular trafficking and exocytosis. | **2314310** | 1-38 |
| 127 | HERV-H_22q11.2 provirus ancestral Gag polyprotein OS=Homo sapiens PE=2 SV=3 | Q8N8A4 | MGNLPPSIPPSSPLACVLKKLKPLQLTPDLKPK | Unknown function. | **12145,9** | 1-33 |
| 128 | Leucine-rich repeat-containing protein 37B OS=Homo sapiens GN=LRRC37B PE=4 SV=1 | J3KTP0,J3QLI7,J3QSU1, O75427,Q68EN5,Q68EN5-2 | MAFAECIAPACVMSWLRFWGPWPLLTWQLLSLLVKEAQPLVWVK | Unknown function. Enhanced expression in skin keratinocytes and testis (early and late spermatids). | **19,135** | 1-44 |

**Supplementary Table S3: Low abundance Psoriasis-IgG-bound immunopeptides, identified within the immunoreactive 16 kDa *C. simulans*-WB-Band**

The 16 kDa-WB-band of a freshly generated, heat-treated *C. simulans* extract was cut out and analyzed by proteomic analyses of on-membrane trypsin-digested proteins by LC-MS/MS. Analyzes were performed to identify human proteins using Scaffold Software$^{TM}$ (version 5.2.1) and filters to exclude known contaminating proteins. In the Universal Protein Resource (UniProt) databank identified *Homo sapiens* proteins are shown as Peptide Report. The relative abundance of the peptides is given as “TICs” (Total Ion Chromatograms). Most of the information about the identified proteins shown in the comments, was from The Human Protein Atlas (https://www.proteinatlas.org/). Note the unique similarity of the identified HERV-K epitope with chain A of diphtheria toxin and the AAA family ATP-ase of *C. simulans* (highlighted in yellow).
